# Computational validation of clonal and subclonal copy number alterations from bulk tumour sequencing

**DOI:** 10.1101/2021.02.13.429885

**Authors:** Alice Antonello, Riccardo Bergamin, Nicola Calonaci, Jacob Househam, Salvatore Milite, Marc J Williams, Fabio Anselmi, Alberto d’Onofrio, Vasavi Sundaram, Alona Sosinsky, William CH Cross, Giulio Caravagna

## Abstract

The identification of chromosome number alterations is now widespread in cancer research, but three features of genomic data hinder copy number calling and downstream analyses: the purity of the tumour sample, intra-tumour heterogeneity, and the ploidy of the tumour. To assess these features, consensus methods are often utilised, though these become onerous in projects that involve thousands of genomes. To facilitate the validation of clonal and subclonal copy number variants we present CNAqc, an evolution-inspired toolset that leverages the known quantitative relationships of purity, ploidy and heterogeneity. We validate the algorithms in CNAqc using low-pass single-cell data, as well as extensive simulations. Its application is demonstrated using over 4000 whole genomes and exomes from TCGA, and PCAWG. A real world application of CNAqc in the analysis of clinical tumour samples, has been demonstrated by its incorporation into the validation of clinically accredited bioinformatics pipeline at Genomics England. Our approach is compatible with most bioinformatic pipelines and designed to augment algorithms with automated quality control procedures for data validation.

## Background

Modern cancer genomics studies leverage a combination of tissue bulk sampling and genome sequencing [1–3]. This permits the identification of somatic single nucleotide variants (SNVs), insertions and deletions (indels), copy number alterations (CNAs) [4,5], driver mutations [6,7], mutational signatures [8–11] and intra-tumour heterogeneity as part of clonal deconvolution [12–18]. Whole-genome sequencing (WGS) and whole-exome sequencing (WES) have entered the clinic [19], and the number of public databases of tumour genomes is continuously increasing, which presents distinct challenges to cancer genomic analyses. While SNVs have well-established detection tools [4], CNAs, which are a particularly important aspect of the cancer genome [17,20], are challenging to assess since the baseline ploidy of the tumour (the total chromosome copy number) as well as the percentage of tumour DNA in the assay (i.e., tumour purity), have to be jointly inferred [21–26]. Of particular difficulty is the detection of CNAs occurring in a subset of tumour cells (subclonal CNAs), since this is limited by the current resolution of bulk assays [27] and the available software to infer clonal compositions. While single-cell approaches can identify small sets of cells with shared CNAs, the resolution and quality of that data is still too low to be adopted in the clinic, meaning bulk approaches are favoured even if they can only detect ‘large’ subclones [27]. Unless the issue of calling CNAs in the context of variable tumour purity and intra-tumour heterogeneity are overcome, the efficiency and quality of tumour molecular profiles will continue to be affected.

To address the issue of inaccurate CNA calling we developed CNAqc, the first quantitative framework to integrate somatic mutations, allele-specific CNAs and estimates of tumour purity estimates to quality control (QC) CNA calls generated from WGS and WES assays (Figure 1a). CNAqc maps SNVs and indels to CNA segments, and computes the expected variant allele frequency (VAF) profile based on the particular copy number state called, and tumour purity. Here, the expected VAF of a given mutation varies depending on allele copy state, tumour purity, and clonality, peaking at a theoretical value affected by observational noise [27]. This means that all three genomic features can be assessed simultaneously. We apply CNAqc to several different datasets representing different resolutions and cancer types. These include 2788 WGS samples from the Pan Cancer Analysis of Whole Genomes (PCAWG) cohort [28], with median coverage 45x and somatic data generated by more than six tools plus a consensus algorithm, and 235 WGS samples from the Genomics England 100.000 Genome Project [19], with median coverage 100x and data generated by Illumina’s DRAGEN latest pipeline (v3). Moreover, we tested our tool on 1464 WES samples from The Cancer Genome Atlas (TCGA) cohort [29] and, finally, with 10 WGS samples from two multi-region colorectal cancers at median coverage 80x. Results show that CNAqc can achieve excellent performances with little computational costs. Moreover, the tool is flexible to work with data from many different callers, and we find it capable of improving even over pipelines that develop consensus-calling strategies, often adopted in large cohort studies.

**Figure 1.**
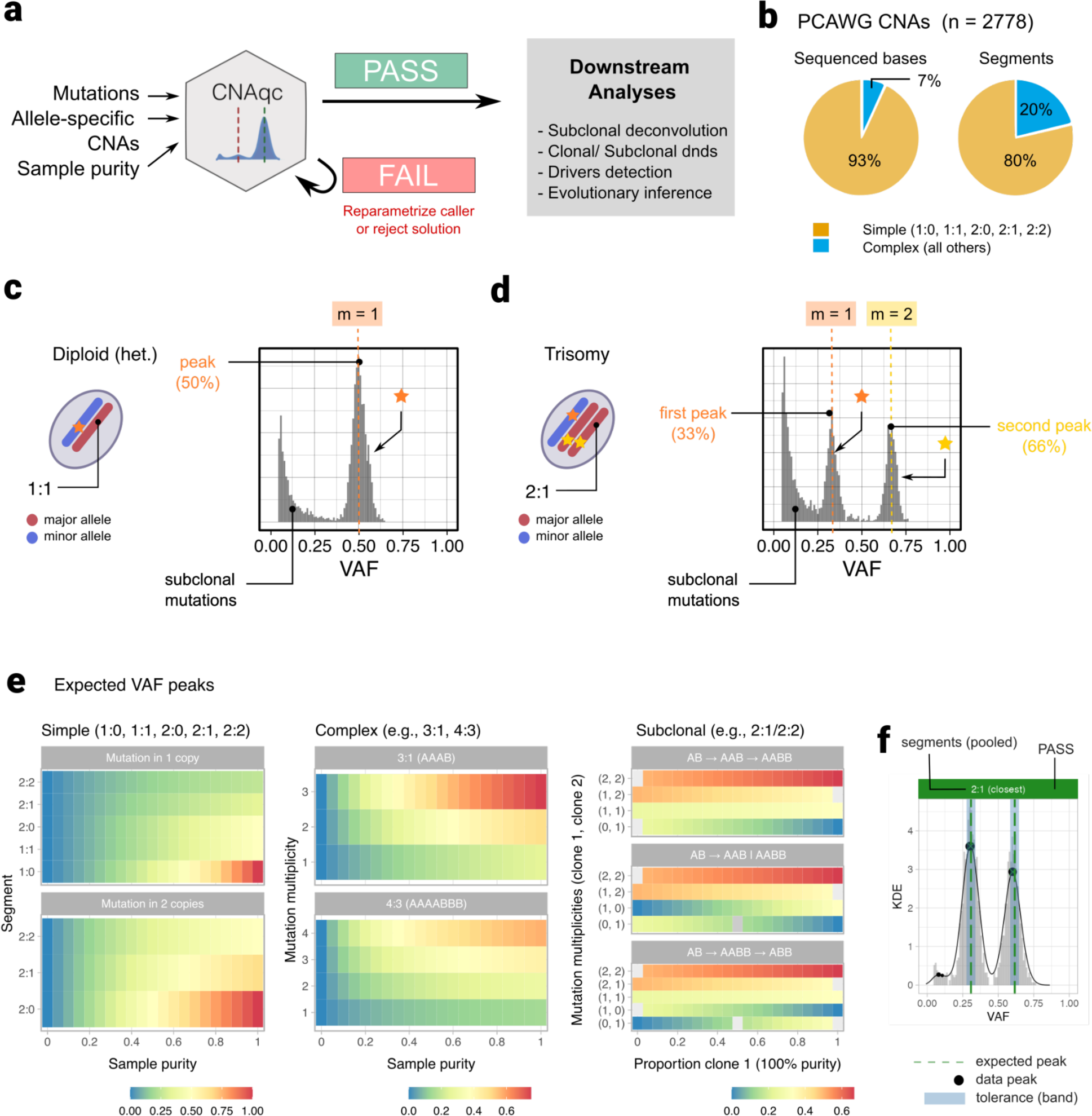
**a.** CNAqc integrates mutations, allele-specific copy number data and tumour purity to computes a score λ ∈ ℜ and quality control (QC) clonal and subclonal tumour aneuploidy. For a sample and a purity error tolerance > 0, the final output is a pass (λ ≤ ɛ) or fail (λ > ɛ) status, which allows propagating good quality calls downstream. The score can be used to correct and re-parametrise the copy number caller, or to select among multiple estimates (e.g., a diploid versus a tetraploid solution). **b.** CNAqc considers clonal simple CNAs, defined as 1:0 (LOH), 1:1 (heterozygous diploid), 2:0 (copy neutral LOH), 2:1 (triploid) and 2:2 (tetraploid) states, the most prevalent CNA in PCAWG both in terms of segments and genome covered. All other CNAs are called complex. **c,d.** VAFs for mutations sitting on 1:1 and 2:1 segments, when π = 1. For 2:1 there are 2 peaks of clonal mutations, at 33% and 66% VAF respectively, corresponding to 100% CCF. Multiplicities determine whether a mutation sits on the amplified segment, or not. **e.** CNAqc equations (1) predicts VAF peaks for any type of CNA (clonal/subclonal, simple/complex), the distance between data and expected peaks is at the core of the QC approach in CNAqc. The colour scale ranges from 0 (blue; low VAF) to 1 (red; high VAF) for the expected peaks. Note that for complex CNAs multiplicities range from 1 to the copies of each allele, and for subclonal CNAs peaks depend on clone sizes, and the evolutionary linear or branching models. **f.** QC for mutations in a 2:1 segment for a ∼90% pure assay. The vertical dashed lines are expected VAF peaks *v*_1_and *v*_2_, from panel (e), the matching bandwidth (shaded area) is obtained by adjusting for the CNA, multiplicity and purity. Black dots are the peaks estimated by CNAqc; since they fall within the bandwidths around *v*_1_and _2_ the final QC result is PASS (green status bar).

## Results

### The CNAqc framework

#### QC of sample purity and copy number segments

CNAqc performs QC of allele-specific CNAs and sample purity estimates prior (Figure 1a). It adopts different algorithms depending on a classification of the CNA, based on the complexity and clonality of the segment. The complexity involves the number of allelic modifications required to generate a certain CNA from a wildtype diploid heterozygous reference, the clonality captures the proportion of cancer cells harbouring a CNA. CNAqc considers simple segments to be: diploid heterozygous (1:1), monosomy (1:0), copy-neutral loss of heterozygosity (CNLOH, 2:0), trisomy (2:1), or tetrasomy (2:2), all of which can be acquired through one copy number aberration event (Figure 1b; Supplementary Figure S1a). These are the most frequently observed CNAs in PCAWG [28] and allow CNAqc to make inferences with precision (simple CNAs in PCAWG: ∼80% of ∼600,000 total segments, covering ∼93% sequenced bases; Supplementary Figure S1b). Simple CNAs are also detectable in bulk sampling datasets, where they also account for the majority of subclonal segments (∼70% in PCAWG; Supplementary Figure S1c-f). Other less frequent CNAs (3:0, 4:1, 5:2 etc.) are more complex to model and QC, as they are acquired via multiple mutation steps, e.g., to achieve a 3:0 one single-copy gain is required on top of a CNLOH.

Bulk sampling makes the true extent of subclonal CNA heterogeneity difficult to estimate, as only major subclones can be detected with current sequencing resolution. CNAqc supports both clonal or subclonal CNAs, and all QC algorithms were designed using the same logic: given a set of major (*n_A_* ≥ 1) and minor (*n_B_* ≥ 0) allele states, a tumour purity of 0 < *π* ≤ 1, and SNVs that overlap the CNAs, the VAF distribution will contain peaks at expected intervals which depend on the percentage of cells harbouring the CNA (Figure 1c,d and Supplementary Figure S2 for two examples). Said differently, CNAqc mathematically links *n_A_* and *n_B_* with the VAF of overlapping SNVs given clonality and tumour content. Deviations from expected VAF peaks indicate errors, which can be quantified and used to suggest adjustment of the input parameters or data. It is possible to QC clonal and simple CNAs using this logic and simple algorithms, though subclonal and complex CNAs require more exhaustive assessments because they depend on the percentage of cells associated to a segment, and the evolutionary steps a CNA has taken. Overall, by combining many segments from the tumour genome, CNAqc determines a pass or fail QC result per sample, which can be used to i) re-parametrise the copy number caller, or to ii) select among alternative copy number profiles returned by an algorithm (e.g., a 100% pure diploid tumour versus a 50% pure tetraploid).

The key CNAqc equation (Figure 1e; Online Methods) predicts a VAF peak for a mutation as: 1 ≤ *m* ≤ *n_A_* alleles out of *n_A_* + *n_B_* total, and is present in a proportion 0 < *c* ≤ 1 of tumour cells (*c* = 1 for a clonal mutation). The expected peak is a function of the mutation multiplicity and; in real data this is clearly observed with noise that, for sequencing, is well captured by Binomial or Beta-Binomial distributions [30].

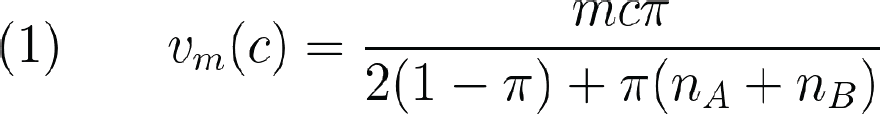

For CNA segments and at least two alleles (*n_A_*> 1 ≥ *n_B_*), the multiplicity phases mutations mapping on amplified and non-amplified segments. For example, for a 2:1 trisomy segment (*n_A_* = 2, *n_B_* = 1), mutations on the amplified allele have *m* = 2 and have been acquired before trisomy. Equation (1) shows that, for a CNA segment, there could be multiple expected VAF peaks as a function of (Figure 1d). As presented below, this equation can be generalised for subclonal CNAs, assuming two subclones and given type of evolutionary relationship. CNAqc is the only method we are aware of that can consider both linear and branching evolutionary models: the two subclones B and C emerge linearly (i.e., nested, A B C) from a unobserved ancestor A, or a branching event (A B → C) of a common ancestor (Figure 1e, Supplementary Figure S3).

CNAqc detects VAF peaks using fast peak-detection algorithms (Supplementary Figure S4) that adopt both nonparametric kernel density estimation and Binomial mixtures to measure *v_m_*(*c*) against data. These algorithms compute an error between data peaks and expected peaks, and require a threshold > 0 (in Euclidean space) on error magnitude to determine if a peak is matched. To make interpretable, however, CNAqc formalises its link to tumour purity and implements a non-linear error propagation to link with (Figure 1f; Online Methods). This allows the user to input in terms of purity units, which are interpretable and intuitive. The overall QC of a bulk sample is determined from all clonal simple CNAs, and a sample score *λ* ∈ ℜ that is a linear combination of the errors accumulated across segments. Because of its interpretability, the score reflects corrections to purity (e.g., +3*%*, −7*%*) that are useful for automatic decision making. For example, for heterozygous diploid mutations and 2.5% purity tolerance (*ɛ* = 0.025) in a sample with *π* = 0.6 (input purity 60*%*, clonal VAF peak 30*%*), CNAqc will accept data peaks in [27.5*%*; 32.5*%*], and a purity estimate in [55*%*; 65*%*].

#### QC of Cancer Cell Fraction (CCF) estimates

In cancer genomics, detecting CNAs along with estimating sample purity and ploidy, is performed as part of Cancer Cell Fraction (CCFs) analyses. Many pipelines that interpret tumour evolutionary trajectories utilise CCFs which, at their core, rely on the estimation of from the data. CNAqc, implements the first algorithm to assess the quality of multiplicity estimates and, in turn, of CCF estimates, from VAFs.

For a mutation with observed VAF, CNAqc defines the CCF (Online methods) by solving equation (1) for [17]

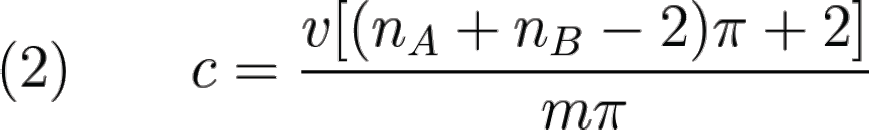

Note that the Binomial noise affecting propagates to, therefore for a diploid clonal mutation the CCF *c*= 2*v* spreads around 1 because spreads around 0.5. After QC, subclonal deconvolution algorithms can denoise *c* by clustering. The real challenge to applying equation (2) is therefore phasing *m* from VAFs. For simple CNAs, this restricts to estimating if *m* = 1 or *m* = 2. CNAqc uses a Binomial mixture to phase *m* from VAFs, and the entropy *H*(*z*) of the mixture latent variables *z* to identify a VAF range where *m* cannot be phased reliably because both *m* = 1 and *m* = 2 seem likely. This information-theoretic approach allows CNAqc to provide a confidence measure over, and therefore. *c*.

As for general CNA/purity-based QC, a final status (pass or fail) can also be determined for CCFs, which help discriminate for which mutations a CCF score cannot be unequivocally computed from VAFs. For this task, other algorithms are also present to aid CCF identification regardless of entropy (Online Methods).

#### Other features of CNAqc

CNAqc provides functions to visualise copy number segments, read counts, CCFs and peak analysis for clonal and subclonal CNAs (Figures 2 and 3). For example, in Figure 3c we show results from QC of complex CNAs for a sample with >3 allele copies, and a large subclonal CNA on chromosome 11, intermixing a 2:1 (21% of cells) and a 2:2 subclone (79% of cells), compatible with both linear and branching evolution. Moreover, the tool contains auxiliary algorithms to smooth segments and detect patterns of over-fragmentation from breakpoints distributions, helping to prioritise additional analysis to determine events of chromothripsis, kataegis or chromoplexy [5,17].

**Figure 2.**
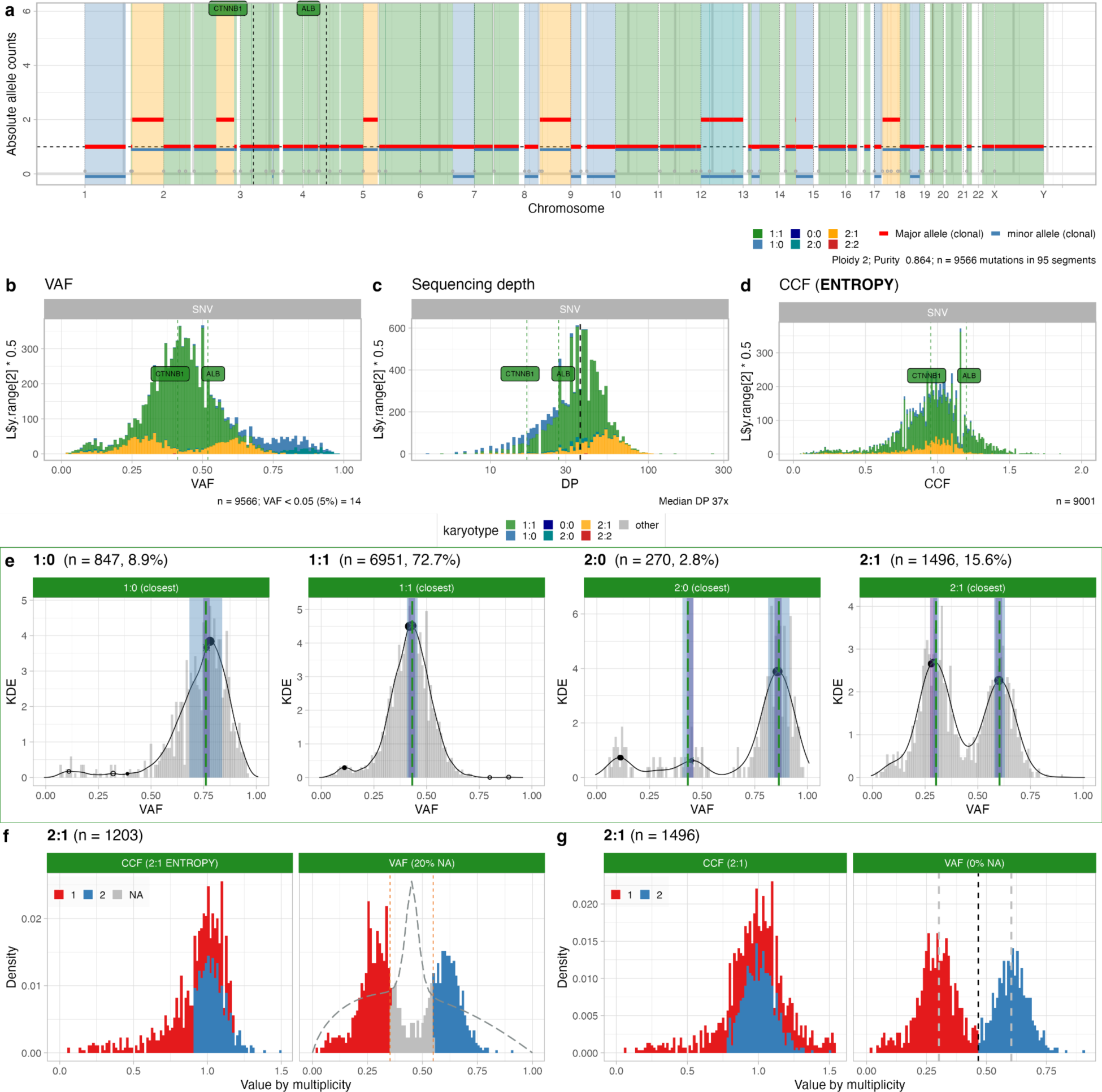
**a.** CNAqc visualisation of PCAWG hepatocellular carcinoma sample: allele-specific consensus CNAs (ploidy 2, purity ∼85%), where the plot shows major and minor allele counts per segment. This sample harbours two driver SNVs hitting genes CTNNB1 and ALB, annotated in diploid heterozygous segments (1:1). **b,c,d.** Read counts for SNVs visualised as variant allele frequencies (VAFs) and depth of sequencing (DP). Cancer Cell Fractions (CCF) obtained by CNAqc show that the two drivers are clonal. **e.** Peak detection QC for simple clonal CNAs. Thick vertical lines are expected peaks, black dots are peaks found in the data (VAF peaks); bars represent areas of peak matching, and are adjusted non-linearly based on mutation multiplicities and CNA configuration. Peaks are checked independently, and the final QC depends on the number of mutations per peak, and whether the peak is matched. The sample-level QC is a linear combination of results from each CNA; here calls are passed (green plot; numbers represent mutational burden). **f,g.** CCF estimation for mutations mapping to triploid 2:1 segments, obtained using the entropy-based and the rough methods. CCF values of clonal mutations spread around 1, CCFs and VAFs are coloured by mutation multiplicity. The entropy profile (dashed line) delineates crossings of Binomial densities where CNAqc detects multiplicity uncertainty; the entropy method detects uncertainty in 20% of the SNVs. The alternative method in panel (g) assigns multiplicities regardless of entropy/uncertainty. In both cases the CCF estimates pass quality control.

**Figure 3.**
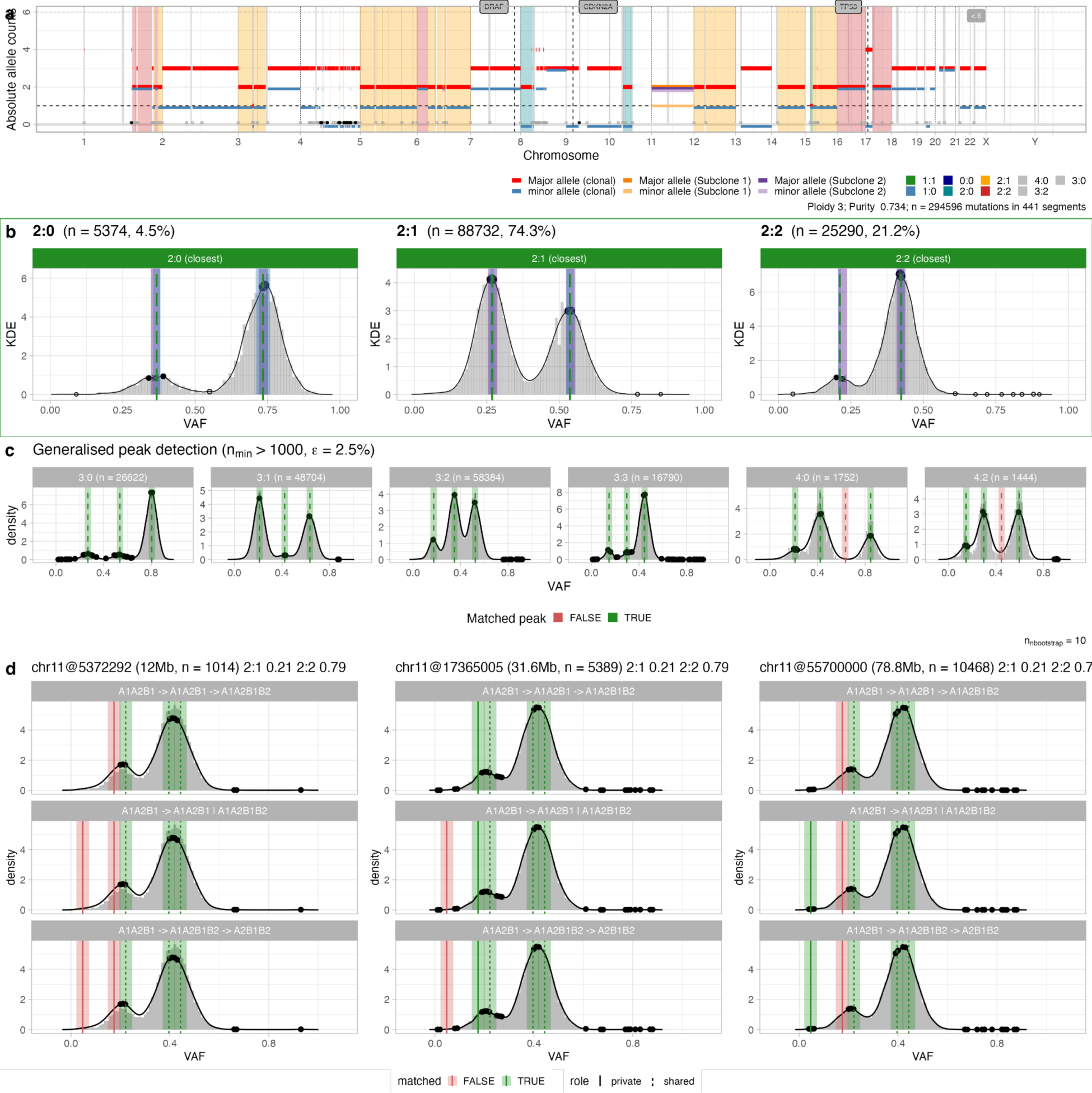
**a** CNAqc visualisation of a skin melanoma PCAWG sample, which presents high aneuploidy (mean ploidy 3.69) and most of the genome is triploid (2:1 segments). The sample has a very large mutational burden (∼294,000 mutations), and large subclonal CNAs on chromosome 11. **b.** CNAqc validates the calls by peak detection; most of the signal is due to ∼80,000 mutations mapping to 2:1 segments. **c.** CNAqc validates 18 out of 20 expected peaks in more complex CNAs, which in this sample are 3:0, 3:1, 3:2, 3:3, 4:0 and 4:2. **d.** CNAqc validates subclonal CNAs on chromosome 11, where two subclones with 2:1 genome (21% of cells) and 2:2 genome (79% of cells) are detected and found to be compatible with both linear and branching models of evolution.

An important feature of CNAqc is its flexibility and speed. The tool has been designed and tested in a variety of settings and pipelines, also against alternative subclonal deconvolution methods which can also detect peaks from VAF data. Tests with hyper-mutant tumours with thousands of mutations (e.g., Figure 3) were used to measure the wall-time performance of our method. Notably, CNAqc was able to load and process ∼500,000 mutations in ∼60 seconds (peaks-based QC) on a standard laptop, which was orders of magnitude faster than alternative methods. For example (Supplementary Figure S5), for our range of tests, variational inference and Monte Carlo methods were from 4/16 times to 100 times slower than CNAqc.

### Simulations, single-cell validation and parameters calibration

CNAqc algorithms and parameters were validated by synthetic simulations, and controlled bioinformatics experiments with single-cell data (Online methods).

#### Validation of error metrics and automatic tool parameterization

We used synthetic data to show that the error metrics implemented in CNAqc work as expected for both purity/CNA and CCF QC algorithms, and to parametrise the algorithm to work best considering coverage and purity of the input assay (sequencing parameters).

From ∼20,000 synthetic tumours with variable coverage (30x to 120x) and known purity (0.4 to 0.95) we ran CNAqc with input purity corrupted by a known error, and observed that the proportion of rejected (fail QC) samples approached 100% when the error exceeded. These tests showed that the performance of CNAqc is affected by sample purity and simulated coverage (Supplementary Figure S6 and S7). Therefore, from >350,000 other synthetic tumours we regressed false positive rates (FPR), i.e. the probability of passing a sample that should fail, against coverage and purity. In this way, CNAqc can suggest, for a desired upper bound on FPR (e.g., maximum 10% of false positives), the best considering the coverage and purity of the input dataset (Supplementary Figure S8).

#### Validation of purity adjustments with single-cell data

We artificially created pseudo-bulks datasets from single-cell datasets with associated low-pass data (Figure 4a), and used that to validate purity-adjustment metrics implemented in CNAqc.

**Figure 4.**
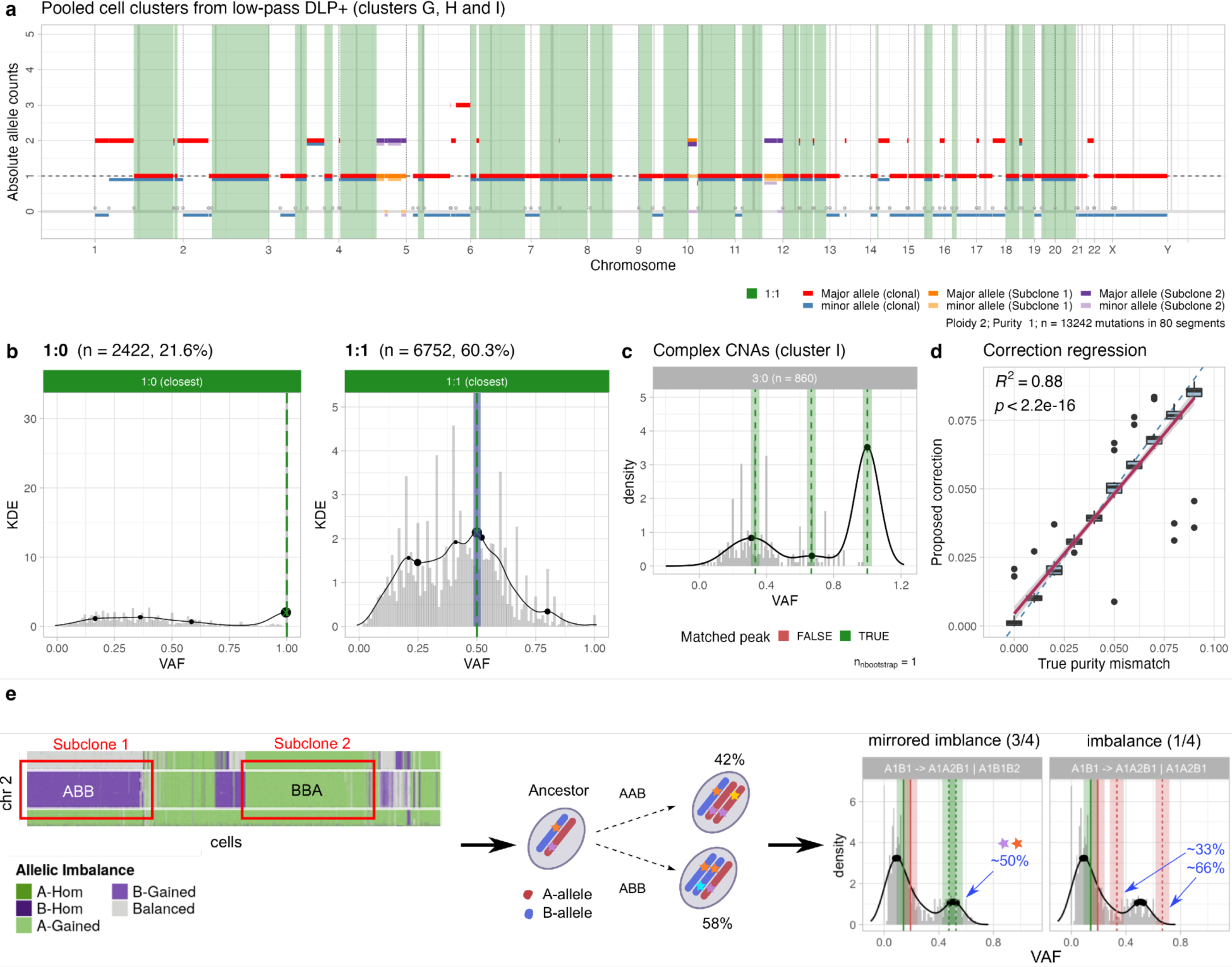
**a.** Pseudo-bulk CNAs from single-cell low-pass data of an ovarian cell line [31]. This profile is obtained from pooling CNAs across several diploid tumour subclones, and used to validate CNAqc (Supplementary Figure S9). **b.** QC of LOH and diploid heterozygous segments from panel (a), using true tumour purity (100%). **c.** Quality control of complex 3:0 segments (cluster I; Supplementary Figure S9). **d.** Correlation between purity-adjustments suggested by CNAqc and errors created artificially for single-cell data. By construction, the desired correction sits on the diagonal; the tool achieves *R*^2^ = 0.88, p-value *p* < 10^−16^. **e.** QC of subclonal CNAs from admixing of tumour subclones with mirrored allelic imbalance [33], i.e. two triploid subclones with mirrored alleles (AAB versus ABB). CNAqc can validate these calls via peak detection, and identify the true branching patterns of evolution (AB AAB | ABB), which is characterised by a peak of shared mutations at around 50% VAF.

We collected mutations and CNAs from single-cell low-pass whole-genome data of ovarian cancer cell lines [31,32]. From 3 tumour clones with distinct CNAs (Supplementary Figure S9), we created a pseudo-bulk dataset and consensus clone-level CNAs (Online Methods), which we validated using CNAqc with true 100% purity (Figure 4b-c, Supplementary Figure S10). We then pooled clonal segments across the 3 subclones to create a larger population, and observed that the purity adjustment proposed by CNAqc with miscalled input purity follows linearly (0.88 ≤ *R*^2^ ≤ 0.99; *p* < 10^−16^; Figure 4d, Supplementary Figure S11).

#### Validation of subclonal CNAs and CCFs

CNAqc validated subclonal CNAs and CCFs in artificial datasets from admixed single-cell data and real data, showing its ability to retrieve the tumour subclonal evolution and implement QC accordingly.

We validated two subclones with trisomy and tetrasomy artificially mixed from low-pass data (Supplementary Figure S12). Then, using 10x data [33] and a pseudo-bulk mixture of 2 subclones with a trisomy and mirrored allelic imbalance (Supplementary Figure S13), we tested the evolution-based QC of complex subclonal CNAs. This test was particularly interesting because the two clones have the same segment ploidy (3), but the joint presence of AAB and ABB genotypes (mirrored allelic imbalance) can only be explained by branching from an AB ancestor (Online Methods). CNAqc validated these subclonal CNAs identifying the expected AB AAB | ABB model for the clones (Figure 4e, Supplementary Figure S14).

Moreover, we computed CCFs from VAFs in pseudo-bulks and, from a cluster of cells with a triploid amplification, CNAqc did flag as uncertain the same mutations for which we could not compute multiplicity from single-cell data (Supplementary Figure S15). Finally, we compared CCFs computed by CNAqc to standard subclonal deconvolution tools. On real data, CCFs by CNAqc were consistent with standard methods (Supplementary Figure S16) but, importantly, the uncertainty metrics from CNAqc did identify spurious subclonal clusters explained by miscalled CCFs, showing the importance of using QC metrics to avoid propagating errors in downstream analyses (Supplementary Figure S17).

### Large-scale WGS pan-cancer PCAWG calls

The PCAWG cohort (*n* = 2778 samples, 40 tumour types) contains WGS samples at median depth 45x and purity ∼65%, comparable to our simulations. This cohort comes with copy number and mutation data generated from 6 state-of-the-art algorithms, as part of a curated consensus [16]. Excluding samples with lack of data or too few mutations (Methods), we ran CNAqc on 2589 samples in less than one hour with a standard computer, confirming the speed of our QC.

Overall PCAWG consensus calls for clonal simple CNAs (n = X segments) were passed in 2339 out of 2589 samples (∼90%) with 3% error purity tolerance (= 0.03), confirming the quality of the consensus copy number data (Figure 5a). As with simulations, the QC pass rate was determined by tumour purity and depth of coverage (Figure 5b). We observed paradigmatic examples with low mutational burden that failed QC (Supplementary Figure S18), or rare cases with excessively high-purity (∼100%) that, upon re-analysis, were better fit with very low tumour content (Supplementary Figure S19). Similarly, we validated cases with very high purity >95% (Supplementary Figure S20). We also examined complex clonal CNA segments with at least 150 mutations (610 samples from esophageal, liver, melanoma, ovarian, pancreatic and breast cancers; Figure 5c). The most prevalent CNAs were 3:1, 3:2 and 3:0 (15%, 13% and 10%), for which CNAqc matched >60% of peaks on average (Online Methods).

**Figure 5.**
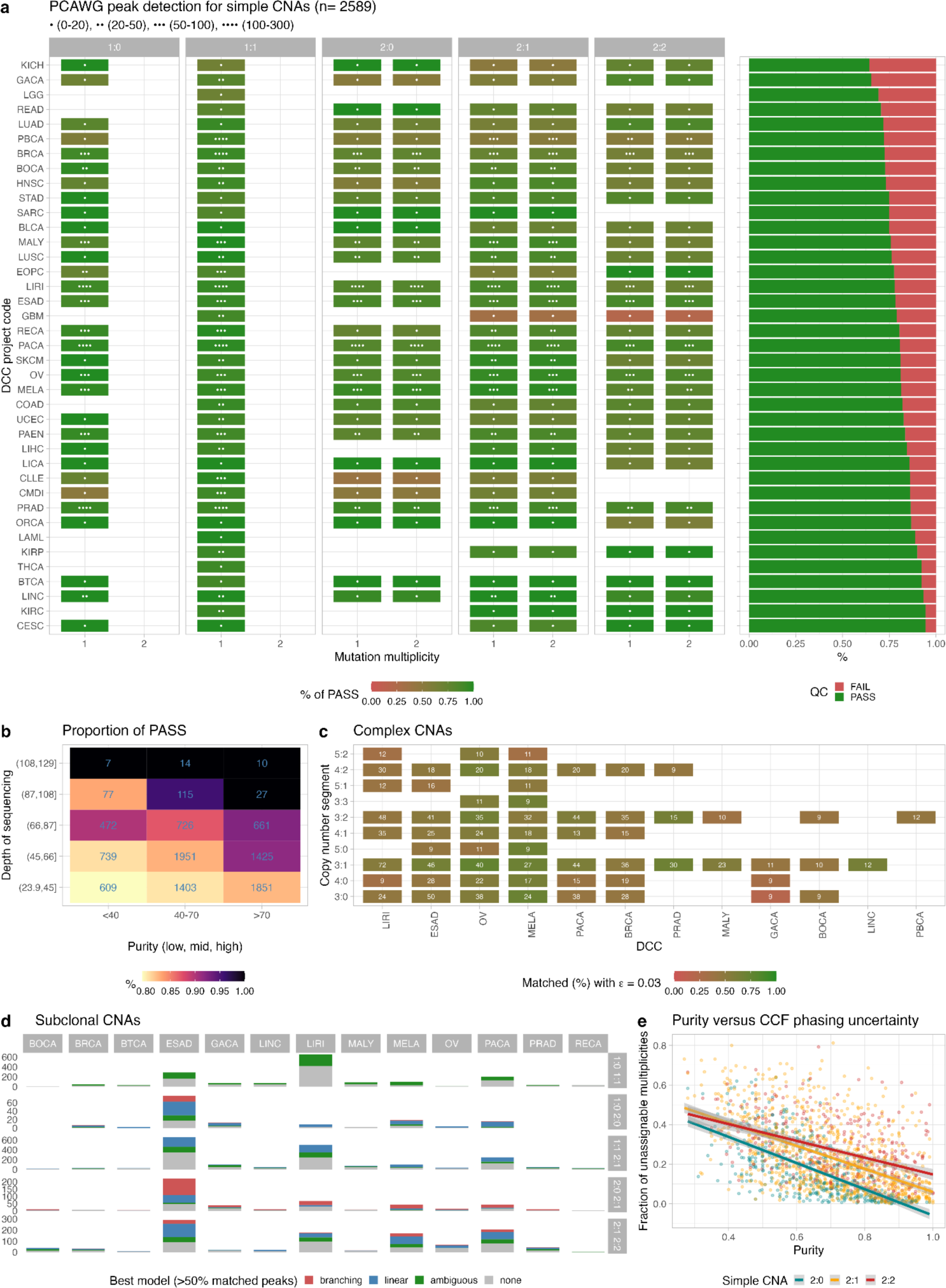
**a.** QC for simple CNAs in *n* = 2589 PCAWG samples (median coverage 45x), using consensus CNA and mutation calls. The plot shows the percentage of cases with pass status, split by CNA type, multiplicity and tumour type, with sample size shown with the asterisks. **b.** Proportion of cases that pass QC, split by purity (low, mid and high) and median depth of sequencing (removing outliers with depth <24X or >129x). **c.** QC for *n* = 570 PCAWG samples with complex CNAs and >150 mutations per segment, with tumour types ranked by number of cases. Numbers report the absolute number of cases. **D.** Best evolution model used to QC *n* = 538 PCAWG samples with subclonal CNAs and >150 mutations per segment, with tumour types ranked by number of cases. **e.** Regression of tumour purity against the proportion of segments with unassigned CCF values using the entropy method in CNAqc.

We applied CNAqc to 538 cases with subclonal CNAs called by Battenberg (Figure 5d) with at least 150 mutations per segment, which were found mostly across esophageal, liver, melanoma, pancreatic and gastric cancers. Interestingly, some of these tumour types also carried complex CNAs, suggesting pervasive chromosomal instability. The most frequent clonal compositions (listed as clone 1 - clone 2) were 1:0 - 1:1 (∼33%), 1:1 - 2:1 (∼31%); 2:1 - 2:2 (19%), 2:0 - 2:1 (∼9%), and 1:0 - 2:0 (∼7% of cases). For each segment, we determined the best fitting model by assessing the percentage of matched peaks, avoiding assigning a model if less than 50% of the expected peaks were matched. Overall, CNAqc assigned a model to ∼87% of the subclonal CNAs (Supplementary Figure S21 and S22). Interestingly, subclones 1:0-2:0, 1:1-2:1 and 2:1-2:2 were generally better explained by linear (A→B→C) evolution implying a temporal ordering among the subclones (in 39%, 48% and 52% of cases, including for subclones 2:1-2:2 in triploid 2:1 or tetraploid 2:2 tumours). This inference can be explained biologically. For example, 1:0-2:0 subclones could emerge as a CNLOH gain (2:0), after a loss (1:0) from a diploid ancestor (1:1). Similarly, 2:1-2:2 subclones can follow a linear amplification path where alleles are progressively gained over time. Conversely, subclones 2:0-2:1 were better explained by branching models (38%), implying the independent formation from a common ancestor. This is intuitive, because, for instance, the evolution from CNLOH to trisomy cannot be linear. In general, these statistics also reflected in tumour types, with 2:0-2:1 subclones in esophageal adenocarcinomas explained by both models, while 2:1-2:2 subclones are better explained by linear models for liver and pancreatic cancers, and melanoma.

Finally, we computed CCFs on the entire PCAWG cohort. As with our simulations, the percentage of unassigned CCFs negatively correlated with sample purity (Figure 5e). The CCFs produced by CNAqc (Supplementary Figure S16) were comparable to those computed by Ccube, the official PCAWG tool to compute CCFs [34]. Comparing CCFs and peak-based analyses we could conclude that, while peaks could be detected for all PCAWG samples, multiplicity phasing would have required higher coverage and purity to reduce uncertainty.

### High-resolution WGS calls at Genomics England

The Cancer Programme of the 100,000 Genomes Project was a transformational UK government project designed to incorporate WGS into NHS clinical service. Genomics England, in partnership with NHS England, generated whole-genome analysis for over 16,000 fresh frozen tumour samples, with a median coverage of 100x. These data provide an ideal retrospective test set for CNAqc, which is now being routinely applied in the validation process of the clinically accredited bioinformatics pipeline at Genomics England.

We gathered a subset of the WGS data (n = 235 samples) with mutation and CNA calls generated by the Illumina DRAGEN (Dynamic Read Analysis for GENomics; >v3.9) platform [35]. These tumours split into groups of distinct subtypes, with the largest groups being paediatric tumours (PT, n = 17), acute lymphoblastic leukaemia (ALL, n = 104), acute myeloid leukaemia (LAML, n = 29), breast cancers (BRCA, n = 41) and sarcomas (SARC, n = 44). This test is of particular importance because these samples are used to optimise the CNA-calling pipelines implemented at Genomics England, which serve both clinical reporting and research.

Results from CNAqc (run using the same parameters as the PCAWG analysis) show a high-quality variant call set with pass rates in all cancer types above 90% (Figure 6a). In comparison, in lower-coverage PCAWG data some tumour types reached only ∼70% pass rate (even if consensus calling was used). For tumour types with large numbers of samples and CNAs the pass rates for Genomics England data with variant calling done by DRAGEN are much higher than the PCAWG consensus (Figure 6b). For instance, for breast tumours (BRCA), we achieved a pass rate of >95% (n = 45 segments) with the Genomics England dataset whereas in PCAWG only ∼75% of segments passed QC (n = 232 segments). Similarly, for sarcoma tumours (SARC) we achieved a pass rate of >90% (n = 47 segments) with the Genomics England dataset whereas in PCAWG only ∼75% (n = 20 segments). In general, with 100x data we could also achieve a high pass rate for subclonal CNAs (Figure 6c), as well as complex clonal CNAs (Figure 6d). In the case of breast cancers we could identify subclonal LOH events validated in 35 out of 36 cases, and the same happened for 32 cases among sarcomas. Overall, all the QC computations reported higher success rates with the Genomics England dataset as compared to PCAWG (example fits in Figure 6g-6h). This trend was confirmed also when we computed CCFs, where we reached ∼15% of unknown estimates when tumour purity in Genomics England samples was ∼50% while in the PCAWG dataset at ∼50% purity ∼40% of the mutations could not be assigned a reliable CCF. The increased coverage in the Genomics England cohort allowed better estimates of tumour CNAs and tumour purity, providing a strong motivation for considering the depth of coverage of a sequencing assay as a key aspect when designing specific analyses.

**Figure 6.**
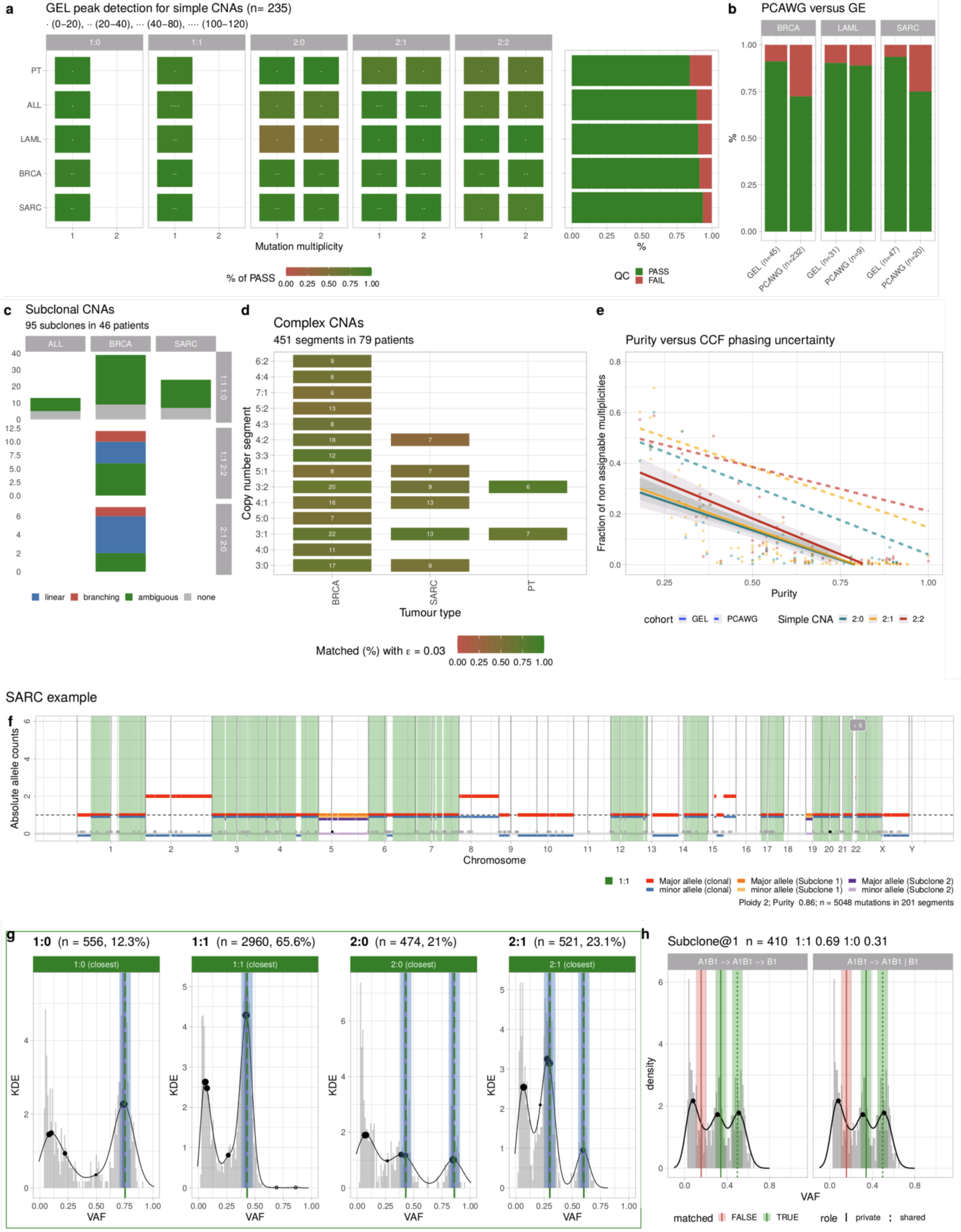
**a.** QC from simple CNAs in *n* = 235 Genomics England samples (median coverage 100x), using DRAGEN data. The plot is like panel (a) in Figure 5 for PCAWG. **b.** Proportion of cases that pass QC, comparing Genomics England and PCAWG, for three tumour types: BRCA (breast cancer), LAML (acute myeloid leukaemia) and SARC (sarcoma). **c-e.** QC for subclonal CNAs in ALL (acute lymphocytic leukaemia), BRCA and SARC (c), complex CNAs in BRCA, SARC and PT (paediatric tumours) (d) and proportion of segments with unassigned CCF values against purity. Plots are as in panels (c-e) in Figure 5 for PCAWG. **f-h.** Example QC of clonal simple (g) and subclonal CNAs (h) for a SARC sample with associated segmentation (f), where DRAGEN detects aneuploidy as well as a subclonal CNA involving an LOH event associated with 410 distinct mutations (in 31% of the tumour cells). These estimates are validated by CNAqc which detects, in the data, VAF peaks.

### Large-scale WES pan-cancer TCGA calls

The statistical signals used by CNAqc are spread through the whole tumour genome, but many assays are limited to sequence, upon capture, only the whole-exome (WES). We used data from TCGA to show i) that the performance of CNAqc is robust also with WES data, and that ii) consensus procedures for purity estimation were imprecise in many TCGA samples.

First, we collected data for *n* = 48 lung TCGA adenocarcinomas [29], a tumour type with high aneuploidy, for which the sample purity and segments are available from popular bioinformatic tools (ESTIMATE [36], LUMP [37] and ABSOLUTE [38]), as well from a TCGA consensus purity estimation (CPE) obtained by immunohistochemistry analysis and the joined tools. We selected the lowest and highest-purity cases to capture different levels of data quality (Online Methods), and applied CNAqc successfully to rank fits from all callers (Supplementary Figure S23). Strikingly, we found some cases where CNAqc failed all purity estimates, including CPE, but passed the one by ABSOLUTE (see Supplementary Figure S24 for an example with 80% CPE, failed, and 69% ABSOLUTE, passed).

To investigate the frequency of this type of errors, we extended this test to *n* = 1464 TCGA samples from multiple tumour types, retaining cases with at least 200 mutations (Online Methods; Supplementary Figure S25 and S26). We generalised our finding with 901 cases (60% of 1464) where CPE purity was failed by CNAqc, while the purity proposed by ABSOLUTE often passed QC. We assessed that, had we used CNAqc to select the best purity instead of consensus, 785 out of 901 cases (∼88%) would have passed QC, obtaining a purity estimate more precise than the TCGA consensus. This shows convincingly that CNAqc can recover a good purity/CNA estimate, even when a consensus approach is invariably confused, i.e. most consensus inputs misscall purity.

### QC-powered automatic copy number calling pipeline

We used CNAqc to assemble the first automatic copy number calling pipeline (Supplementary Figure S27) to iterate CNA calling until QC is passed (or for a maximum number of steps). This pipeline leverages Sequenza [22] and combines CNAqc purity-adjustment scores together with ranking of alternative solutions determined by the Sequenza algorithm. By the generality of CNAqc, this approach could be extended trivially to other CNA calling algorithms.

We used the Sequenza-CNAqc pipeline to generate clonal copy number data for two colorectal cancer patients with multi-region WGS data associated. From a total of 10 samples with median coverage ∼80x and purity ∼80% (Supplementary Figure S29), the pipeline successfully generated calls that passed QC. Moreover, the tool was also able to identify the true CNA profile when artificially-miscalled copy number profiles were given in input.

## Discussion

Cancer precision medicine, boosted by the large-scale adoption of bulk sequencing in the clinic, will increasingly rely on data generated from landmark cancer genomics programs [19,28,39]. This poses data quality under the spotlight, rendering manual curation and consensus calling strategies that either are difficult to scale, or come with significant bioinformatic overheads. Therefore, automatic procedures to determine the quality (QC algorithms) of mutation calling pipelines are hihgly desired to enable cancer precision medicine [4,40,41].

To the best of our knowledge, CNAqc is the first framework to formalise QC algorithms for bulk assay, leveraging SNVs and indels mutations, along with allele-specific CNAs, tumour purity, and estimations of clonality. CNAqc can be used to process and QC the most common CNAs found in human cancers, using distinct algorithms based on clonality and the complexity of CNAs under scrutiny. In particular, CNAqc can delineate the evolutionary history of subclonal CNAs and quantify the likelihoods of the underlying evolutionary processes, implementing a QC algorithm inspired by tumour evolution principles. Importantly, all algorithms presented in our framework have been validated in controlled single-cell experiments and synthetic simulations. CNAqc can also support downstream analysis using innovative algorithms. In fact, CNAqc is the first model to compute per-mutation CCFs within an information-theoretic uncertainty model for the estimation of mutation multiplicities. This is biologically very relevant for downstream analyses that rely on CCFs, which are at the cornerstone of all copy number timing [17] and tumour evolutionary inference analyses [12–18].

CNAqc was used to analyse bulk WGS and WES data from TCGA, PCAWG and Genomics England. Notably, this included copy number and mutation data generated from more than 10 widely used bioinformatic pipelines, as well as with consensus calls in TCGA and PCAWG. Our analysis proved that sequencing coverage impacts the rate of successful copy number calling, with a clear advantage observed when comparing ∼100x WGS from Genomics England against earlier PCAWG cohorts, both in terms of clonal and subclonal copy number detection, as well as CCFs computation. Particularly interesting results emerged comparing consensus approaches that combine predictions of distinct algorithms. Consensus-based methods are popular in bioinformatics, and rely on the assumption that by gathering data from many tools we can improve the quality of our predictions. While these strategies have obvious computational overhead, the sensitivity of such procedures is largely dependent on the quality of its input methods. Strikingly, we found that in ∼60% of TCGA samples (from a scrutiny of ∼1500 samples) the consensus was not robust to confounders impacting individual tools, and CNAqc was required to identify and fix imprecise consensus purity estimates. In general, the possibility that QC-based methods could substitute or at least augment consensus approaches would make somatic calling pipelines faster and simpler to organise or maintain. In this respect, we released the first automatic CNA calling pipeline that joins a popular copy number calling algorithm with CNAqc.

The quality of QC by CNAqc depends on the capacity to detect VAF peaks, which creates issues when a sample has low purity (e.g, below 20%), low coverage (e.g, below 30x), low mutational burden or when there is significant tumour contamination in a germline sample [42,43]. Most of these limitations are ‘technological’ and can be overcome with higher depth of sequencing and upfront assessment of tumour content. That said, there is an intrinsic limit to how well bulk sequencing designs can capture subclonal copy numbers, especially at the scale of single-cells [44–47]. This limitation of bulk assays may place focus on alternative emerging technologies such as tissue laser capture, though in the short to long term this challenge does not reduce the importance of assessing data quality from widespread bulk sampling.

Summarising, generating high quality copy numbers and mutation data is a necessity for successfully interpreting cancer genomes [12–14,17,48–55]. CNAqc can help to assess whether the quality of the sequencing data is sufficient to ask specific research questions related to tumour aneuploidy, evolution and general molecular profiling. With the ongoing implementation of large-scale sequencing efforts, CNAqc offers a modular solution to augment established pipelines, aiding the self-tuning of bioinformatics parameters based on quality scores. To our knowledge, this is the first stand-alone tool which combines a tumour-evolution perspective with common types of cancer mutations to automatically control the quality of a sequencing assay.

### Data Availability

PCAWG calls are publicly available at (https://dcc.icgc.org/), the ICGC Data Portal; we used the following files:

- Somatic consensus SNVs and indels

○ https://dcc.icgc.org/releases/current/Summary#:~:text=simple_somatic_mutation.aggregated.vcf.
- Somatic allele-specific CNAs

○ https://dcc.icgc.org/api/v1/download?fn=/PCAWG/consensus_cnv/consensus.20170119.somatic.cna.annotated.tar.gz
- Purity and ploidy cohort table

○ https://dcc.icgc.org/api/v1/download?fn=/PCAWG/consensus_cnv/consensus.20170217.purity.ploidy.txt.gz

TCGA calls are publicly available at the GDC Data Portal (https://portal.gdc.cancer.gov),

Results from our analysis of PCAWG and TCGA cohorts have been made available at Zenodo as R outputs: https://doi.org/10.5281/zenodo.6410935

The Genomics England data set can be accessed by joining the community of academic and clinical scientists via the Genomics England Clinical Interpretation Partnership (GeCIP), https://www.genomicsengland.co.uk/about-gecip/. Instructions available at those web pages should be followed in order to join a GeCIP domain.

Multiregion colorectal cancer data is deposited in EGA under accession number EGAS00001003066.

### Software Availability

CNAqc is implemented as an open source R package at

https://caravagnalab.github.io/CNAqc/.

The tool webpage contains RMarkdown vignettes to run analyses, visualisation inputs and outputs, and parametrise the tool. All analyses presented in this paper can be replicated following those vignettes; multiregion colorectal cancer data to replicate our analysis is hosted in the GitHub repository.

https://github.com/caravagnalab/CNAqc_datasets.

### Authors contribution

GC, WC and JH conceived the project. GC, AA and RB formalised the model. GC, RB and SM carried out simulations. MW, NC and GC analysed single-cell data. SM, AdO, AA, JH and GC analysed data. GC implemented the tool. GC and WC drafted the manuscript, which all authors approved in final form.

### Competing interests

The authors declare no competing interests.

## Acknowledgments

The research leading to these results has received funding from AIRC under MFAG 2020 - ID. 24913 project – P.I. Caravagna Giulio. Some research was performed using the Cancer Research UK City of London Major Centre High performance computing facility (colcc.ac.uk) and was also funded by a Wellcome Trust grant (ID: 202778/Z/16/Z). Research on data from Genomics England has been carried out under project “Quality Control of somatic calls from whole-genome sequencing” (research registry identifier 878).

## Methods

### CNAqc

#### Taxonomy

CNAqc supports both GRCh38 and hg19, the two most popular human reference genome assemblies. The tool distinguishes between different types of allele-specific CNAs:

- “simple clonal CNAs”: clonal copy number segments including states of: heterozygous diploid (AB or 1:1), monosomy loss of heterozygosity (LOH) (A or 1:0), copy-neutral loss of heterozygosity (LOH) (AA or 2:0), triploid amplification (AAB or 2:1) and tetraploid amplification (AABB or 2:2);
- “complex clonal CNAs”: clonal copy number segments that generalise simple ones, with any combination of major and minor allele copies (e.g., 3:2, 4:0, 6:1, etc.).These are complex because they require multi-step genetic alterations to accrue from a baseline wildtype 1:1 state (normal);
- “subclonal simple CNAs”: subclonal simple copy number segments in 2 subclones with known proportions.

We curated this taxonomy to balance modelling complexity (simple versus complex), and still cover most of the CNAs reported in PCAWG (Supplementary Figure S1a.), where 36% of clonal CNAs are 1:1, 15% are 2:1, 11% are 1:0, 8% are 2:2 and 8% are 2:0. Simple CNAs in PCAWG are >75% of the whole set of >600,000 segments, and span 93% of all CNA-covered bases (Supplementary Figure S1b.). In the same cohort, most CNAs are clonal, and on average, simple CNAs cover ∼80% of the overall genome per sample (Supplementary Figure S1c-d.). At the subclonal level, Battenberg segments with simple CNAs are ∼70% of the overall subclonal CNAs (Supplementary Figure S1e-f.), and the limitation of handling 2 subclones in CNAqc mirrors the functioning of popular subclonal callers such as Battenberg, ReMixT and CloneHD (Nik-Zainal et al., 2012; Fischer et al., 2014; McPherson et al., 2017).

## Notes

https://caravagnalab.github.io/CNAqc/

https://zenodo.org/record/6410935#.ZGTUrC9ByqA

### Competing Interest Statement

The authors have declared no competing interest.

### Summary of Updates

A new analysis of 235 WGS samples from the Genomics England cohort has been added. The text has been revised.

