## Supplementary material for "Computational validation of clonal and subclonal copy number alterations from bulk tumour sequencing": Methods

602.

59. Jiang Y, Qiu Y, Minn AJ, Zhang NR. Assessing intratumor heterogeneity and tracking longitudinal and spatial clonal evolutionary history by next-generation sequencing [Internet]. Proceedings of the National Academy of Sciences. 2016. p. E5528–37. Available from: http://dx.doi.org/10.1073/pnas.1522203113

60. Gillis S, Roth A. PyClone-VI: scalable inference of clonal population structures using whole genome data. BMC Bioinformatics. 2020;21:571.

61. McInnes L, Healy J, Melville J. UMAP: Uniform Manifold Approximation and Projection for Dimension Reduction [Internet]. arXiv [stat.ML]. 2018. Available from: http://arxiv.org/abs/1802.03426

### Methods

#### CNAqc

**Taxonomy.** CNAqc supports both GRCh38 and hg19, the two most popular human reference genome assemblies. The tool distinguishes between different types of allele-specific CNAs:

- "simple clonal CNAs": clonal copy number segments including states of: heterozygous diploid (AB or 1:1), monosomy loss of heterozygosity (LOH) (A or 1:0), copy-neutral loss of heterozygosity (LOH) (AA or 2:0), triploid amplification (AAB or 2:1) and tetraploid amplification (AABB or 2:2);

- "complex clonal CNAs": clonal copy number segments that generalise simple ones, with any combination of major and minor allele copies (e.g., 3:2, 4:0, 6:1, etc.).These are complex because they require multi-step genetic alterations to accrue from a baseline wildtype 1:1 state (normal);

- "subclonal simple CNAs": subclonal simple copy number segments in 2 subclones with known proportions.

We curated this taxonomy to balance modelling complexity (simple versus complex), and still cover most of the CNAs reported in PCAWG (Supplementary Figure S1a.), where 36% of clonal CNAs are 1:1, 15% are 2:1, 11% are 1:0, 8% are 2:2 and 8% are 2:0. Simple CNAs in PCAWG are >75% of the whole set of >600,000 segments, and span 93% of all CNA-covered bases (Supplementary Figure S1b.). In the same cohort, most CNAs are clonal, and on average, simple CNAs cover ~80% of the overall genome per sample (Supplementary Figure S1c-d.). At the subclonal level, Battenberg segments with simple CNAs are ~70% of the overall subclonal CNAs (Supplementary Figure S1e-f.), and the limitation of handling 2 subclones in CNAqc mirrors the functioning of popular subclonal callers such as Battenberg, ReMixT and CloneHD (Nik-Zainal et al., 2012; Fischer et al., 2014; McPherson et al., 2017).

Overall these assumptions serve to limit the computational complexity of the computational problems approached in CNAqc (Dentro, Wedge and Van Loo, 2017). Simple CNAs are evolutionarily acquired in one step from a heterozygous germline diploid state. For instance, from a wildtype 1:1 a tumour cell might either amplify one allele and reach a 2:1 trisomy, or lose an allele and reach a 1:0 LOH. With slightly more complex biological mechanisms, states such as 2:0 or 2:2 can also be reached in a single evolutionary "move". More complex CNAs require instead articulate modelling; for instance, a 4:0 might be the final endpoint of a tumour undergoing whole-genome doubling, then further amplifying to 4:2 and eventually reaching 4:0. CNAqc attempts some degree of evolutionary inference at the level of subclonal CNAs, where it uses simple CNAs to determine whether the subclones have evolved linearly or branching out of a common ancestor. For instance, if the subclones are 1:0 and 2:0, CNAqc will try to determine whether the tumour branched from 1:1 with 1:0 and 2:0 siblings 0 (1:1→1:0 | 2:0), or if the tumour jumped from 1:1 to 1:0 and then from 1:0 to 2:0 0 (1:1→1:0→2:0), modelling 1:0 and 2:0 as nested, or vice versa (1:1→2:0→1:0). In this formulation there is also some degree of simplification, at least as far as what CNAs are supported to make reasonable inferences.

CNAqc is conceptualised to work with high-resolution - i.e., high purity and coverage - whole-genome and whole-exome sequencing (WGS) data (see Analysis of patient data). For clonal CNAs, the method pools mutations from segments with the same copy numbers (e.g., all 2:1 segments), either across the whole-genome, or per-chromosome. Subclonal segments are instead analysed without any pooling to support algorithms (e.g., Battenberg) that compute segment-specific CCFs. The main challenge with WES or low resolution WGS data is the reduced mutational burden and noise in the VAF, which decreases the signal strength. The key determinant to detect VAF peaks for CNAs is therefore the number of mutations per copy state, with the idea that thousands from a high-quality WGS assay, are certainly better than hundreds from WES or from low-quality WGS. For tumours which are genomically unstable, or exposed to endogenous mutant factors such as smoking or UV-light, or with high mutation rate like microsatellite unstables, the observable mutational burden in exomes might be enough. At the level of subclonal CNAs, the problem is also present since the number of mutations per segment is smaller compared to the clonal counterpart.

In general low-purity or low-coverage data are expected to impact the QC performance, as measured by false positives and negatives rates. In terms of how this impacts the automatisation of the QC step, in our best practices we randomly check samples if the median coverage is lower than 50x and the purity is below 50%. These cutoffs have been determined by cross-referencing simulation results, as well as by the large-scale application of our tool to both WGS and WES data.

#### Simple clonal CNAs

**Expected clonal VAF peaks.** Certain equations for VAF peaks apply to all types of clonal CNAs; subclonal extensions are discussed in a separate section. A bulk is a mixture of tumour and normal cells present in proportion $\pi > 0$ and $(1 - \pi)$, respectively. Each mutation is then present in a percentage $0 < c < 1$ of tumour cells; to introduce our framework, we consider clonal mutations (with $c = 1$). We use a simple equation describing our belief about the position of the clonal VAF peak in the data, assuming the input clonal segments and purity are correct. This equation links all segments with the same allele-specific CNA. In this manuscript, we denote as $n_A : n_B$ allele-specific segments with $n_A$ and $n_B$ copies of the major and minor alleles (e.g., 1:1 has $n_A = n_B = 1$; 1:0 has $n_A = 1, n_B = 0$).

We introduce the multiplicity $m \geq 1$ (or copies) of a clonal mutation mapping: for simple CNAs $1 \leq m \leq 2$, for complex CNAs $1 \leq m \leq n_A$ and $1 \leq m \leq n_B$. As in ASCAT [21], the expected proportion of reads that can be attributed to a mutation with multiplicity $m$ is $m\pi$. The difference between ASCAT and CNAqc is that the former considers germline single-nucleotide polymorphisms (SNPs), while the latter considers somatic mutations (i.e., germline is removed). While the conceptualisation is similar and already appears in [27] although for reasons other than QC, there are at least two advantages in using mutations. First, CNA callers use B-allele frequencies from SNPs and not VAFs, which makes VAFs an interesting orthogonal measurement. Second, certain genome configurations - e.g., whole genome duplications - can not be detected from BAFs, while they can be clearly detected from VAFs.

For segments $n_A : n_B$, the proportion of all reads from the tumour is $\pi(n_A + n_B)$, where $n_A + n_B$ is the ploidy of the segments (not of the overall tumour). For a healthy diploid normal and tumour clonal mutations sitting on $n_A : n_B$ segments, we expect a VAF peak

(1) $v_m(c) = m\pi c / [2(1 - \pi) + \pi(n_A + n_B)]$ .

that for clonal mutations is defined by $c = 1$. This formula describes our belief about the position of the VAF peak in the data, assuming the input segments and purity $\pi$ are correctly estimated. The formula is intuitive: for $\pi = 1$, in an heterozygous diploid segment $(1 = n_A = n_B)$ clonal heterozygous mutations have $m = 1$ and VAF $v_m(1) = 0.5$. Instead, for whole-genome duplication segments $(2 = n_A = n_B)$ under a simple CNA model, clonal mutations in single $(m = 1)$ or double $(m = 2)$ copy give $v_1(1) = 0.25$ and $v_2(2) = 0.5$. Note equation (1) holds for both SNVs and indels, and it can be used to "phase" mutations on the amplified or non-amplified alleles from VAFs, a feature that we use to compute CCFs.

**Conversion among VAF and purity space.** Simple clonal CNAs are the most prevalent type of CNA, and due to their prevalence are the most important ones to QC a copy number profile. The peak-matching algorithm in CNAqc leverages equation (1), and is summarised in Supplementary Figure S4. It works by pooling mutations from segments with same allele counts (genome-level or chromosome-level), and determining a score per mutation multiplicity. Eventually, this score is propagated to the sample with a linear combination weighted by the number of mutations in each segment.

The scores reflect purity adjustments to fit equation (1) better, and can be used to determine a pass or fail status per multiplicity, segment or sample, depending on whether the score is larger or lower than a desired cutoff. This provides a fine-grained QC, where we might decide to fail only a certain portion of the genome. The peak detection strategy take as input $\epsilon > 0$, the upper bound on the error that the user is willing to tolerate on sample purity. For example, if $\epsilon = 0.05$, we can accept a 5% error on the purity; if the true purity was 60%, we would pass a value in [55%, 65%]. Mathematically, the range associated to $\epsilon$ and $\epsilon$ itself are adjusted to account for ploidy and mutation multiplicity, converting error measures from VAFs to purity units (see below).

For every simple CNA we match either 1 or 2 peaks: a single peak is matched for 1:0 and 1:1, two for all others. Here we discuss the strategy to detect a generic peak, assuming to work with copy state $n_A : n_B$ as in equation (1). The tool implements methods (described below) to detect $n$ peaks $d_1, \ldots, d_n$ in the VAF distribution, and match them against equations (1). To compute the match we select one specific $d_*$, in one of two possible ways:

- by closest match, i.e., $d_* = arg_i \, min \, |d_i - v_m|$, $d_*$ is closest to $v_m$;
- by rightmost match: where $d_*$ is computed as above but from the subset of peaks $D = \{d_i > v_m | \ i = 1, .., n\}$.

The default strategy is the first, because the second one can also identify miscalled segments. For instance, LOH segments miscalled as diploid 1:1 lead to an extra VAF peak on the right of the clonal cluster, which would be detected by the rightmost match.

As anticipated, CNAqc can re-scale errors from VAF to purity units, an operation that serves to adapt the input parameter $\epsilon$ to work with different CNAs. The general equation

(1.1) $$\epsilon_m = v_m(\pi + \epsilon) - v_m(\pi) \sim \frac{\partial v_m}{\partial \pi}\epsilon = \frac{2m\epsilon}{[2(1-\pi) + \pi(n_A + n_B)]^2}$$

where $v_m(\pi) = m\pi / [2(1-\pi) + \pi(n_A + n_B)]$ is the VAF as function of purity for clonal mutations, and we truncate the Taylor expansion of $\nu_m(\pi + \epsilon)$ at the first order assuming $\epsilon$ small. This means that, for a purity error $\varepsilon$, the error on VAFs depends on $\pi$ and $m$. Consider, for instance, a 2:1 segment for a tumour with purity 90%, $\varepsilon = 0.05$ (5%) corresponds to an error in the VAF of approximately 1% and 2% for the VAF peaks with multiplicity $m = \{1,2\}$ respectively. Inverting equation (1), one can express the purity as a function of the VAF, ploidy and multiplicity and derive the error propagation formula from the VAF to the purity space. Using the same approach as above, we can treat the purity as a function the VAF by inverting equation (1) with respect to $\pi$ and setting $c = 1$. Then, we shift the VAF by a small error $\epsilon_m$and truncate the Taylor expansion at the first order to get the error propagation formula

(1.2) $$\pi(v_m) = \frac{2v_m}{m + [2-(n_A+n_B)]v_m},$$

(1.3) $$\pi(v_m + \epsilon_m) - \pi(v_m) \sim \frac{\partial \pi}{\partial v_m}\epsilon_m = \frac{2m\epsilon_m}{[m + v_m(2-(n_A+n_B))]^2}.$$

Peaks are matched by including a VAF tolerance $\epsilon_{VAF} > 0$, which helps ameliorate the fact that we do not explicitly model noise affecting peak detection. The intervals

(1.4) $$I_m^{VAF} = [d_* - \epsilon_{VAF};\ d_* + \epsilon_{VAF}] \quad \text{and} \quad I_m = [v_m - \epsilon_m, v_m + \epsilon_m]$$

are created with centre at $d_*$ with size $2\epsilon_{VAF}$, and tested for overlap with the interval. The clonal peak for multiplicity $m$ is matched by $d_*$ only if the intervals overlap, i.e., $|I_m^{VAF} \cap I_m| > 0$ ,

The QC status per copy state with two possible multiplicity values is defined by taking the status of the peak associated with the largest number of mutations $n_m$, i..e, as majority voting weighted by mutation counts. The value of $n_m$ is determined by binning the VAF distribution with 100 bins from 0 to 1 (size 0.01), and counting the number of mutations per bin of the matched peak. In this way, CNAqc passes a copy state if the tallest of its peaks is a pass, and is associated with more mutations than any failed peak. The sample-level QC status is based on an error metric that uses the actual distance between the centres of the intervals, $d_*$ and $v_m$, which is given by $d_* - v_m$ and described below.

**Sample-level score.** An error metric is assembled across simple clonal CNAs to determine a sample-level score. Consider $w_k$the normalised number of mutations mapped to copy state $k$, rescaled by 2 if the CNA is supposed to have two peaks. For every copy state and multiplicity, we have a pass or fail status from peak detection. We split pass ($P_k$) from fail ($F_k$) peaks, and define two scores by linear combination

(1.5) $$\lambda_k^{\text{PASS}} = \sum_{d_*^m \in P_k} w_k(d_*^m - v_m^k)$$
$$\lambda_k^{\text{FAIL}} = \sum_{d_*^m \in P_k} w_k(d_*^m - v_m^k)$$

where the subscript denotes the copy state (i.e., 1:0), and $d_*^{\ k}$ denotes the peak matched for multiplicity $m$ in state $k$. We define the CNAqc overall sample score $\lambda$

(1.6) $$\lambda = \sum_{k \in P_k} (\lambda_k^{\text{PASS}} + \lambda_k^{\text{FAIL}})$$

as a linear combination of terms that can be either positive or negative, depending on whether the matched peaks are on the right or left of the expected peaks. The sample score $\lambda$is a weighted mean since by construction all the $w_k$ sum to one, and the terms constitute a partition.The overall sample status is finally taken by comparing $\lambda_{k,}^{PASS}$ and $\lambda_{k,}^{FAIL}$ and selecting the status corresponding to the largest.

**Peaks detection algorithms.** CNAqc implements multiple strategies to detect $n$ peaks $d_1, \ldots, d_n$ in the VAF distribution:

1. Kernel-based: a smoothed VAF is obtained via kernel density estimation with default adjustment and fixed bandwidth. Peaks are then estimated from the discretized smoothed signal using R packages for peak detection, and removing peaks with density below 1/20 (empirical cut) of the maximum peak.
2. Mixture-based: via finite Dirichlet Binomial mixtures a peak is associated with each Binomial probability, using our BMix [30] package (https://caravagn.github.io/BMix/),

The latter strategy is inspired by subclonal deconvolution methods, and computes the model density for $w$ clusters (default $w < 5$), with model-selection to optimise $w$ using the Integrated Classification Likelihood score [30]; the likelihood is

$$f(X \mid \boldsymbol{\pi}, \boldsymbol{p}) = \prod_{x \in X} \sum_{i=1}^{w} \pi_i \mathrm{Bin}(r_x \mid n_x, p_i) \tag{1.7}$$

where $\pi_i$ are the mixing proportions of the mixture, not to be confused with sample purity. Here we use a Binomial likelihood for $r_x$ successes determined as the number of reads with the mutant covering mutation $x$, $n_x$ is the total trials given by sequencing depth at the locus, and $p_i$ the Binomial probability. Assuming that calls have passed QC, then $p_i$ is the expected theoretical VAF from equation (1). A key advantage of BMix over other deconvolution tools is the fast maximum likelihood implementation, with full access to the model parameters.

#### Complex clonal CNAs

Complex clonal CNAs are also QCed by a peak detection algorithm, but with a procedure that is simpler than the one proposed for simple clonal CNAs. So, while the sample-level QC is determined by simple CNAs, this procedure helps understanding if more complex states of aneuploidy inferred by a copy number caller are supported by data.

The procedure implemented in CNAqc works by matching VAF peaks, using a subset of the algorithms discussed above.Expected peaks are considered by applying equation (1) with mutation multiplicity ranging from 1 to the maximum between the major and minor allele counts of the considered segment (e.g., for a 4:2 copy state multiplicities from 1 to 4 are tested). Somatic mutations, as in the previous analysis, are pooled among segments with the same CNA, either across the whole genome, or at the chromosome level.

To make this analysis faster, peaks in complex CNAs are inferred solely by the KDE heuristic, matching with the "closest" modality adopted for simple CNAs. Moreover, for every expected peak the input  parameter (purity tolerance) is used without performing any conversion from purity to VAF space. For complex CNAs a table is compiled with every expected data peak, which depends on the mutation multiplicity, and a pass or fail status. No segments-level or sample-level scores are assembled in this case.

#### Subclonal CNAs

CNAqc can QC subclonal CNAs for 2 subclones with proportions $\rho_1$ and $\rho_2$ so that $\rho_1 + \rho_2 = 1$, and with simple CNAs. Compared to clonal segments, the QC of subclonal CNAs requires one to elicit the evolutionary trajectory of the subclones, because VAFs peaks depend on the particular phylogenetic relationship between the subclones. In particular, VAF peaks depend on whether subclones originate upon branching from an unobserved ancestral state, or whether they evolve linearly. If they branch they are siblings, otherwise only one descends directly from the ancestral state. For instance, if the subclones are 1:0 and 2:0 with ancestral 1:1 it could either be that an heterozygous diploid cell, upon division, originated two distinct types of LOH (branching model 1:1 → 1:0 | 2:0), or could be that the diploid ancestor first loses one allele (generating 1:0), and then amplifies the remaining allele (linear model 1:1 → 1:0 → 2:0). One might also think that longer paths (e.g., 1:1 → 2:1 → 2:0 → 1:0) are less likely, and assume that the shortest one is followed in most situations.

Fixing the time in which one subclone separates from the other is crucial for determining the multiplicity of the mutations accumulated before the split, which are shared across the subclones. Suppose, for instance, that subclones originated independently from a 1:1 cell: only mutations present in the original clone (ancestral mutations) will be shared, and subsequent CNAs will alter their multiplicity independently in each subclone. Conversely, if subclones evolved together up to a certain configuration before one of the two acquired a further alteration and expanded independently, the multiplicity of shared mutations will depend on that of the last shared copy state.

In order to account for the multiplicity of both shared and private mutations, CNAqc implements a recursive tree-generation algorithm that first *i)* reconstructs the evolutionary tree (linear versus branching) that led a starting cell to develop into two subclones, and then *ii)* computes multiplicities based on the tree. The starting state is a 1:1 diploid cell (default), but can be changed. The algorithm simulates progression from the initial to the final states by performing single allele duplication, deletion and mutation accumulation. Branching and linear scenarios are considered separately, and for every progression from X to Y only the shortest path is retained. Given paths, shared and private mutation multiplicities are determined based on the ordering of the amplifications and deletions, and the expected peaks are determined.

Consider two subclones $n_{A,1}:n_{B,1}$ and $n_{A,2}:n_{B,2}$, the expected peak for a shared mutation with multiplicity $m_1/m_2$ is

(1.8) $$v_{\{m_1,m_2\}} = \frac{(m_1\rho_1 + m_2\rho_2)\pi}{2(1-\pi) + \pi(\rho_1(n_{A,1}+n_{B,1}) + \rho_2(n_{A,2}+n_{B,2}))}.$$

The total multiplicity of mutations shared is the sum of the total multiplicities. Private mutations found solely on subclone $i \in \{1,2\}$ have multiplicity $m \in \{1,2\}$, and the expected peaks are

(1.9) $$v_{m_i} = \frac{m_i\rho_i\pi}{2(1-\pi) + \pi(\rho_1(n_{A,1}+n_{B,1}) + \rho_2(n_{A,2}+n_{B,2}))}.$$

Supplementary Figure S3 shows the application of equations (1.8) and (1.9) to subclones with clonal fraction $\rho_1 = 0.75$ and $\rho_2 = 0.25$, and tumour purity $\pi = 0.8$. In Supplementary Figure S3a the subclones have CNAs $n_{A,1}:n_{B,1} = 1:1$ and $n_{A,2}:n_{B,2} = 1:0$, and have three expected peaks :

- $v_{\{m_1=1,m_2=1\}}$, for shared mutations;
- $v_{m_1=1}$, for private mutations on the first subclone and;
- $v_{m_2=1}$, for private mutations on the second subclone.

Since the multiplicities are either 1 or 0, expected peaks for the linear and the branching models are equivalent, though they correspond to different mutation groups. For example, a shared peak at 37% VAF comes from

$$v_{\{m_1=1,m_2=1\}} = \frac{(m_1\rho_1 + m_2\rho_2)\pi}{2(1-\pi) + \pi(\rho_1(n_{A,1}+n_{B,1}) + \rho_2(n_{A,2}+n_{B,2}))} = \frac{(0.75 + 0.25)0.8}{2(1-0.8) + 0.8((2)0.75 + 0.25))} = 0.37,$$

and two private peaks instead come at $v_{m_1=1} = 0.28$ and $v_{m_2=1} = 0.09$.

More interesting is the case of two 2:1 and 2:0 subclones, where the branching and linear models have different peaks. The branching model (AB $\rightarrow$ AAB | BB) has the amplification and the deletion of the same allele (A) for the two subclones, and the amplification of the other (B) in one of the subclones. In this way, the major allele of one subclone (B) is the minor allele in the other. Five peaks are expected, of which only $v_{\{m_1=1,m_2=2\}} = 0.38$, is shared and corresponding to mutations present on the ancestral allele, while $v_{m_1=2} = 0.46$, $v_{m_1=1} = 0.23$, and $v_{m_2=2} = 0.15$ and $v_{m_2=1} = 0.08$, correspond to private peaks for the subclones.

Instead, the second branching model (AB $\rightarrow$ ABB | BB) sees the independent amplification of the same allele (B) in the two subclones. Private peaks remain as in the above model, but shared peaks have a different position with $v_{\{m_1=2,m_2=2\}} = 0.61$. Finally, in the only possible linear model (AB $\rightarrow$ AAB $\rightarrow$ AA) the 2:0 subclone arises from the 2:1 upon loss of the minor allele. This scenario is quite different from the previous two, as the number of expected peaks becomes four, two of which ($v_{\{m_1=2,m_2=2\}} = 0.61$, $v_{\{m_1=1,m_2=1\}} = 0.31$) are shared and two ($v_{m_1=1} = 0.23$, $v_{m_2=1} = 0.08$) are private.

#### CCF estimation

**From VAFs to per-mutation CCFs.** The equation [21,27] to compute CCFs in CNAq converts the VAF (observed )$v > 0$ of a mutation with multiplicity $m$ (to be estimated) into the CCF $c$ as follows

(2) $c = [v[(n_A + n_B - 2)\pi + 2]/m\pi$ .

Per-mutations CCFs derive from VAFs, and for this reason harbour the same observational noise. For example, an heterozygous clonal diploid mutation in a pure tumour has 50% theoretical VAF and CCF $c = 1$, because 100% of cells harbour the mutation. If it sits on a 2:1 segment, its theoretical VAFs are either 33% - $m = 1$ out of 3 copies - or 66% - $m = 2$ out of 3 copies - but still $c = 1$. However, since VAFs are observed with Binomial noise their observed values spread around the theoretical ones (e.g., around 33%), then CCFs are subject to the same noise rescaled by equation (2). This leads to a contradiction with the term "fraction" which by definition cannot exceed 1 but, for per-mutations CCFs converted by equation (2), VAF-associated noise spreads CCF values around their theoretical estimate. In this sense, the CCF of a clonal mutation spreads around $c = 1$ using equation (2), as already reported in other popular papers - see, e.g. Figure 12 and 13 in Methods of [16]. Nonetheless, subclonal deconvolution methods, e.g. [13] can filter out Binomial noise and return values that range in [0,1]. However, this computation is carried out after CNAqc, as it goes beyond the idea of performing QC.

**CCF computation algorithms in CNAqc.** Tumour subclonal deconvolution algorithsm, e.g, PyClone [13], denoise CCFs by computing cluster-level rather than per-mutation CCFs. Instead, CNAqc determines CCFs for mutations mapped to simple clonal CNAs, and a pass or fail status for the CCFs, determined by a metric to filter out mutations with uncertain estimates. In this way, CCFs by CNAqc can be used to QC tumour evolution inferences that leverage CCF clusters, for instance [30,56]. A limitation of CNAqc compared to other methods is to consider a subset of CNAs; this is motivated by the difficulty in phasing, from VAFs, mutation multiplicities from any CNA segment (complex and subclonal). Notably, the computations by CNAqc are however much faster, compared to analogous deconvolution tools [13,15].

CNAqc offers two approaches to compute CCFs: *i)* an entropy model that uses Binomial mixtures peaked at VAFs from equation (1), that phases mutation multiplicity using the mixture latent variables (capturing uncertainty from the latents), *ii)* a rough model that uses the mixture, but does not model uncertainty. The former model, if there is too much uncertainty on the multiplicity of a mutation, can leave it undetermined (as its CCF), and return NA; this is how uncertainty is reported in CNAqc. The rough method, instead, will always assign a multiplicity $m \in \{1,2\}$. The final QC status of some segments (e.g., all 2:1) is determined from the proportion of mutations with available CCF, with the idea to pass QC only if the number of assignable CCFs exceeds a user-defined threshold (default 10%). Therefore, the latter computation method will always pass QC because it does not implement uncertainty.

We first detail the rough approach; we describe the case of 2:0, 2:1 and 2:2, the others being trivial. To initialise a mixture:

1. we build two Binomial densities from the theoretical expectations of the VAF peaks, i.e., $v_1$ and $v_2$, depending on the copy state, as defined in equation (1). This will create, for instance, one Binomial with parameter $p = 0.33$ and one with $p = 0.66$ for a pure ($\pi = 1$) tumour and 2:1;
2. We fix the number of Binomial trials to the median coverage of the mutations, and compute the 1% and 99% quantiles of the data distributions to obtain a VAF range around each peak.
3. Finally, we count mutations that, according to VAF, map to either one or the other computed range. The number of mutations $n_1$ and $n_2$, associated to multiplicity $m = 1$ and $m = 2$, is then used to obtain the mixing proportions $\boldsymbol{\pi}_1 = n_1/(n_1 + n_2)$ and $\boldsymbol{\pi}_2 = 1 - \boldsymbol{\pi}_1$ to complete the model .

With these parameters, denoting by $B(x \mid v_m)$ the Binomial likelihood for mutation $x$ with multiplicity $m$, we can compute the mixture likelihood

(2.1) $$f(\mathbf{X} \mid v_m) = \prod_{x \in \mathbf{X}} \sum_m \boldsymbol{\pi}_m B(x \mid v_m)$$

In a mixture model we have latent variables $z$ as a matrix of mutations by clusters, for which we define, the probability of assigning read counts data for mutation $n$ to component $i \in \{1,2\}$

(2.2) $$\mathbf{z}_{n,i} = f(X \mid v_i)/[f(X \mid v_1) + f(X \mid v_2)]$$

With these latents, every row of matrix $z$ is a categorical random variable reporting the probability of assigning $m = 1$ or $m = 2$ to a mutation, for which we define the entropy

(2.3) $$H(\mathbf{z}_n) = -\mathbf{z}_{n,1} log(\mathbf{z}_{n,1}) - \mathbf{z}_{n,2} log(\mathbf{z}_{n,2}) .$$

The entropy is maximal if $\mathbf{z}_{n,1} = \mathbf{z}_{n,2}$, i.e., the mutation is equally likely in single and double copy, and is therefore uncertain to be assigned. As opposite, the entropy is minimal if $\mathbf{z}_{n,1} = 1$ and $\mathbf{z}_{n,2} = 0$, or vice versa. If the entropy is low, the mutation is then difficult to phase to single or double copy mutations, using VAFs. The shape of the entropy resembles - by construction - a growing curve with a central spike, which we use to create a simple criterion to discriminate high from low entropy. The geometric intuition of this criterion is that at the crossing of Binomial densities peaked at $m_1$ and at $m_2$, if the $H(\mathbf{z}_n)$ is high we cannot confidently phase mutation multiplicities. The amount of Binomial overlap depends on coverage and purity, which is the technical reason CCF is more uncertain for low resolution data.

CNAqc uses a simple peak-detection heuristic (similar to the one for QC) to inspect $H(\mathbf{z}_n)$ and determine peaks $\{h_1, h_2\}$ around the spike. Every mutation in the range

$$I_{NA} = [h_1, h_2] \qquad (2.4)$$

cannot be confidently assigned multiplicity values, and are therefore undetermined using the entropy method. Their CCF is also reported as an NA value (Not assigned).

The rough approach works as opposite, as it determines the midpoint $o = v_1 + (v_2 - v_1)\boldsymbol{\pi}_1$ between the two expected theoretical VAF peaks $v_1$ and $v_2$, given the mixing proportion $\boldsymbol{\pi}_1$ of the first mixture component. The midpoint is computed by weighting each of the two peaks proportionally to the number of mutations that appear underneath each peak, which we compute like with the entropy method. The midpoint is a cut: $x < o$ are phased to a single copy, values above to two copies. This procedure requires data with good general quality because it assumes that all mutations can be phased correctly by a hard VAF split, a fact that depends largely on coverage and purity.

When multiplicities have been determined, CCFs are computed with equation (2).

#### Other features

**Genome fragmentation detection.** Some recently identified patterns of somatic CNAs can be attributed to the presence of highly fragmented tumour genomes, termed chromothripsis and chromoplexy, or localised hypermutation patterns, termed kataegis [57]. While these can be identified using dedicated tools, CNAqc offers a simple statistical test to detect the presence of potential over-fragmentation in a region of interest, a prerequisite that could point to the presence of such patterns. CNAqc analysis does not substitute dedicated tools, but provides preliminary information to determine what parts of the genome might be run with ad hoc methods.

At the level of chromosome arms (1p, 1q, 2p, 2q, etc., or subsets), CNAqc uses the length CNA segments to classify “long” and “short” fragments with a cut parameter $\mu > 0$ (default 0.2), and a segment longer than a fraction $\mu$ (rescaled to 100) of the arm is considered long. Recent evidence from large pan-cancer studies can be used to calibrate this parameter to cancer-specific values [5].

Then, a null hypothesis is used to compute a p-value using a Binomial test based on $k$, the number of trials given by the total segments in the arm, and the observed number of short segments $s$. The Binomial distribution for $H_0$ is defined by $\mu$, and the null is the probability of observing at least $s$ short segments. CNAqc defines a one-tailed test for whether the observations are biassed towards short segments, adjusting the p-value for family-wise error rate by Bonferroni, i.e., dividing the desired $\alpha$-value by the number of tests. This test is applied to a subset of chromosome arms with a minimum number of segments, and that "jump" in ploidy by a minimum amount (empirical default values estimated from trial data). The arm-level jump is determined as the sum of the difference between the ploidy of two consecutive DNA segments. These covariates are similar to those used to infer CNA signatures from single-cell low-pass WGS [11].

**Segment smoothing.** Smoothing is an operation that can be carried out, at the level of clonal segments, before testing for over-fragmentation. This operation does not affect the ploidy profile of the calls, but reduces the amount of breakpoints that otherwise inflate the p-value of the Binomial over-fragmentation test in CNAqc.

We implemented this operation to reduce the number of segments reported by a caller, because we observed that in real data several callers break contiguous segments even without actual copy number changes (i.e. the same numbers for the major and minor alleles are reported, but a breakpoint is present to break a segment). These types of behaviours are arguably linked to the segmentation algorithm of the caller, and its ability to call segments over a certain genome length. Therefore, in CNAqc, by smoothing we merge two contiguous clonal segments if they have exactly the same allele-specific profile. The smoothing procedure is controlled by a distance parameter with 1 megabase as default value, which avoids merging segments that are above that distance apart.

**Chromosome-level analyses.** CNAqc can perform QC-based analysis at the chromosome level, namely for each chromosome separately. This functionality can be useful to spot samples where the estimate of the bulk purity is correct, but there are large segments with miscalled allele-specific segments (e.g., large sections of the genome that are called triploid while they should be diploid).

We explain the advantage of this functionality by modifying the calls for the PCAWG hepatocellular carcinoma ca5ded1c-c622-11e3-bf01-24c6515278c0. First, we retrieved SNVs mapped to diploid (1:1) and triploid (2:1) segments; genome-wide allele-specific consensus CNAs are characterised by ploidy 2 and a purity of ~85%. We then simulated the unlikely case in which a copy number caller fails to call the diploid regions as such, and instead assigns them a triploid 2:1 state (Supplementary Figure 2a-c).

Genome level peak analysis computes a quality score for pooled 2:1 segments, failing the whole-triploid solution at the correct purity, and proposing a purity correction of ~7% (Supplementary Figure 2c). For this weird case, however, the chromosome-level CNAqc analysis can identify the source of error (Supplementary Figure 2d). In this case we easily verify that CNAqc fails the triploid solution for each chromosome that contains mostly non-triploid segments – chromosomes 2, 3, 4, 6, 7, 10, 11, 13, 14 ,15, 16, 18, 19, 20 and 21 – while CNAqc passes the ones containing a significantly large triploid region – chromosomes 1, 5, 8 and 17. While these types of errors in the data are unlikely because a copy number should detect different input depth ratios and B-allele frequencies, this type of analysis can be helpful to inspect, at narrower resolution, the quality of the input segments.

#### Simulations, validation and comparison to deconvolution tools

**Peak detection (base simulations).** We tested CNAqc on a synthetic dataset of ~20,000 tumours, generated to mimic data that we observed in real patient tumours. We first simulated synthetic VAFs from clonal CNAs generated with breakpoints distributions following Poissons (6 segments per chromosome, on average, and a Dirichlet copy state concentration 1 for 1:0, 1 for 2:0, 6 for 1:1, 2 for 2:1 and 1 for 2:2). Then we simulated Poisson-distributed coverage with median depth 30x, 60x, 90x and 120x, and set purity to 0.4, 0.6, 0.8 or 0.95. The idea of this test was to simulate a tumour with purity $\pi$and run CNAqc with an input purity that contained a positive or negative error $\varepsilon_{err}$, i.e., we imputed CNAqc purity $\pi + \varepsilon_{err}$. Then, for different values of the input tolerance $\epsilon$, i.e., the maximum purity error we want to tolerate in CNAqc, we run the tool with default peak-matching parameters and perform quality control. Ideally, when the input error $\varepsilon_{err}$ is lower than tolerance $\epsilon$, $\varepsilon_{err} < \epsilon$, CNAqc should pass the sample.

We performed QC applying an error on the purity in range [0; 0.2] with intervals of length 0.02, setting a tolerance on the purity error ranging in [0.01; 0.05] with intervals of length 0.004. We tested CNAqc on 100 simulated tumours for any combination of all the parameters and consistently observed that, as the purity error $\varepsilon_{err}$ exceeds tolerance $\epsilon$, the proportion of failures approaches 100% (Supplementary Figure S6). For instance, setting a tolerance parameter of 2%, we can accept a purity error of 5% at most. Over this threshold the proportion of failed samples reached maximum at ~7%. One can check this behaviour for the samples of purity 0.95 and coverage 90x: for a tolerance of ~2%, the proportion of rejected samples is close to 0% when the purity error is smaller than 5%, it increases to 70-75% for a purity error of ~5/6%, while for a purity error of ~10% the fail proportion is 100%. From the test we also observed that the ability of CNAqc to detect samples with incorrect purity improves consistently as we increase coverage, with this effect more evident for samples with high purity. For the same tumours we also computed CCFs and the proportion of mutations for which CNAqc could not phase multiplicity (for 2:0, 2:1, 2:2). We see the percentage of unassignable mutations (Supplementary Figure S7) to decrease as we increase coverage and purity, meaning that the computation of CCFs and multiplicities depends on these parameters. The observed trend was expected, since at low coverage and purity we have the overlaps between clonal clusters which makes it harder to phase multiplicity from VAFs.

**Validation with single-cell copy number data.** We validated the methodologies implemented in CNAqc by adopting complementary single-cell copy number data. We collected low-pass single-cell data using the Direct Library Preparation (DLP+) protocol from an ovarian cancer cell line [31]. DLP+ is an amplification-free library preparation protocol to generate high-resolution single-cell WGS data suitable for cell-level calling of both CNAs and SNVs. We used this type of data to assemble monoclonal and polyclonal pseudo-bulk populations, and validate all the functionalities of CNAqc.

We first clustered cells with similar allele-specific CNAs which we computed using SIGNALS [32]. These clusters correspond to monoclonal populations composed of 100% tumour cells, and are characterised by specific CNAs (Supplementary Figure S9). Then we obtained SNVs per cell, generated also in [31]. From cluster assignments and read counts per cell, we generated a pileup of read counts per clone (sum of both reference and alternative allele counts for all cells in a cluster/clone) mimicking a WGS assay for each tumour clones. We then selected clusters G (111 cells), H (77 cels) and I (177 cells) because among the larger clusters they are the ones with the least noisy VAFs and the most common monosomy and tetrasomy segments. We used CNAqc to QC the expected purity of 100% (purity by assembly) per clone (Supplementary Figure S10). As expected, our model assigned a pass score to all datasets, for both simple and complex CNAs (3:0 and 4:0).

Then, we tested how accurate the predictions of CNAqc are in terms of purity correction estimates, if one uses a wrong input purity. From the cases above (true purity 100%), we imputed a purity in the form $1-\varepsilon$, where $\varepsilon$ models the error, and measured the $R^2$ correlation between $\varepsilon$ and $\lambda \in \mathfrak{R}$, the purity adjustment returned by CNAqc (Supplementary Figure S11). Peak analysis was run multiple times decreasing input purity from 100% to 90% (1% step, 15 repetitions per point). The tested samples from Supplementary Figure S9 using (a) a pileup of clonal CNAs common to clusters G, H and I (Supplementary Figure S12), in 1:0 and 2:0 regions, (b) cluster G restricted to 1:0 segments, (c) cluster H restricted to 1:0 segments and (d) cluster I restricted to 1:0 and 2:0 segments. In every case we measured that the proposed correction is in perfect agreement with the input mismatch (correlation coefficient $0.88 < R^2 < 0.99$, p-value $p < 2.2e-16$), therefore showing that purity correction estimates in CNAqc are precise.

Finally, we measured accuracy to QC subclonal CNAs (Supplementary Figures S10) by merging some clusters from previous tests into larger clones with more mutations; this was necessary since single clusters were quite small and had few SNVs mapping on subclonal CNAs. We pooled cells from clusters G, H and I in Supplementary Figure S9, and retained clonal CNAs common to all clusters, plus subclonal segments where clusters H and I have the same CNA and differ from cluster G. Then, we mixed all the cells (111 for G, 77 for H and 177 for I), obtaining a mixture with ~70% cells from merged cluster H+I, and ~30% from cluster G. CNAqc could easily validate clonal CNAs as in the previous tests. In this sample we performed QC of 2 subclonal CNAs on chromosomes 4 and 11, harbouring 323 and 271 SNVs each (Supplementary Figures S10). CNAqc detected the expected peaks, therefore supporting the presence of subclonal CNAs: we could validate a mixture of 1:1/2:2 populations, and a mixture of 1:1/2:1 populations. In both cases, peaks from the linear and branching evolution models were observed, making it hard to decide precisely what evolutionary model explains best the origin of these populations. In order to stress test the evolutionary modelling underneath our QC procedure for subclonal CNAs, we also took further data from [33] and generated allele-specific CNAs for these cells using SIGNALS [32]. This time we also phased alleles (Supplementary Figures S11), which allowed us to identify a sample with clear subclonal CNAs and allelic imbalance, consistent with the original publication. In particular, we found a set of cells with monoclonal 2:1 chromosome 1, and polyclonal chromosomes 2-4 (2 clones). On chromosomes 3 and 4 the populations were found to be triploid and tetraploid, while on chromosome 2 they were found to be triploid with mirrored allelic imbalance, as reported in the original publication. In practice, on chromosome 2 one population (58% of cells) had genotype AAB and the other (42% of cells) was ABB, where A and B are the alleles of the unobserved ancestral diploid population (or, in our notation, they were 2:1 and 1:2). CNAqc could validate the subclonal CNAs in all the chromosomes (Supplementary Figures S12). The case of chromosome 2 was particularly interesting, because the tool compared the evolutionary models

1. A1B1 → A1A2B1 | A1A2B1 (branching with imbalance)
2. A1B1 → A1A2B1 | A1B1B2 (branching with mirrored imbalance)
3. A1B1 → A1A2B1 → A1A2B1 (linear with imbalance)

Models 1 and 3 predict the same peaks, with shared mutations peaking at 0.667 and 0.333, and private mutations peaking at 0.192 and 0.141. Model 2 instead predicted shared mutations (in 3 alleles out 6 due to imbalance) with peaks (2*0.58+1*0.42)/(0.58*3+0.42*3) = 0.57 and (1*0.58+2*0.42)/(0.58*3+0.42*3) = 0.47. The data distribution showed a clear peak at VAF around 50%, and CNAqc was therefore able to validate the subclonal segment and identify the correct branching evolution model A1B1 → A1A2B1 | A1B1B2 with mirrored allelic imbalance.

Finally, we sought to use single-cell data also to validate mutation multiplicity phasing in CNAqc, while accounting for uncertainty in the estimate. One limitation of the data at hand was that the multiplicity of input mutations at the single-cell level is unknown. Therefore, we opted to validate CNAqc computations by checking if mutations flagged as uncertain from the pseudo-bulk are also difficult to phase at the single-cell level, which seemed a reasonable test for these data. Using single-cell data for cluster A (Supplementary Figure S9), we computed per-cell VAFs for mutations in 2:1 regions with good mappability and quality scores. Second, we computed per-cell multiplicity per mutation: as with bulk, we assigned one copy ($m = 1$) if the single-cell VAF was closer to 0.33 than to 0.66, and two copies ($m = 2$) for the opposite case (closer to 0.66 than to 0.33). We used a majority score from all cells to vote for multiplicity and resist noisy VAFs from single-cells (caused by low-coverage per-cell <5 reads, not shown). We also registered the proportion of cells that vote for single or double copy. Then, we computed VAFs from the pseudo-bulk of these segments (Supplementary Figure S15a), and identified mutations which CNAqc phased as uncertain in terms of multiplicity. We compared CNAqc assignments from bulk to majority voting with single-cells, after classifying consensus in three ranges: >85% (high), >50% (intermediate) and <50% (low). Results (Supplementary Figure S15b) show that CNAqc assignments of multiplicity from pseudo-bulk match assignments from single cells for 83% of mutations, and for 98% of mutations with consensus >85% among single cells, while CNAq considers uncertain the phasing of a set containing 13% of the mutations (40% of mutations with low single-cell consensus). Notably, only for 3% of the mutations CNAqc phasing fails to match consensus-based phasing. Finally, we computed the histograms of voting values split by CNAqc mutation assignment (one copy, two copies and uncertain, Supplementary

Figure S15c), and we observed mean voting support 70% for mutations assigned one copy, 61% for two copies and 43% for uncertain mutations. Overall these analyses show that the mutations flagged as uncertain by CNAqc are largely the same for which single-cell multiplicities are difficult to estimate.

**Automatic $\epsilon$-calibration via false positive rate curves.** The result of sample-level QC (pass or fail) depends on the maximum purity-error $\epsilon > 0$ specified by the user. CNAqc offers a function to automatically determine what purity error parameter should be used, for a particular combination of coverage and tumour purity, in order to minimise the false positive rate (FPR) of the tool, as determined by simulations.

To calibrate this functionality (Supplementary Figure S8a) we sampled 100 distinct tumour genome segmentations, spanning sample purity $0.15 \leq \pi \leq 0.9$ and median coverage $20 \leq cov \leq 120$. For each tumour and each purity/coverage value, we tested the input purity error $0.01 \leq \epsilon \leq 0.1$ (1%-10%) discretized by 1%; and for each of these values we sampled 10 datasets mimicking a CNAqc input run with tumour purity of the form $\pi + \epsilon + \varphi$ where $\varphi \sim U[0, 0.03]$. So, we have imputed to the tool a purity that is close to the true one ($\pi$) modulo the tolerance ($\pi + \epsilon$), but positioning the tool in the scenario in which the actual input should be failed because $\varphi > 0$. Note that since the error margin is 3%, this borderline scenario represents a configuration in which CNAqc should fail a sample, but the task is difficult because the input is “close” to the cut point where the sample could be passed (theoretically). In this way, we could compute the FPR for each value of purity/coverage as a function of $\epsilon$, which were used to fit a generalised linear model (GLM) for every purity/coverage value (Supplementary Figure S8a). Results from these tests give the expected results; in particular, we clearly observe gradients for both purity and coverage, and the slope of the regression correlates with data quality (higher resolution data allows lower FPR for broad values of $\epsilon$). For instance, with coverage 20 and purity 0.15 the lowest FPR is still 30% at $\epsilon = 0.01$ - because data quality impacts peak matching - and peaks at 60% at $\epsilon = 0.1$. For better quality (e.g, coverage 120, purity 0.9), instead, FPR is well below 10% even with very stringent $\epsilon = 0.01$, and remains substantially low for larger $\epsilon$.

The algorithm to determine which $\epsilon > 0$ minimises FPR for a particular combination of coverage and tumour purity, works as follows. First, one selects the maximum FPR accepted $\mu > 0$ (default 10%); the GLM fits of the training set are used to invert the FPR and determines the largest $\epsilon$ with desired FPR below $\mu$ (Supplementary Figure S8b,d). If the regressed $\epsilon$ exceeds some input range of values (determined by the user), the regressed value is capped. All the regressed values are then interpolated with a 2D Akima non-smoothing spline that gives good fits to curves with a second derivative that changes rapidly [58]; the required point estimate for $\epsilon$ is determined from the interpolation of the spline values (Supplementary Figure S8c,e). The interpolation is carried out only in the range of values of the training set, and constraints on $\epsilon$ required by the user are finally enforced (e.g., so that one can determining the $\epsilon$ which associates with FPR $\mu < 0.7$ while requiring no purity errors below 5%).

**Comparison to deconvolution methods.** Some of the functioning of CNAqc is inspired by the design of subclonal deconvolution methods [13,15,17,27,30,50,56,59]. Therefore, we sought to compare CCFs by CNAqc with the one obtained by Ccube (default parameters), a CCF-computation method developed by the PCAWG Evolution and Heterogeneity Working Group [34].

In Supplementary Figure S16 (panel a) we show the correlation among the CCF values computed by Ccube and CNAqc (entropy method) in PCAWG. In the plot we annotate the proportion of cases, split by copy state and mutation multiplicity, where the estimates are different after rounding to the second digit. We observe that the tools report the same CCF for ~99% of the analysed mutations, whenever CNAqc identifies a reliable CCF value. We remark that a feature of CNAqc is reporting the percentage of mutations where the CCF cannot be unequivocally determined. In the above statistics, the CCF values are therefore computed only for mutations where the uncertainty is not present in CNAqc. The information regarding uncertainty is however very helpful to integrate CNAqc with other tools for CCF computations, as we show with two examples from our test. In Supplementary Figure S16 (panels b-g) we report an example PCAWG case where the CCFs are in perfect agreement (1 out of 307 mutations in 2:2 segments with different CCF). In Supplementary Figure S17, instead, we show a case where CNAqc detects uncertainty in 14% of input triploid mutations, informing of potential challenges in using CCFs for those mutations. In that case the uncertainty is explained by the intermixing between two clonal picks in triploid 2:1 segments. Ccube assigns multiplicity 2 to a group of clonal SNVs at the right tail of the lowest clonal pick. The consequent CCF distribution breaks the expected clonal peak around ~1, alluding to the presence of two close CCF clusters. This is due to Ccube assigning some single-copy mutations $m = 2$, and vice versa. The entropy-based method by CNAqc highlights 14% of 2:1 mutations as uncertain, including the ones mistaken by Ccube. In turn, CNAqc assigns a FAIL status to these mutations with default values (cutoff >10%). Notably, the CCF distribution returned by CNAqc, which uses 86% of total mutations once the 14% unassignable are removed, is correctly peaked at ~1.

Errors in CCFs can affect downstream subclonal deconvolution, which in turn inflates evolutionary statistics (e.g., number of subclonal clusters, clonal complexity). In this example, miscalled multiplicities generate a spurious cluster in the CCF distribution fit by Ccube, which leads to subclonal cluster 2 (panel g, Supplementary Figure S17). Even after removal of 14% CCFs flagged as uncertain by CNAqc, Ccube still assigns the wrong mutation multiplicity to a significant number of variants and infers the spurious CCF cluster (panel h, Supplementary Figure S17). For this reason, reporting a FAIL status in CNAqc informs that multiplicity computation in this sample is highly confounded by intermixing of VAFs, cautioning the interpretation of downstream deconvolution analyses.

**Wall-time performance against deconvolution methods.** In order to understand how performance scales with sample size, we compared the wall-clock time of CNAqc against common deconvolution tools. We chose sciClone [14], Ccube [34] and Pyclone-vi [60] to represent a diverse set of popular algorithms for deconvolution. To build the dataset we subsetted all the mutations in diploid regions from a melanoma sample of the PCAWG cohort (patient id DO220877) leading to a total of 207508 mutations. This is the PCAWG sample with highest mutational burden in the cohort. Then, we sampled $N = \{500, 1000, 25000, 5000, 1000, 25000, 50000\}$ SNVs; this process was repeated 10 times to have 10 replicates for each N. The CNAqc analysis for peak detection was run with default parameters. Similarly, default parameters were also used for Sciclone (default one-dimensional deconvolution) and Ccube (but with numOfRepeat=1); Pyclone-vi was run with beta binomial likelihood, number of clusters from 1-10 and 30 repetitions (Supplementary Figure S5).

CNAqc was the fastest tool, capable of processing up to 500,000 mutations in ~60 seconds, while tools based on variational inference were about an order of magnitude slower. The latter two algorithms ranged from being 4 to 16 times slower than CNAqc for our range of tests (consider the log-scale in the plot y-axis), and the performance gap increased with larger N. Notably, sciClone took an average of two hours to process 50,000 mutations, which is 128 times slower than CNAqc as suggested by a log-difference of 5. In all tests, CNAqc, Ccube and Pyclone-vi scaled approximately exponentially, while Sciclone showed a jump from 25,000 to 50,000 mutations. All simulations were performed on a machine with 36 Intel(R) Xeon(R) Gold 6140 CPUs @ 2.30GHz and 220 GB of RAM (Ubuntu 20.04 LTS, Python 3.8.2 and R 4.1.0).

#### Analysis of patient data

##### PCAWG  WGS data

We performed QC of the entire Pan-Cancer Analysis of Whole Genomes (PCAWG) cohort [28]. First, we collected the sample list (file pcawg_sample_sheet.tsv at https://dcc.icgc.org/releases) with 2955 identifiers from PCAWG (2834 unique donors), and removed samples for which consensus CNAs were unavailable, identifying 2778 (2658 unique donors) WGS samples from 20 primary sites in 48 distinct projects. Then, we filtered samples that did not carry at least 20 mutations on each of the called copy number configurations, considering simple clonal segments, to compile our final list of working samples. We note that among the samples we did analyse and that passed QC, there were 75 samples that, for distinct reasons, were originally greylisted by the consortium. We report all the samples that we included, excluded a-priori (no consensus copy number calls available) and filtered (less than 20 mutations) in Supplementary Table S1.

Results for the analysis for simple and complex CNAs are shown in Figure 5 and Supplementary Figure S28. Tumour types with higher prevalence of complex CNAs are esophageal adenocarcinoma (ESAD), liver cancer (LIRI), melanoma (MELA),  ovarian cancer (OV), pancreatic cancer (PACA) and breast cancer (BRCA); liver and pancreatic cancers account for the majority of the samples of the cohort. Among these CNAs, those with ploidy beyond 6 are rare and more frequent in those tumour types with a higher number of CNAs. One may argue that this pattern might be linked to a general tendency of those tumour types to acquire these kinds of anomalies. The mean matched peaks for segments with ploidy $\leq 6$ is above 50% in most cases, and tends to be lower for segments with higher ploidy (increasing again in the final part of the spectrum), a phenomenon that could be due to the lack of an evolution-based QC for complex CNAs in our tool.

Results for subclonal CNAs are shown in Supplementary Figures S19 and S20. The top 5 tumour types carrying the most subclonal CNAs are: esophageal adenocarcinoma (ESAD), liver cancer (LIRI), melanoma (MELA), pancreatic cancer (PACA), breast cancer (BRCA). Four out of five of these tumour types also carry clonal complex copy numbers, supporting the hypothesis of some biological mechanism involved in CN instability in these types of tumours. To match peaks in subclonal CNAs, we computed all possible evolutionary models from a 1:1 starting state. We determined which model (linear or branching) best explains the data from the highest percentage of matched peaks; in case of a tie an “ambiguous” flag was assigned and, if no model could match at least 50% of peaks, “none” was assigned. By imposing this restriction, we were able to assign >87% of segments to a model. The first thing we notice from Supplementary Figure S21 is that the most frequent subclonal CNA is either the loss or the acquisition of a single allele, corresponding to 1:0-1:1 and 1:1-2:1. This is not surprising as these events are explainable with a deletion of a single allele in one cell duplication. Other CNA combinations must instead be explained by at least two independent losses or duplication events, making them less likely. The less frequent combination is 1:0-2:0, possibly due to the fact that a duplication and a deletion event occurring on the same allele seem unlikely, unless the tumour has a high instability and predisposition to achieve CNLOH. The same pattern repeats across tumour types, with tumours tending to have more simple subclonal events also showing a higher number of complex events, supporting the hypothesis that some tumour types are more prone to develop CNAs [5,17].

In most cases, a linear modelling of the dynamics of the formation of the subclones (meaning that the second subclone arises from the first), seem to better explain the data with respect to a branching modelling in which both subclones independently originate from a diploid ancestor. There are two exceptions to this statement; first, for subclones with 1:0-1:1 karyotypes (i.e., a 1:0 subclone and a 1:1 subclone), where there is no difference in the peaks expected by the linear and branching models; therefore the models are indistinguishable. This is evident from Supplementary Figure S21b and S21c, where in all tumours the number of matched peaks for this combination of subclonal karyotypes is equivalent. The second exception is for 2:0-2:1 subclones, in which the branching model seems to explain the data better. The reason for this might be that, while both models require at least three steps for the two subclones to develop the karyotype 2:0-2:1 from a diploid 1:1 ancestor, the branching model can take into account both the path in which the major allele is the same in both clones, or is the major in one clone is the minor in the other. The linear model's shortest path, followed by assumption in CNAqc, can only include the first of the two scenarios.

##### Genomics England WGS data

We further validated CNAqc by performing control of 235 samples obtained from the Genomics England Consortium [19] using the Illumina DRAGEN™ pipeline. We started from 301 patients for which we had complete DRAGEN™ calls available and removed those belonging to tumour types with less than 10 samples associated. Since by default the tool can identify putatively heterogeneous regions but does not give an estimate of the number and prevalence of subclonal populations we had to derive those quantities using some heuristics. We developed a simple procedure to estimate CCFs by assuming the presence of two subclones, following the same heuristics implemented in the popular tool BATTENBERG [15]. Without loss of generality we will define the integer solution of DRAGEN™ in terms of minor and total copy number as subclone 1.

We started from the floating point estimates for:

- the minor allele (mCNF);
- the total copy number (CNF);
- minor allele frequency (MAF);

and determined:

- integer minor and total allele copy number for subclone 1 ($mCN_1$ and $CN_1$);
- integer minor and total allele copy number for subclone 2 ($mCN_2$ and $CN_2$)

We proceed as follows:

1. Set a grid of values for $mCN_2$, default 0 to 10;
2. Estimate CCF of the segment using the formula

$$\mathrm{CCF} = \frac{\mathrm{mCNF} - \mathrm{mCN}_1}{\mathrm{mCN}_2 - \mathrm{mCN}_1}$$

3. Estimate $CN_2$ by

$$\mathrm{CN}_2 = \frac{\mathrm{CNF} - (1 - \mathrm{CCF})\mathrm{CN}_1}{\mathrm{CCF}}$$

4. Calculate the MAF error

$$\mathrm{MAF}_{\mathrm{err}} = \left|\mathrm{MAF} - \pi\left[\mathrm{CCF}\frac{\mathrm{mCN}_1}{\mathrm{CN}_1} - (1 - \mathrm{CCF})\frac{\mathrm{mCN}_2}{\mathrm{CN}_2}\right] - (1 - \pi)\frac{1}{2}\right|$$

5. Choose the $mCN_2$ with the lowest MAF error $MAF_{err}$
6. Filtered negative solutions where $CN_2 < 2mCN_2$ as well as solutions where $MAF_{err} < \epsilon$ , with $\epsilon = 0.1$ (default threshold)

The intuition of this procedure is as follows. Given the assumption of a two subclone population and a maximum $mCN_2$ of 10 we need just $mCN_1$ to determine the $CCF$ value using the formula at point 2, together with the $mCNF$. To provide an example let us assume a segment with $mCN_1$ value of 1, a $mCNF$ of 1.8, a $CN_1 = 3$, a $CNF = 4.2$ and, $MAF = 0.4$ we will test $mCN_2 = [2,3,4,5,6]$ and assume purity $\pi = 1$ for simplicity (note that we skipped 1 as it implies a division by zero). For the 5 values of $mCN_2$ we get respectively $CCF$ values of [0.8,0.4,0.26, 0.2, 0.16], by applying the formula at step 3 we estimate $CN_2$, which in this case is [5,7,8,9,11]. We can clearly see how some solutions do not make sense (2$mCN > CN$) and get filtered in step 6. The estimated MAF errors in this case are [0.05, 0.01, 0.05, 0.11, 0.12], so we will chose the one with the lowest error, namely $mCN_2$= 3 and 7, this setting corresponds to two subclones one at frequency 80% with karyotype 1:2 and one at 20% with karyotype 3:4. As the absolute error is lower than 0.1 we accept the solution.

Upon converting DRAGEN™ continuous estimates into clone-level CNAs, we set all the parameters as in the analysis of PCAWG to allow for a fair comparison among Genomics England and PCAWG cohorts.

#### TCGA WES data

We collected WES data from $n = 48$ lung adenocarcinoma samples available in TCGA LUAD [29], selecting the 24 ones with top and bottom consensus purity estimate (CPE) by TCGA. We report example cases in Supplementary Figure 23, where QC values are obtained by using somatic SNVs, CPE purity and default CNAqc parameters. The case in panel (a), sample TCGA-53-7624-01A, is 84% pure and the inferred ploidy is correct, but purity is slightly overestimated. The case in panel (b) is 82% pure, but with a similar pattern. The case in panel (c) is an example of a VAF distribution that is low resolution because the sample has 30% purity, and in this case it is difficult to assess if the small peak matched by CNAqc is a noise artefact. The case in panel (d) is 83% pure, with good calls and the one in panel (e) is 32% pure and passed because most of the tetraploid mutations are correct, but it contains a poorly-peaked VAF distribution in triploid states (2:1, 47% of the mutational burden). In this last case CNAqc struggles to detect peaks from VAF; this is another example of low resolution VAF distribution.

We used CNAqc to select among multiple purity estimates provided by different TCGA callers, focusing on the LUAD case (a) from Supplementary Figure S23. In TCGA, we obtain purity estimates from CPE, which is the consensus among ABSOLUTE, ESTIMATE, IHC and LUMP. For this sample, ESTIMATE, IHC and LUMP provide similar purity and determine the value for CPE. However, we fail that estimate with CNAqc and instead pass only ABSOLUTE (69% purity, Supplementary Figure S24). We extended this test to 1464 TCGA samples from 10 distinct tumour types (Supplementary Figure S25 and S28) with suitable data for CNAqc (i.e., at least 200 somatic mutations). We obtained samples form the cohorts Bladder Urothelial Carcinoma (BLCA), Breast Invasive Carcinoma (BRCA), Colorectal Adenocarcinoma (COAD), Glioblastoma (GBM), Head-Neck Squamous Cell Carcinoma (HNSC), Kidney Renal Clear Cell Carcinoma (KIRC), Lung Squamous Cell Carcinoma (LUSC), Rectum Adenocarcinoma (READ) and Uterine Corpus Endometrial Carcinoma (UCEC).

First, we computed QC with CNAqc with maximum tolerated purity error 5% ($\epsilon = 0.05$) for all possible purity values for the ABSOLUTE, ESTIMATE, IHC and LUMP tools, as well as for consensus CPE purity. We report in panel (a) of Supplementary Figure S25 cases split by QC status (maximum tolerated purity error 5%) as determined from the run with TCGA consensus purity. Strikingly, as in Supplementary Figure S24, we immediately observe a number of cases in which, while CNAqc fails the CPE estimate, there are at least $\geq 1$ method different from CPE that proposes an acceptable purity. Notably, for 901 cases where the CPE purity is failed by CNAqc (60% of 1464), upon splitting the status by tool in panel (b), we note for instance that ABSOLUTE often provides a purity estimate that would pass the sample, similarly to the case shown in Supplementary Figure S24. This is particularly true for samples from the HNSC, LUAD, COAD, LUSC and BRCA cohorts. Conversely, methods such as ESTIMATE very often provide failed purity estimates for the samples (especially from the HNSC, LUAD, LUSC and BRCA cohorts). If we were to rank and select the best purity as determined by CNAqc instead of using consensus estimates reported by TCGA, 785 out of 901 cases (~88%) would be rescued, avoiding using a consensus purity that contains an error that is larger than 5%. Overall, this shows that CNAqc can be used to select among multiple purity estimates even from WES, avoiding at least in principle the need of consensus calling.

##### Multi-region colorectal cancer WGS

A common design of modern cancer genomics is to collect multiple, spatially-separated, samples from the same tumour. We have sought out to test CNAqc on previously published WGS multi-region data of primary colorectal adenocarcinomas [30,53]. We gathered data for 2 patients, for a total of 10 samples with median coverage ~80x, purity ~80% (Figure 6); for these samples we generated mutations and CNAs with Platypus [53] and Sequenza [22]. Since Sequenza can return multiple solutions to the CNA-inference problem, we tested if  CNAqc could select the best tumour segmentation as compared to published calls obtained by the CloneHD [26] algorithm.

Sequenza with default parameters returned a main solution close to CloneHD, proposing an alternative tetraploid solution with halved purity. We used it to re-run Sequenza, and also generated another low-purity alternative solution. We used CNAq to compare these 3 runs; for sample Set7_57 from patient Set7 (Figure 6a) CNAqc selected the correct diploid solution with 80% purity, suggesting only a small purity adjustment that did not change the quality of the QC (Figure 6b, 6c). Interestingly, peak detection scores from CNAqc invariably failed both the tetraploid and low-purity solutions (Figure 6d, 6e), showing how CNAqc can be used to select among alternative solutions proposed by a copy number caller.  An equivalent result was also obtained for 6 WGS samples of patient Set_6 (Supplementary Figure S30).

#### Supplementary Figures

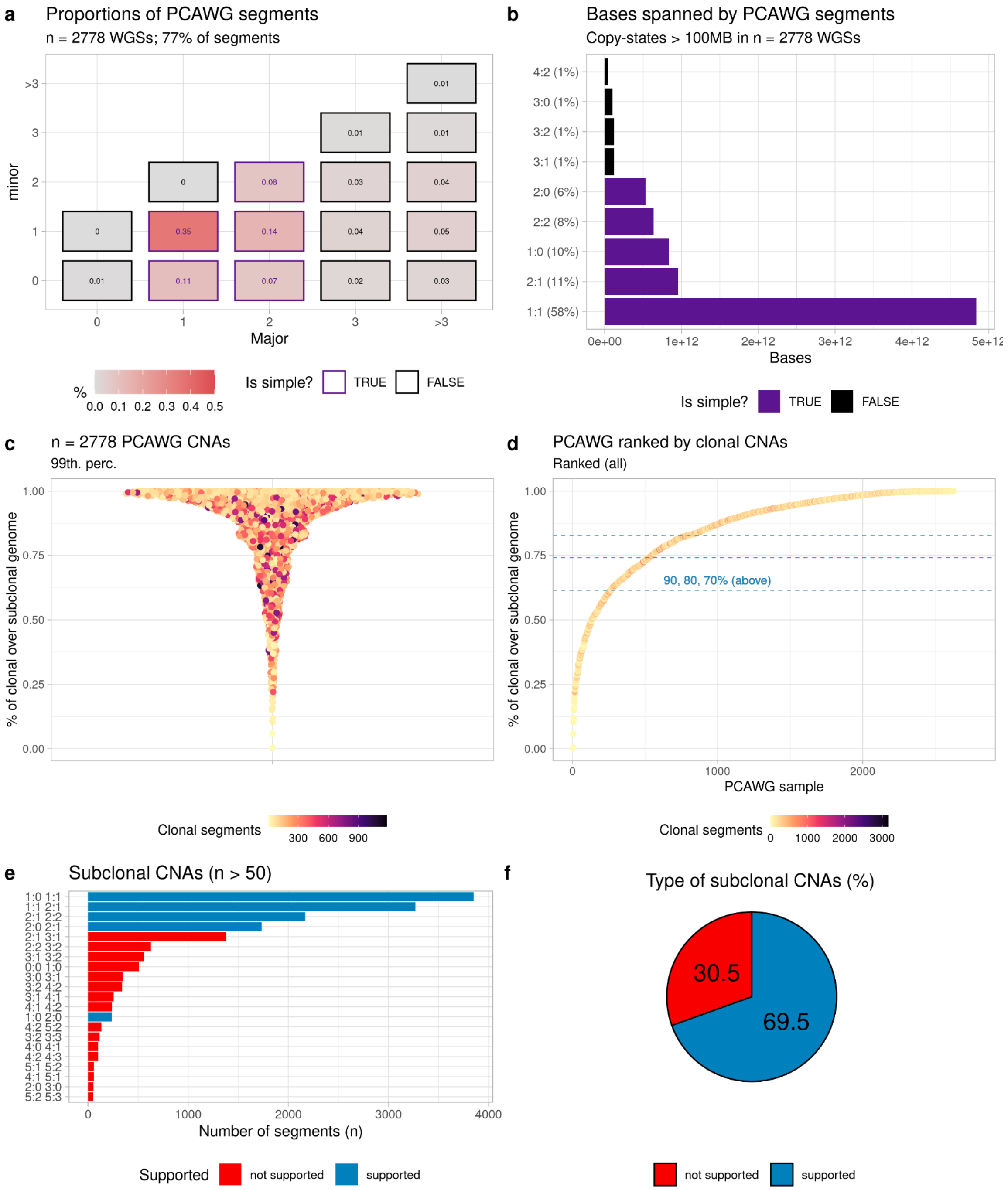


**Supplementary Figure S1. a.** Proportion of PCAWG CNA segments split by copy state, obtained by consensus calling across multiple callers with $n = 2778$ WGS samples of primary tumours. The matrix reports major and minor alleles, the colour and number reflects the proportion of CNA segments with that copy state across total (e.g., 36% of segments are diploid heterozygous, 1:1). CNA segments used by CNAqc are coloured in purple; in total, 78% of the overall set of segments (>600,000) can be processed by our method (36% of segments are 1:1, 15% are 2:1, 11% are 1:0, 8% are 2:2 and 8% are 2:0). **b.** Number of bases covered, and proportions relative to the total genome spanned by all the PCAWG segments in panel (a). Diploid heterozygous segments cover over a thousand billion bases( $> 10^{12}$), accounting for 58% of the genome covered by these segments. The segments supported by CNAqc are the top-5 most common segments reported across all PCAWG, covering 93% of all bases sequenced in this cohort. **c.** Battenberg clonal and subclonal CNAs available in PCAWG. To simplify the visualisation we remove outliers exceeding the 99-th quantile of the data distribution. Every dot is the percentage of the tumour genome spanned by clonal segments, coloured by the number of segments per sample. So if a sample has >50% of clonal segments it is above the horizontal dashed line. **d.** We rank by sorting the percentages shown in panel (c) to note that only $n = 124$ PCAWG samples (vertical dashed red line) have more subclonal than clonal CNAs. **e.** Number of CNA segments covered by subclonal copy number events, divided by karyotype. The plot shows how the most common subclonal events involving "simple" karyotypes are supported by CNAqc. As in panels (c and d), we consider a segment subclonal if the CCF provided by Battenberg is lower than 1, and clonal otherwise. As such subclonal structure is limited to 2 clones **f.** Percentage of subclonal segments supported by CNAqc. The number calculation is done on the number shown in panel (e).

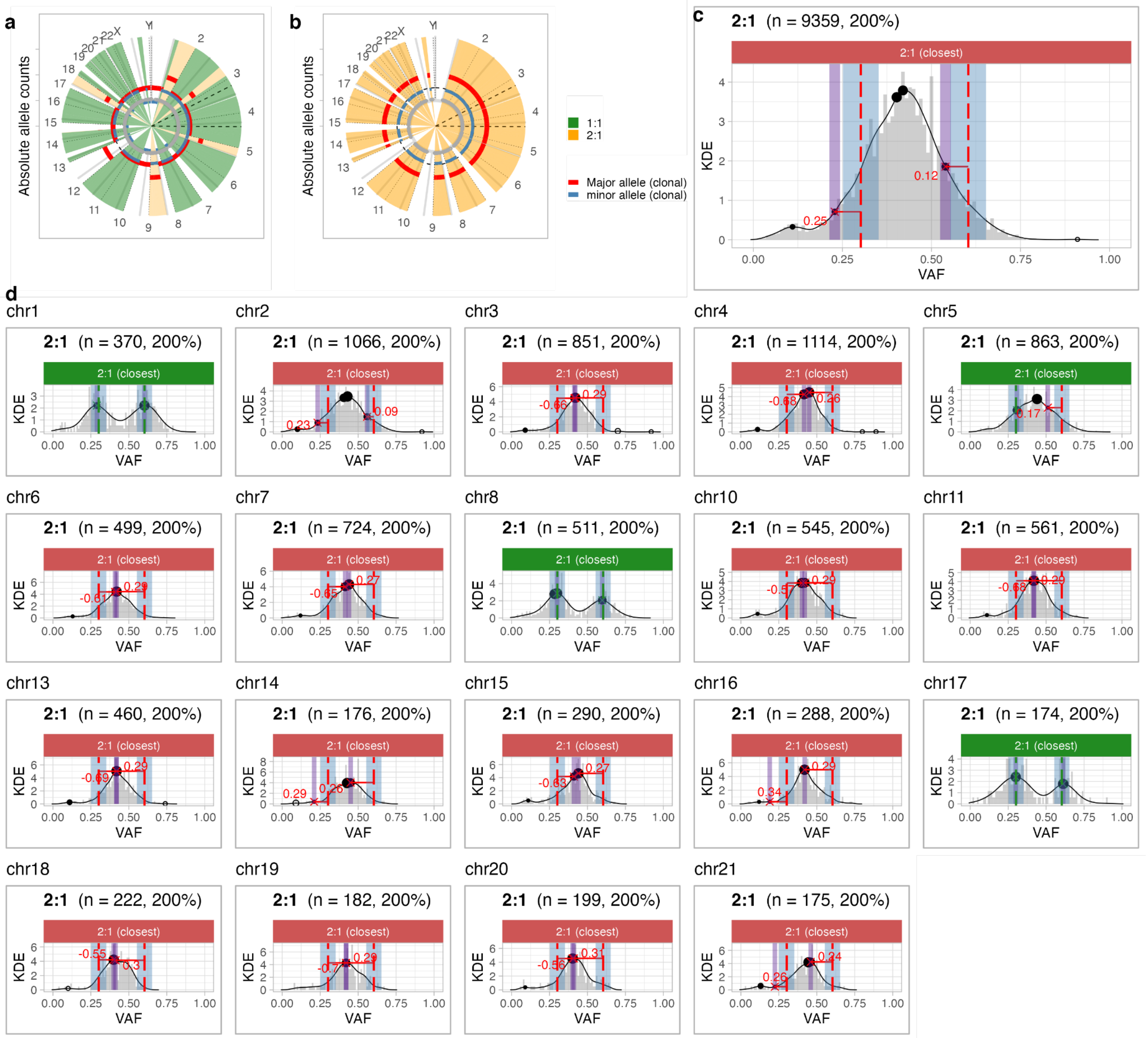


**Supplementary Figure S2. a-b.** Chromosome-level analysis of an artificially miscalled sample, obtained from the PCAWG hepatocellular carcinoma sample ca5ded1c-c622-11e3-bf01-24c6515278c0. From (a) the true copy-number profile called on this sample we simulate (b) a wrong profile in which the whole genome is called triploid. **c.** The incorrect calls are detected by the peak analysis performed by CNAqc on the whole genome, which correctly fails the test and suggests a correction on the sample purity estimate. **d.** By peak analyses at the level of individual chromosomes it is easy to identify those portions of the genome whose copy state has been incorrectly called triploid.

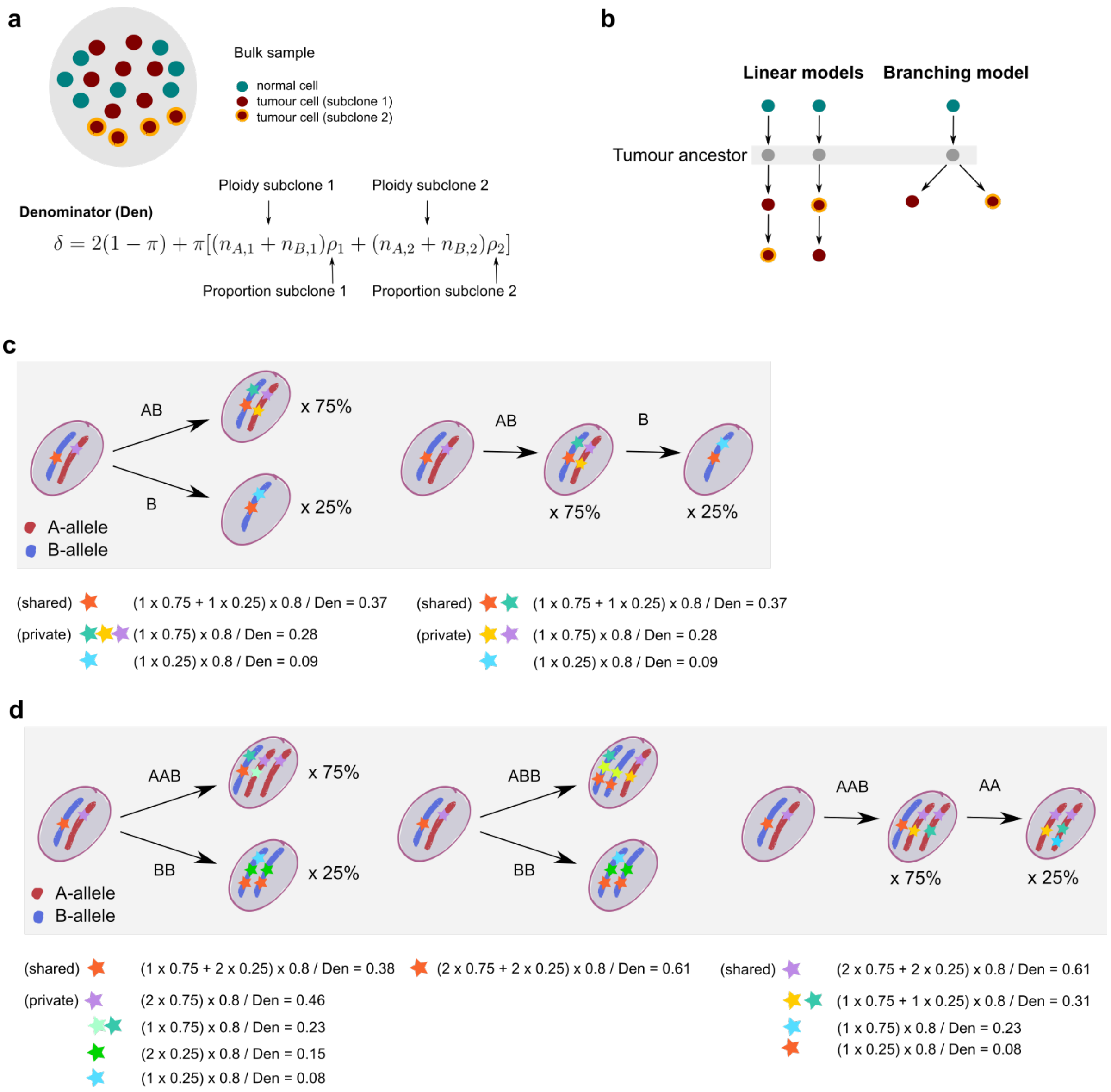


**Supplementary Figure S3. a.** Tumour bulk sample composed of both normal and tumour cells belonging to two distinct subclones. In order to compute all possible expected VAF peaks, one needs to account for the multiplicities of the variants and weight them according to the proportion of normal reads $(2(1-\pi))$ and reads coming from the first and the second subclone $\pi[(n_{A1}+n_{Bi})\rho_1 + (n_{A1}+n_{B2})\rho_2]$. **b.** Linear and branching models representing the possible evolutionary paths explaining the formation of the observed subclones. **c.** Linear and branching model for subclones 1:0 and 1:1 with ancestral diploid state, and computation of the expected VAF peaks. Although the variant groups are different between the two evolutionary paths, it is not possible to distinguish them from data as the expected VAF peaks are the same. **d.** Like in panel (c) but for subclones 2:0 and 2:1. In this case, both the number of peaks and their expected position change according to the sequence of events that led to the formation of the subclones.

Note the difference in the VAF peak for shared mutations in two possible branching models (AAB | BB and ABB | BB).

**Algorithm:** ***Peak detection in CNAqc***

**Input:** Mutations, allele-specific CNA segments, purity,

**Parameters:** purity error tolerance $\varepsilon > 0$ and VAF tolerance $\varepsilon_{VAF} > 0$

---

- set $K = \{1:0,\ 1:1,\ 2:0,\ 2:1,\ 2:2\}$, where $n_A : n_B$ are the copies of the Major/ minor alleles;

*# Peak detection for every copy state*

- for every copy state $k \in K$ and multiplicity $m \in \{1,2\}$:
  - retain mutations $M_k$ mapping to segments with copy state $k$;
  - compute $v_m$ with equation (1);
  - compute $\varepsilon_m$ with equation (3);
  - determine peaks $d_1, ..., d_n$ from the VAF distribution of $M_k$ ;
  - match $d_*^m$ to $v_m$ by either closest or rightmost hit;
  - define interval $I_m$ from $v_m$ and $\varepsilon_m$ with equation (6);
  - define $I_m^{VAF}$ from $d_*^m$ and $\varepsilon_{VAF}$ with equation (6);
  - define PASS for $k$ and $m$ if $|I_m^{VAF} \cap I_m| > 0$, FAIL otherwise;
  - determine the number $n_m$ of mutations mapping below VAF peak $d_*^m$;
  - if $k \in \{2:0,\ 2:1,\ 2:2\}$, compare $n_m$ across peaks and define the status for $k$ from the largest $n_m$, otherwise use the only available peak;

*# Sample level quality control status*

- for all copy states $k \in K$, define $w_k$ by normalising the number of mutations mapped to the copy state, and rescale $w_k$ by 2 if $k \in \{2:0,\ 2:1,\ 2:2\}$;
- for every copy state $k \in K$ define $\lambda_{k,}^{PASS}$ and $\lambda_{k,}^{FAIL}$ with equations (7) and (8);
- define the sample score $\lambda$ with equation (9), and evaluate the sample status from the sum of $\lambda_{k,}^{PASS}$ and $\lambda_{k,}^{FAIL}$ for all $k \in K$, taking the largest.

**Supplementary Figure S4.** Pseudocode of the peak detection algorithm and quality control strategy in CNAqc (Online Methods) that apply to simple clonal CNAs in order to determine purity adjustments for a copy number caller, and sample level QC metrics.

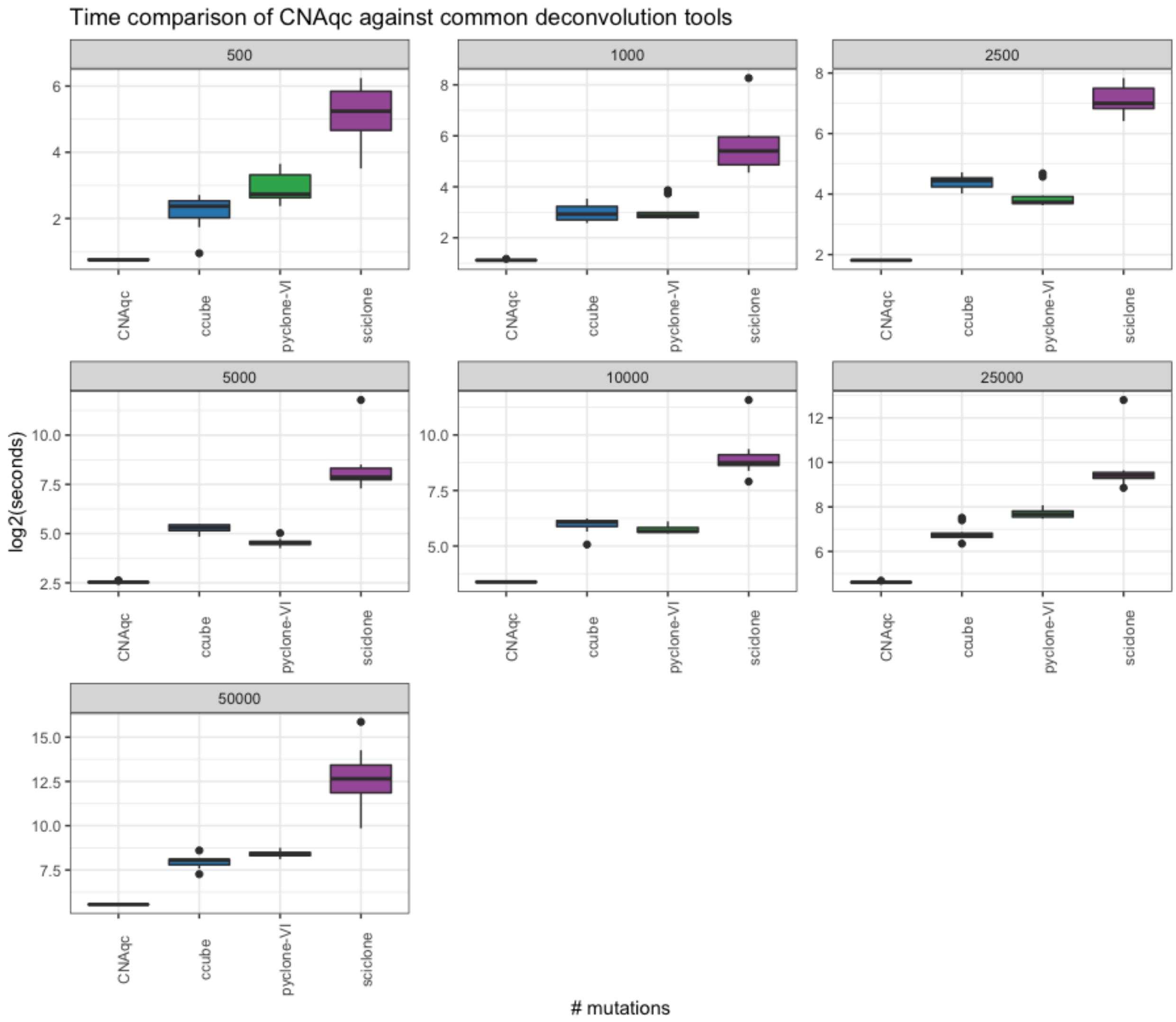


**Supplementary Figure S5.** Wall-clock time of CNAqc, compared with common subclonal deconvolution tools (Ccube, Pyclone-VI and Sciclone) on datasets with 500, 1000, 2500, 5000, 10000, 25000 rr 50000 mutations. The CNAqc peak detection algorithm is extremely fast and preprocesses even 50000 mutations in less than a minute (~47 seconds). The fastest deconvolution tools are Pyclone-VI and Ccube, both implemented using Variational Inference; Sciclone drops rapidly in performance as the number of SNVs increases. Time is reported in log2(seconds).

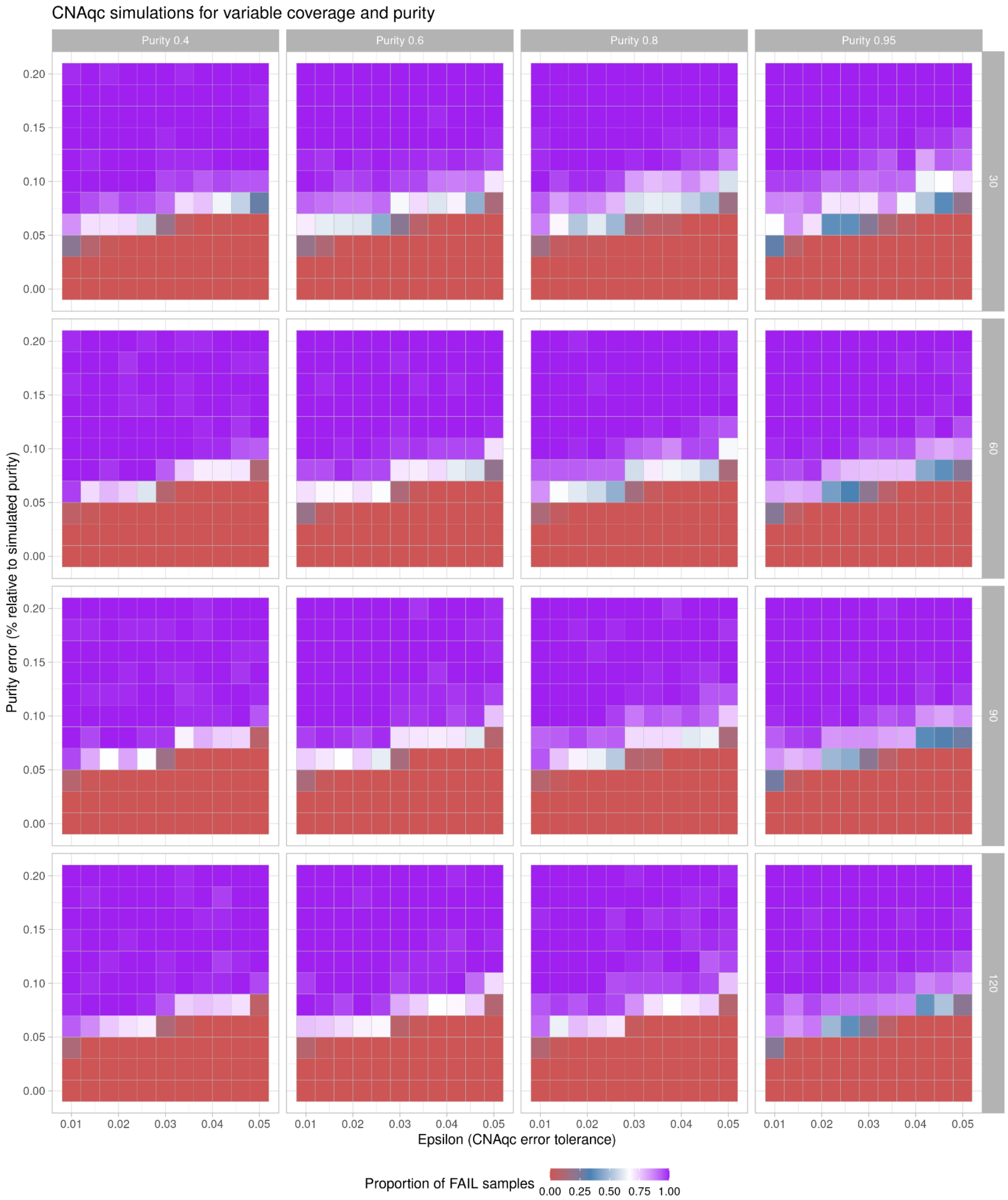


**Supplementary Figure S6.** CNAqc tests on synthetic tumours generated with different coverage and purity. We report the proportion of rejected samples running the tools with an error on the simulated purity (y-axis), and a tolerance to match peaks (x-axis).

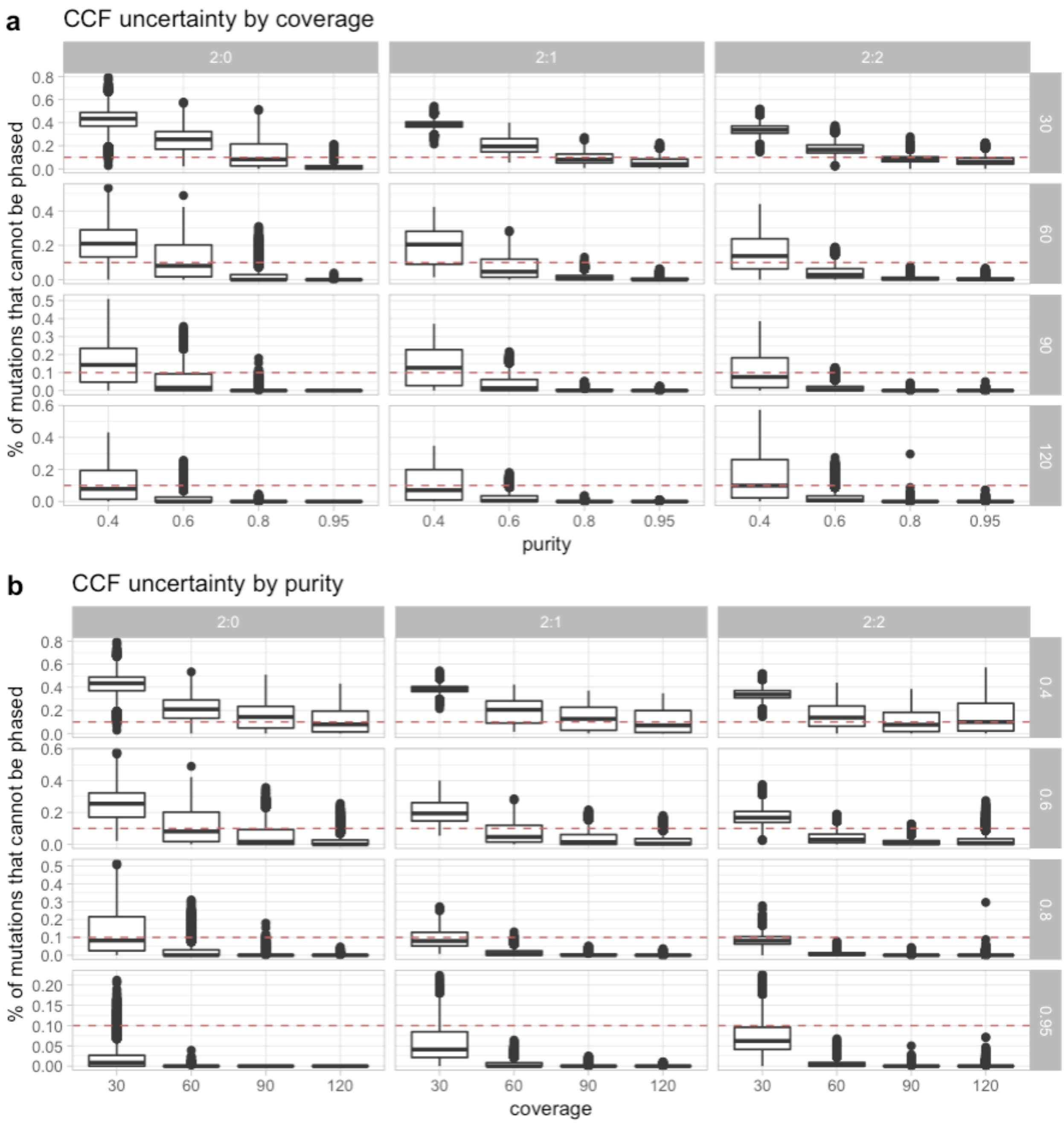


**Supplementary Figure S7. a.** For the simulated tumours in Supplementary Figure S5, we report the proportion of mutations for which CNAqc does assign a CCF (uncertain in phasing multiplicities), as a function of purity at fixed coverage values. The dashed line at 10% is the default parameter value to determine the final PASS or FAIL status per copy state. **b.** As in panel (a), but fixing purity.

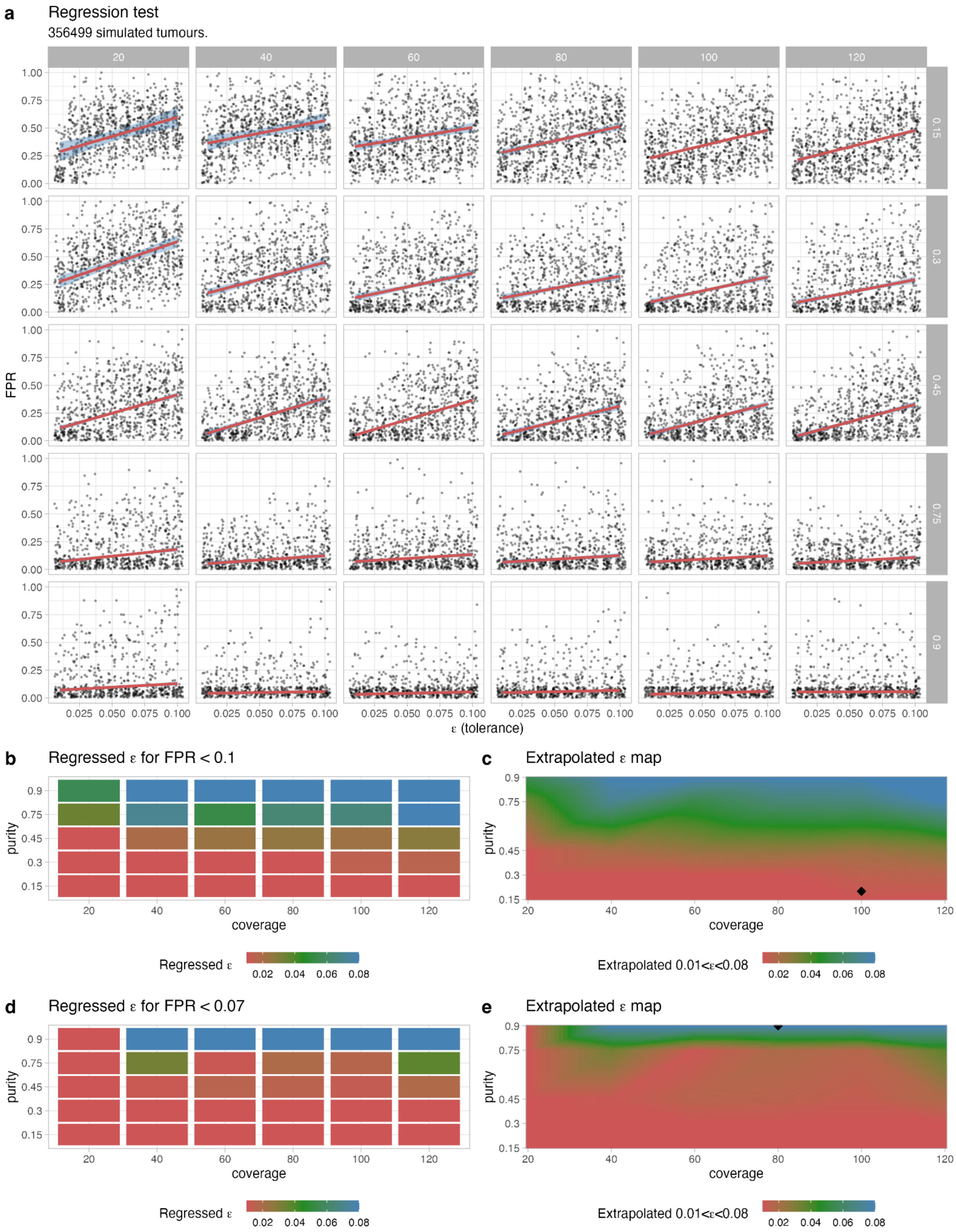


**Supplementary Figure S8. a.** Calibration test for CNAqc, trying to determine the best input $\epsilon$ as a function of purity and coverage with 100 distinct tumour segmentations, where for each $\epsilon$ in each panel we generate 10 datasets. To every dataset we apply a random uniform noise in the range $[\epsilon; \epsilon + 3\%]$, i.e., around the boundary of acceptability; we seek to determine if the tool, with $\epsilon$ in input, can flag the sample as FAIL. The panels show false positives rates (FPR) from PASS samples that should be instead failed, and regressed performance (red line). As expected, for low purity/coverage we observe higher FPR. **b,c.** For every input value of purity (passed by the user) and coverage (obtained from data), we invert the regression fit in panel (a) for a given value of FPR (target 10% in this panel). At this point, we can extrapolate $\epsilon$ to the full range of trained values; for a 100x sample with 20% purity one should run CNAqc with $\epsilon = 0.01$ in order to keep the FPR below 10%. This makes sense since at low purity noise in peak detection can confound the tool which would pass peaks that should not be passed. **d, e.** Same as in panels (b,c), with different parameters (FPR < 0.07; coverage 80x, purity 90%).

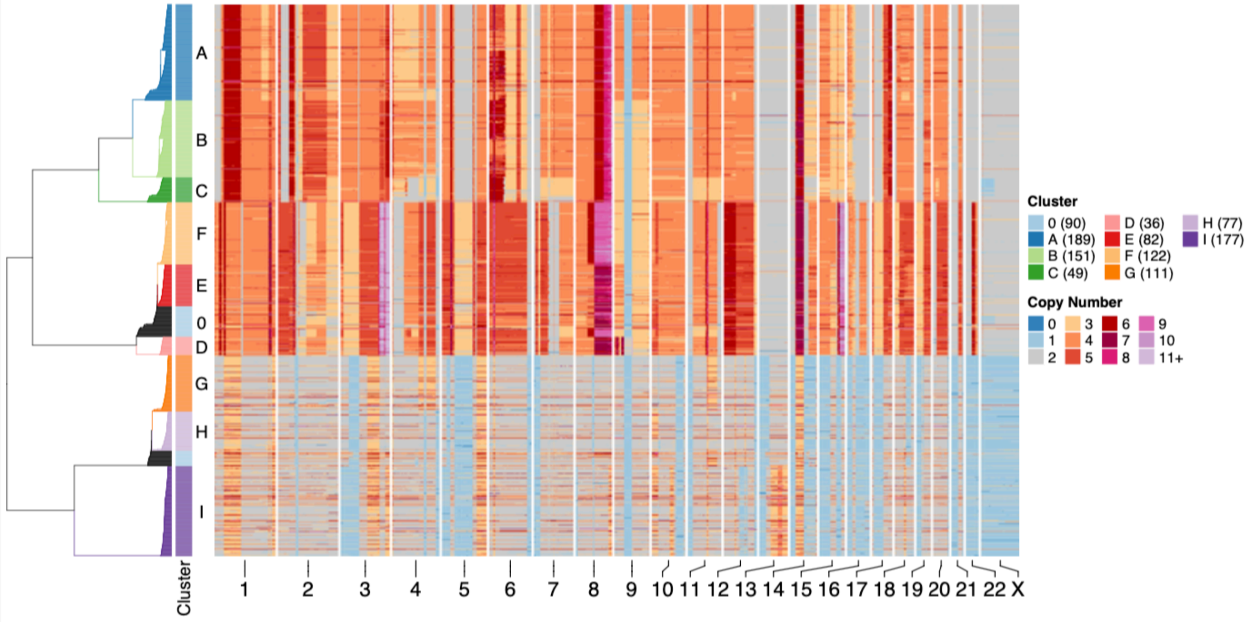


**Supplementary Figure S9.** Single-cell UMAP [61] clusters obtained from low-pass single-cell data of 1084 cells from an ovarian cancer assay. These data are generated by using the DLP+ library preparation protocol [31]. Upon calling of cell-level allele-specific CNAs, cells are clustered into several groups that define CNA-level clones.

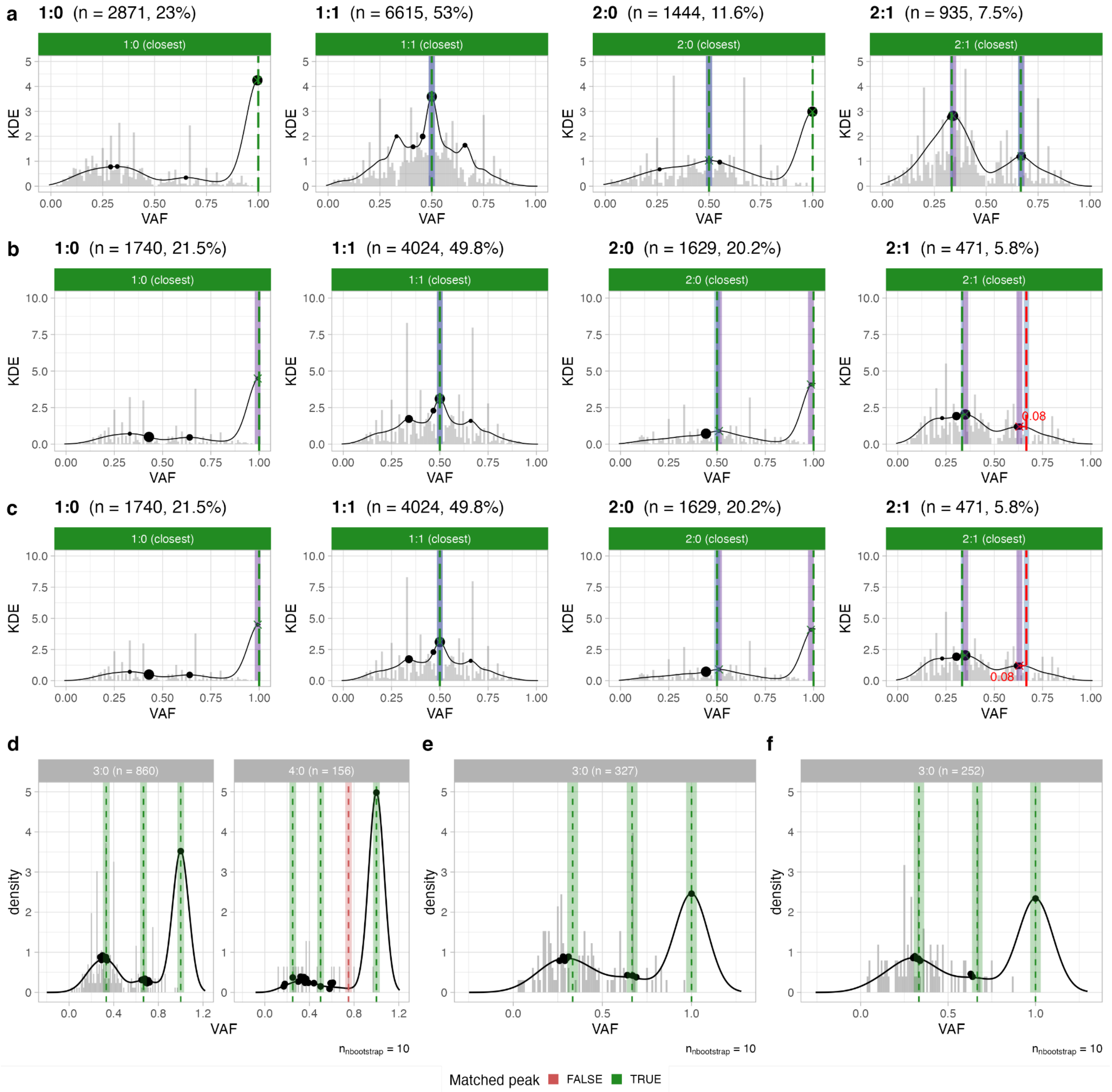


**Supplementary Figure S10. a-c.** Analysis of simple CNAs obtained from pseudobulk data for clusters I (177 cells) , G (111 cells) and H (77 cells) from Supplementary Figure S7. These tumours are 100%-pure populations by construction, and CNAqc can validate the generated calls at both the level of somatic mutations and copy numbers, despite the small number of mutations and noisy VAF profile from single-cell data. **d-f.** CNAqc peak analysis of more rare CNAs from the clones in panels (a-c); LOH calls with 3 or 4 total alleles are matched by CNAqc, therefore validating the peak detection algorithms in the tool.

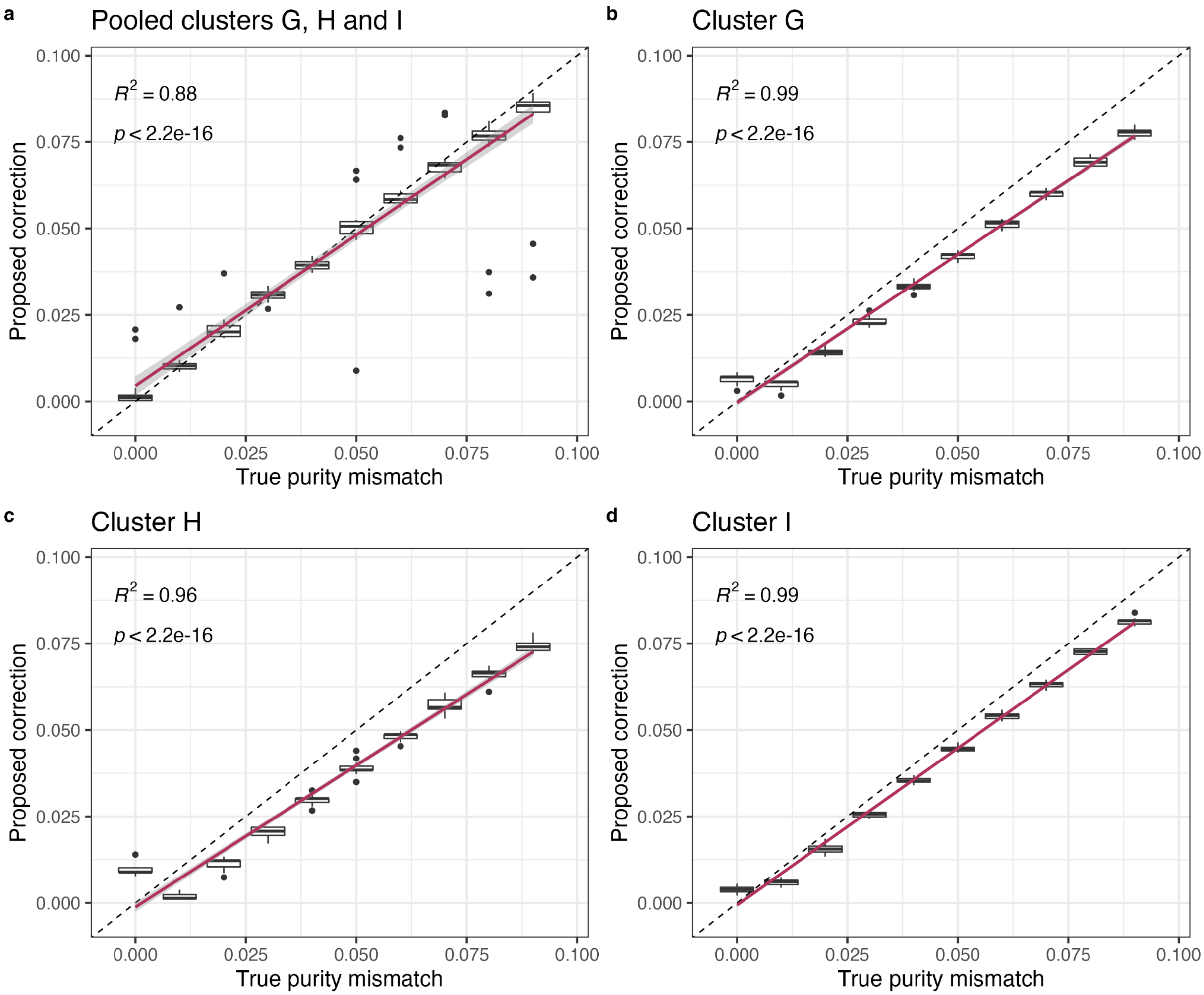


**Supplementary Figure S11. a-d.** Validating the accuracy of the correction on sample purity proposed by CNAqc using low-pass single cell data. The tests are performed from monoclonal pseudo-clones generated from clusters of single-cells with the same clonal copy number profiles, and restricted to genome regions with loss of heterozygosity, where the VAF profile is less noisy and each sample is 100% pure by construction. Peak analysis is run multiple times for each cluster, upon decreasing input sample purity from 100% to 90% with 1% step and 15 repetitions per point. The tested samples from Supplementary Figure S7 are (a) a pileup of clonal CNAs common to clusters G, H and I (also used in Supplementary Figure S10), in 1:0 and 2:0 regions, (b) cluster G restricted to 1:0 segments, (c) cluster H restricted to 1:0 segments and (d) cluster I restricted to 1:0 and 2:0 segments. In every case the proposed correction is in agreement with the input mismatch, the correlation coefficient is $0.88 < R^2 < 0.99$ with p-value $p < 2.2e-16$.

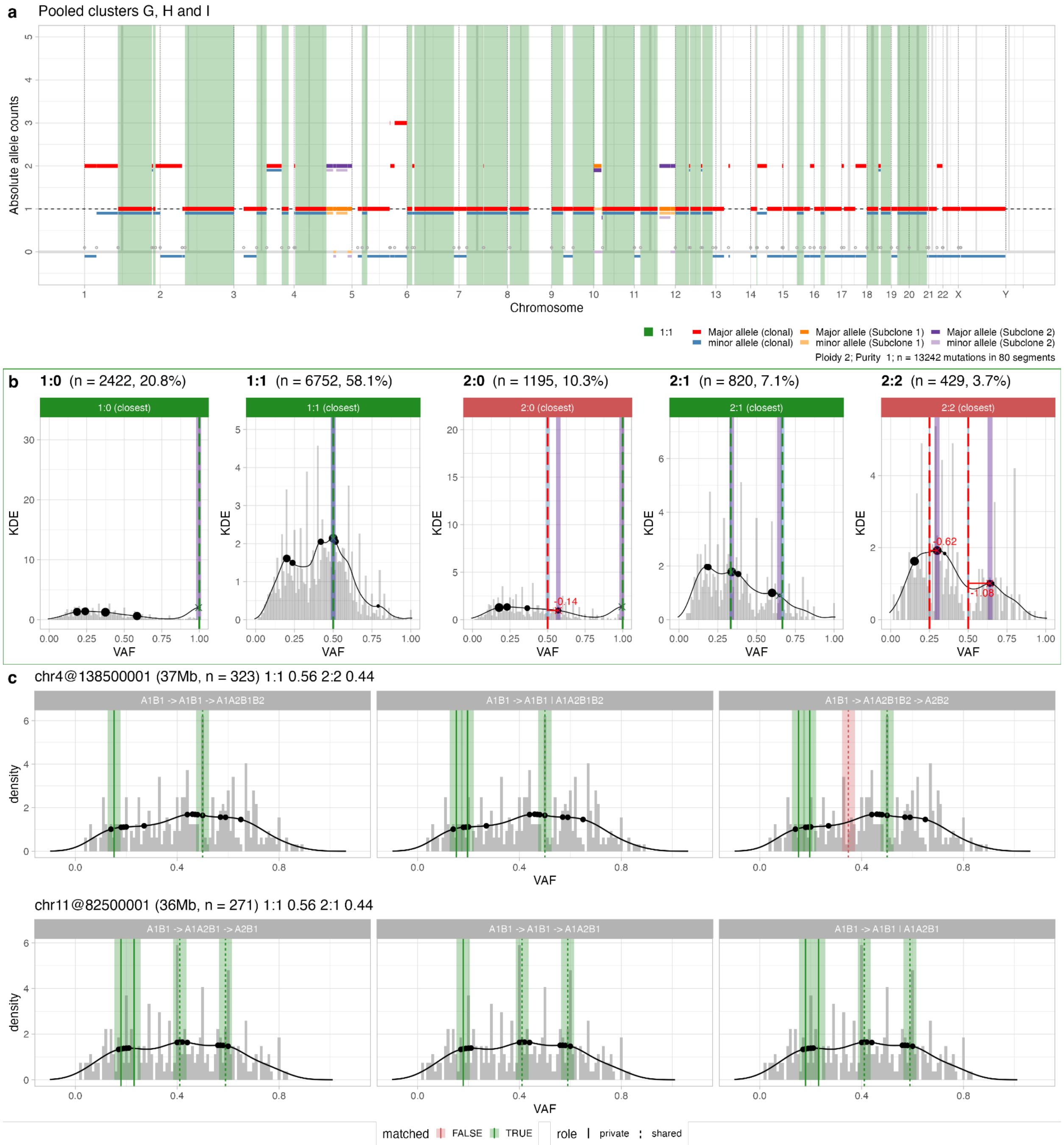


**Supplementary Figure S12 a.** Pseudo-clone generated by merging low-pass sequencing data for cells from clusters G, H and I in Supplementary Figure S7. Here, we retained clonal CNAs common to clusters G, H and I, and identified segments where clusters H and I have a CNA that differs from cluster G. Then, we mixed all the cells (111 for G, 77 for H and 177 for I), obtaining a mixture with ~70% cells from merged cluster H+I, and ~30% from cluster G. **b.** QC of the clonal CNAs for the sample, using purity 100%; CNAqc passess the sample. **c.** QC of 2 subclonal CNAs for the sample, harbouring >250 SNVs each. CNAqc validates the required VAF peaks, therefore supporting the presence of subclonal CNAs in these data. The model cannot distinguish branching versus linear dynamics because, in these VAFs, peaks are found - and matched - at very similar frequencies. For example the bottom panel A1B1→A1A2B1 | A2B1 denotes a branching ("|") model with 2:1 (A1A2B1) and 1:1 (A2B1) siblings, while A1B1→A1B1→A1A2B1 denotes a linear ("→") model with 1:1 (A1B1) ancestral to 2:1 (A1A2B1).

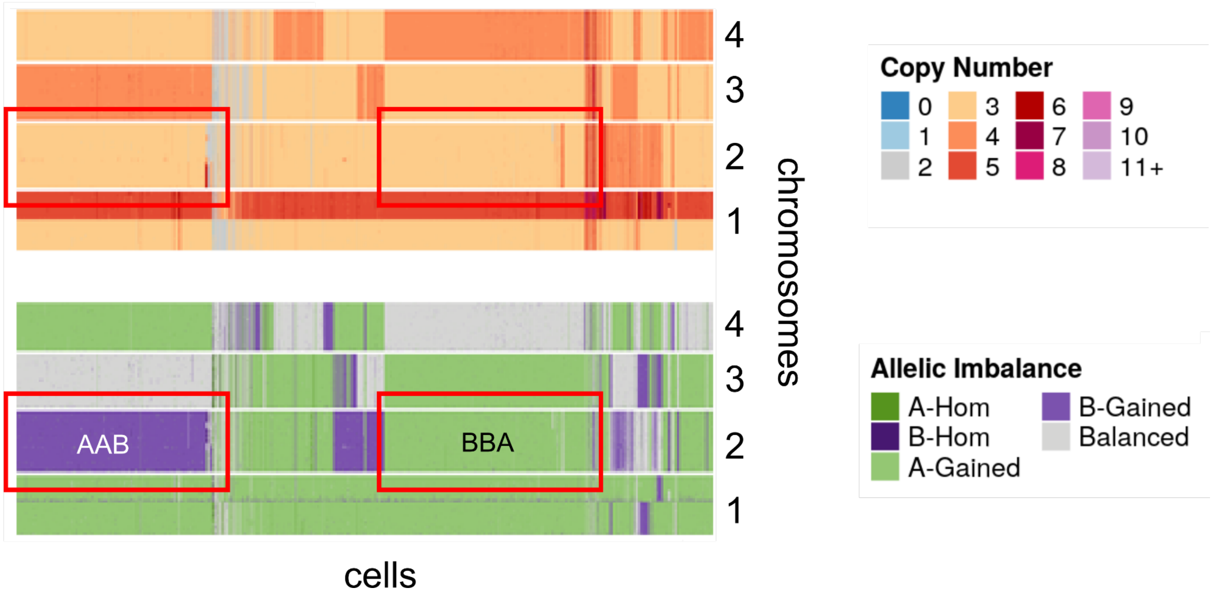


**Supplementary Figure S13.** Low-pass copy number data obtained with SIGNAL, from data released in [33]. The heatmaps show total copy number for chromosomes 1 to 4, as well as phased alleles. Note that chromosome 1 is monoclonal, whereas 2-4 are defined by 2 clones. On chromosomes 3 and 4 the populations are triploid and tetraploid; on chromosome 2 they are triploid with mirrored allelic imbalance (i.e., genotypes AAB and ABB).

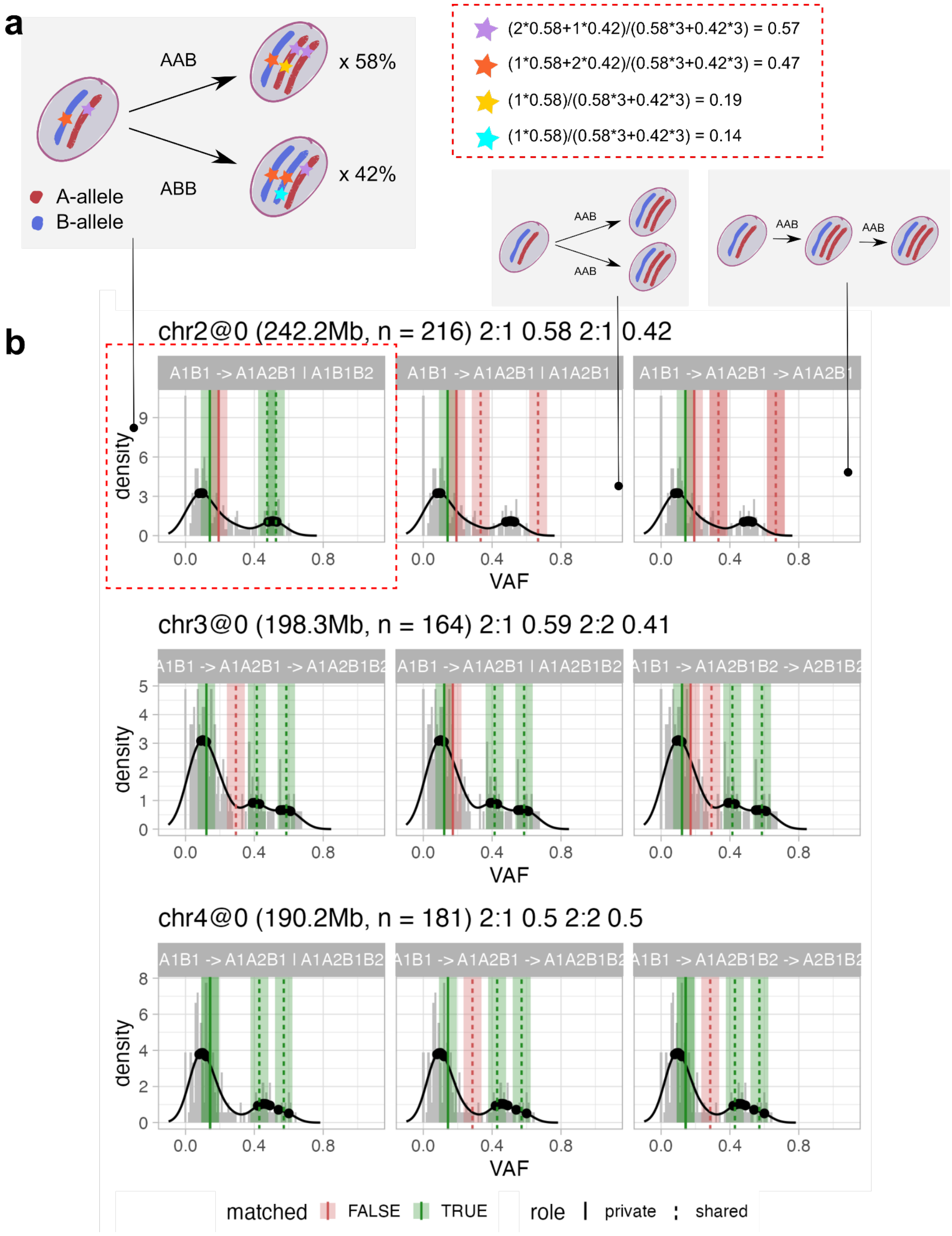


**Supplementary Figure S14. a.** The CNAqc evolutionary model for two 2:1 subclones (Supplementary Figure S11) with genotypes AAB and ABB and proportions 58%/42% predicts two peaks close to 50%, and two below 20%. **c.** From a pileup of data in Supplementary Figure S7 we perform validation of subclonal CNAs with CNAqc. The tool passes most peaks across all all the considered chromosomes, for a variety of possible linear and branching evolutionary models. Note that for chromosome 2 the tool can also identify the true evolutionary model where genotypes AAB and ABB (allelic imbalance), branch from an ancestral heterozygous diploid state (genotype AB, or 1:1).

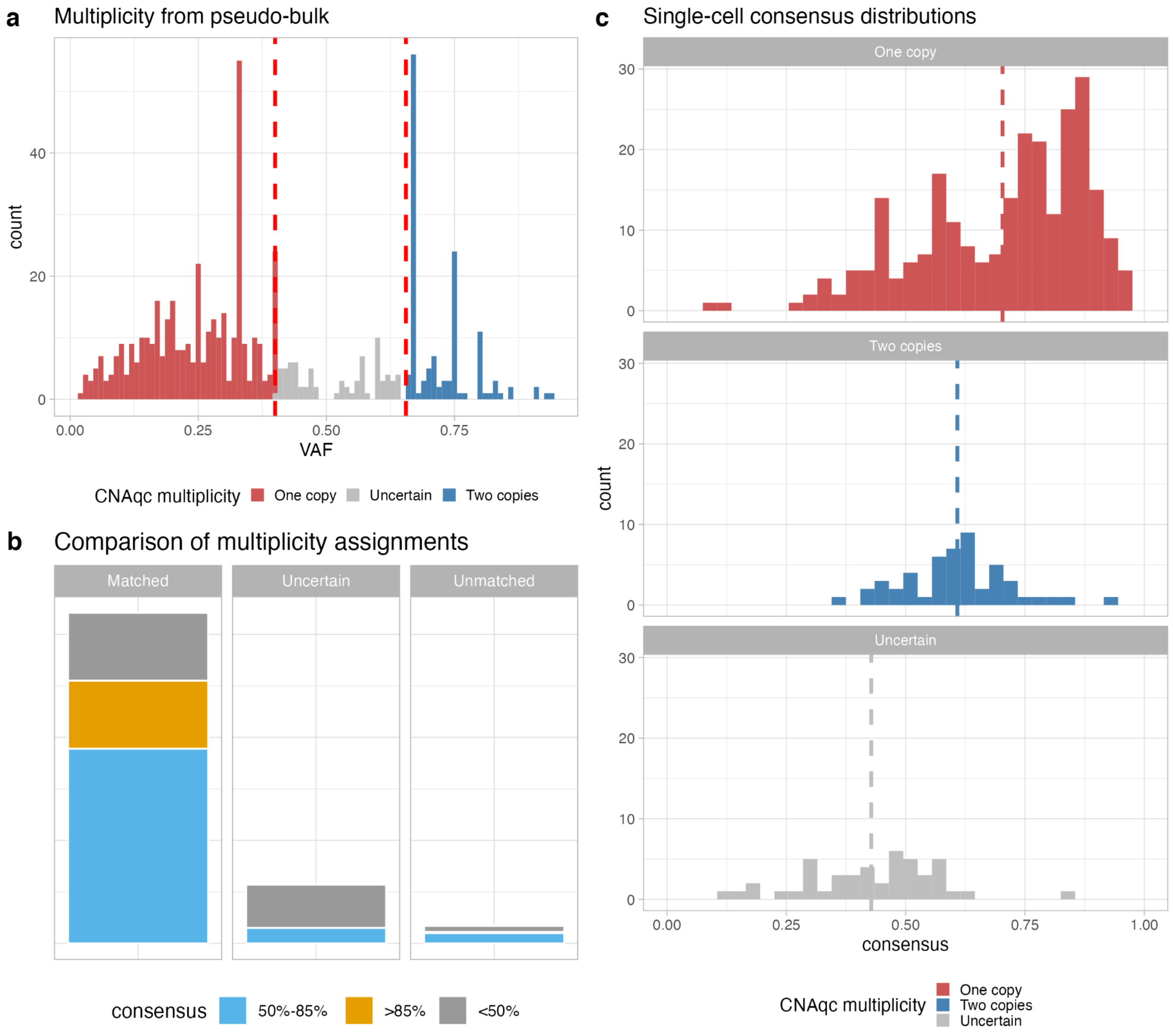


**Supplementary Figure S15. a.** CNAqc multiplicity assignments from pseudo-bulk VAFs of mutations on 2:1 segments against consensus-based assignments from single cells. Test carried out with cells from cluster A from Supplementary Figure S7. Single-cell multiplicities are set to $m = 1$ when the majority of single cells have VAFs closer to 0.33 rather than to 0.66, and are assigned $m = 2$ otherwise. Here CNAqc identifies mutations for which VAFs are uncertain to phase (central cluster). **b.** Most CNAqc multiplicity assignments from pseudo-bulk match the assignments based on consensus of single-cells, especially when the consensus is >85% (high); a small part are unmatched but have intermediate consensus <85% (but above 50%); most mutations with <50% consensus among single cells are flagged as uncertain to phase by CNAqc as expected. **c.** Mutations that CNAqc phases as either in one or two copies have large consensus (mean 70% for $m = 1$ and 61% for 2), while the ones flagged as uncertain to phase have low consensus (mean 43%) among single cells.

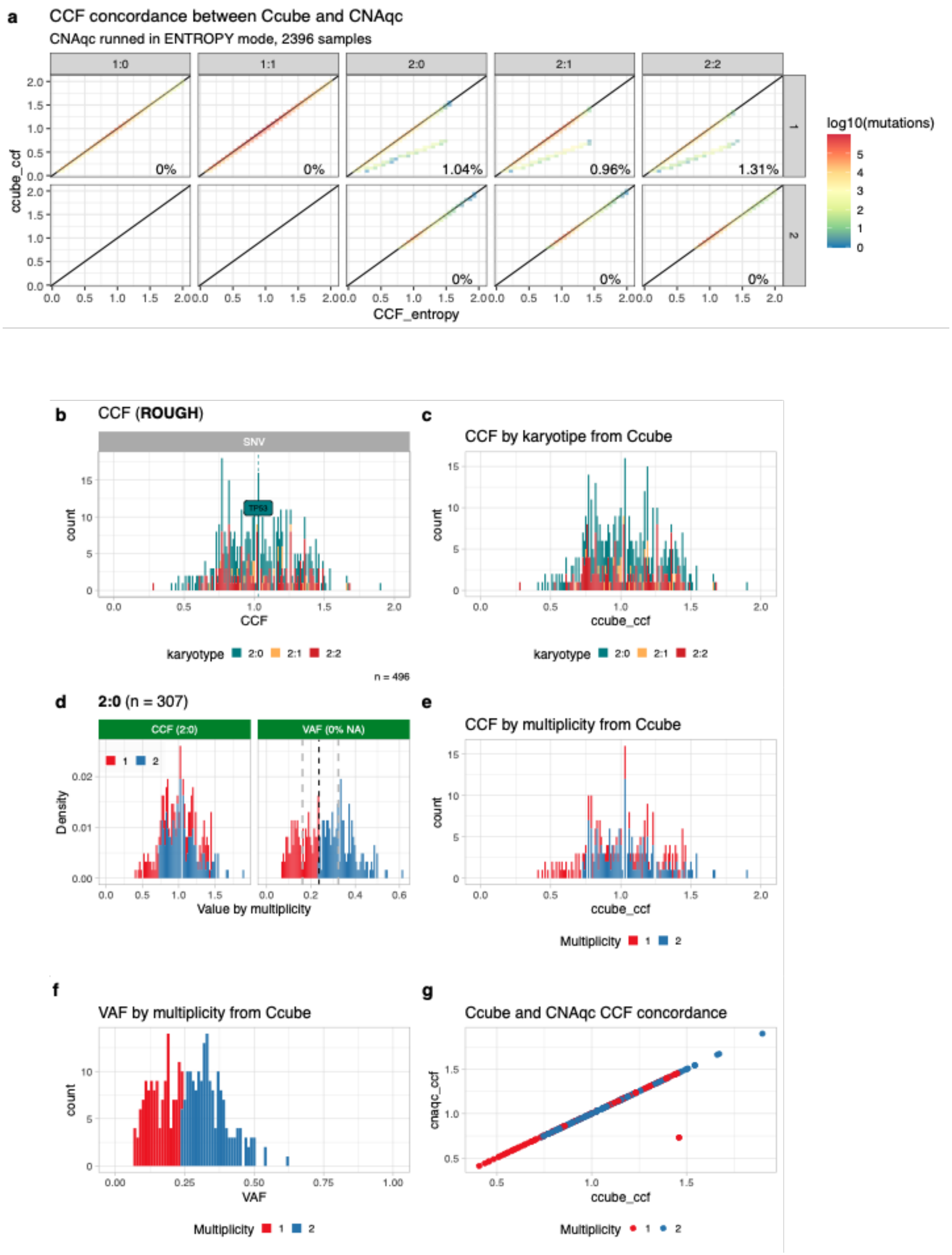
a
CCF concordance between Ccube and CNAqc
CNAqc runned in ENTROPY mode, 2396 samples
1:0
1:1
2:0
2:1
2:2
0%
0%
1.04%
0.96%
1.31%
0%
0%
0%
ccube_ccf
CCF_entropy
log10(mutations)
b
CCF (ROUGH)
SNV
TP53
count
CCF
karyotype 2:0 2:1 2:2
n = 496
c
CCF by karyotipe from Ccube
count
ccube_ccf
karyotype 2:0 2:1 2:2
d
2:0 (n = 307)
CCF (2:0)
VAF (0% NA)
Density
Value by multiplicity
e
CCF by multiplicity from Ccube
count
ccube_ccf
Multiplicity 1 2
f
VAF by multiplicity from Ccube
count
VAF
Multiplicity 1 2
g
Ccube and CNAqc CCF concordance
cnaqc_ccf
ccube_ccf
Multiplicity 1 2

**Supplementary Figure S16. a.** CCF calculated by CNAqc using the “entropy” method against CCF inferred by Ccube on 2396 samples from the PCAWG cohort. Results are divided by karyotype and mutation multiplicity (taking as a reference the one inferred by CNAqc), mutations off the diagonal are discordant between the two methods. On the bottom left, the percentage of those discordant mutations over the total. **b.** CCF calculated by CNAqc using the “rough” method which assigns the multiplicity by splitting the clonal clusters at VAF level. **c.** CCF inferred by Ccube. **d.** Multiplicity assigned by CNAqc, the dashed black line depicts the splitting point to determine multiplicity. **e.** Multiplicity assigned by Ccube. **f.** VAF split by Ccube for multiplicity assignment. **g.** CCF values between Ccube and CNAqc are in almost perfect agreement (just one sample has different multiplicity). Point colour is based on the multiplicity estimated by Ccube.

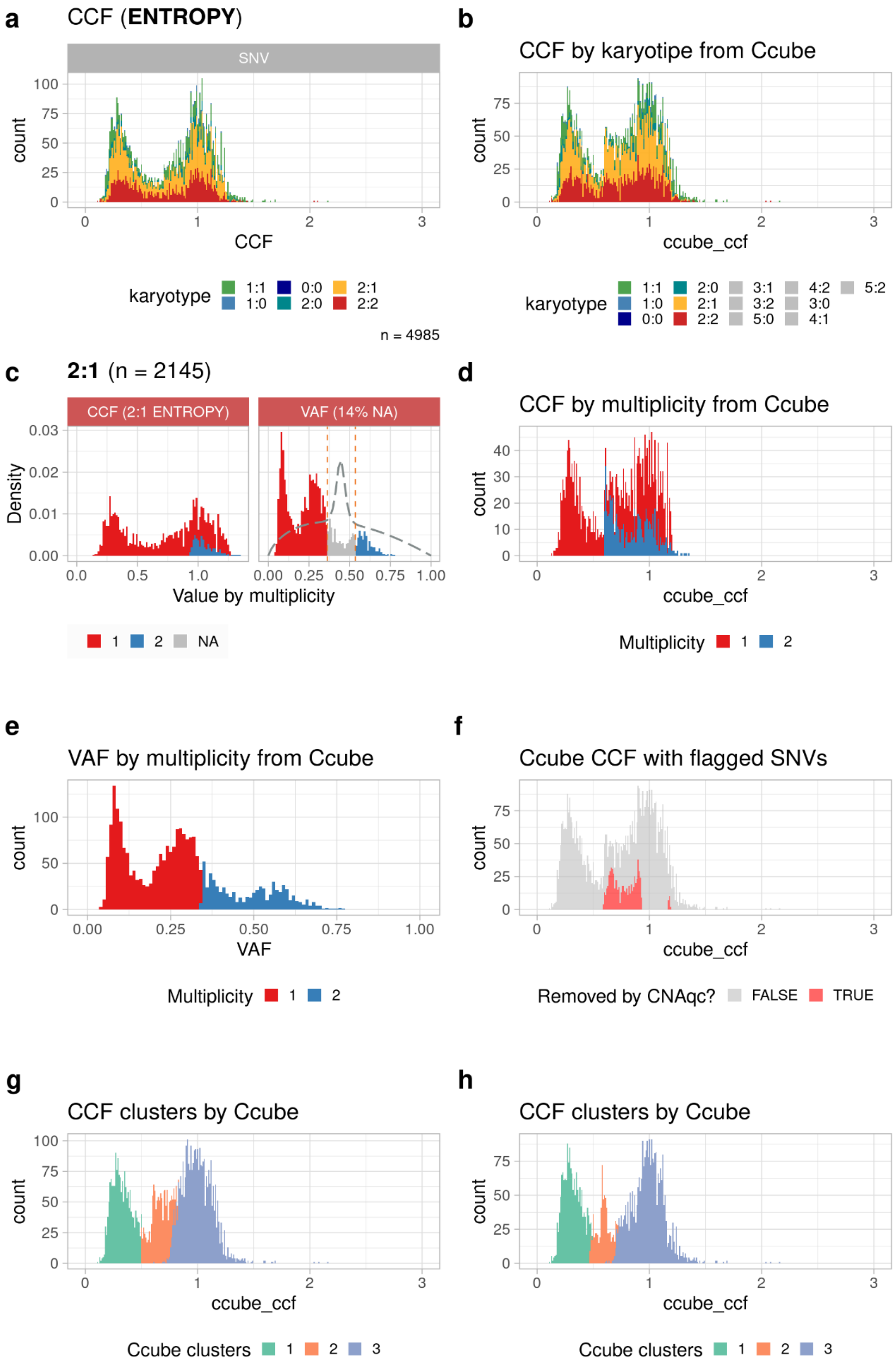


**Supplementary Figure S17. a.** CCF calculated by CNAqc using the “entropy” method which discards mutations with high multiplicity uncertainty. **b.** CCF inferred by Ccube. The main difference between this profile and the one inferred by CNAqc is a bump around CCF 0.6 **c.** Multiplicity assigned by CNAqc, in grey the mutations with non-estimable multiplicity. **d.** Multiplicity assigned by Ccube, it can be noted how Ccube always assigns a definite multiplicity value to each mutation. **e.** VAF split by Ccube for multiplicity assignment. **f.** High-entropy mutations discarded by CNAqc in the Ccube CCF profile. We clearly see the extra spike in CCF which could confound subclonal deconvolution, splitting the clonal cluster in multiple clones. **g.** Ccube recognizes the spurious peak as a subclonal cluster, as it is not able to accommodate for the overdispersion derived by the errors in multiplicity assignments with just one cluster. **h.** Even after removing the mutations with high entropy from the dataset and rerunning Ccube, we can still see a peak caused by some mutations wrongly assigned to multiplicity 2. This is consistent with the choice of CNAqc to FAIL the available CCFs for this karyotype.

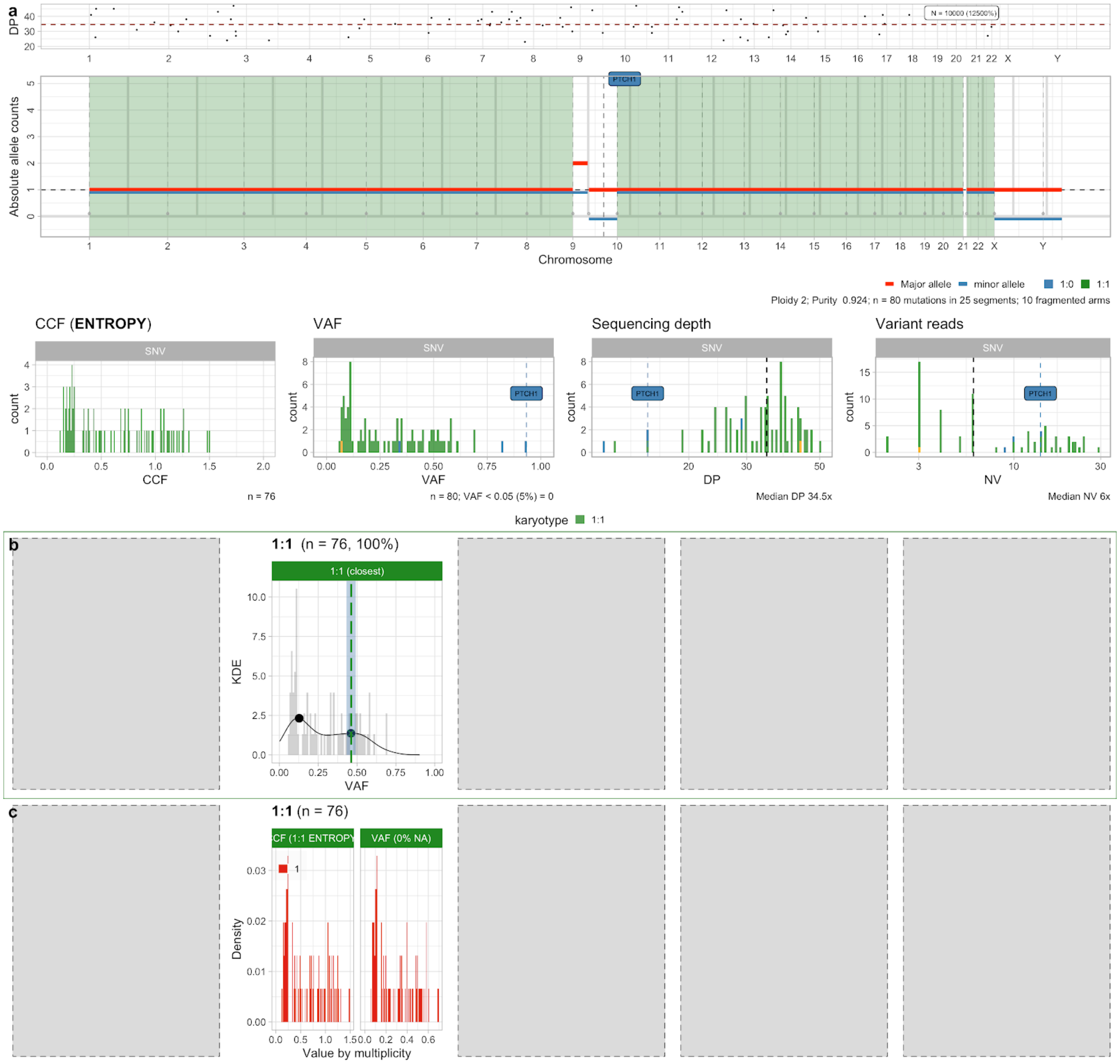


**Supplementary Figure S18.** Example PCAWG medulloblastoma sample with low-mutational burden, which passes data QC with CNAqc. **a.** Data for the sample (genome-wide CNA segments, CCF and read counts distribution). Note that this sample has only 76 SNVs in diploid tumour regions, like we observe in whole-exome assays. **b,c.** Peak analysis and CCF computation for diploid SNVs.

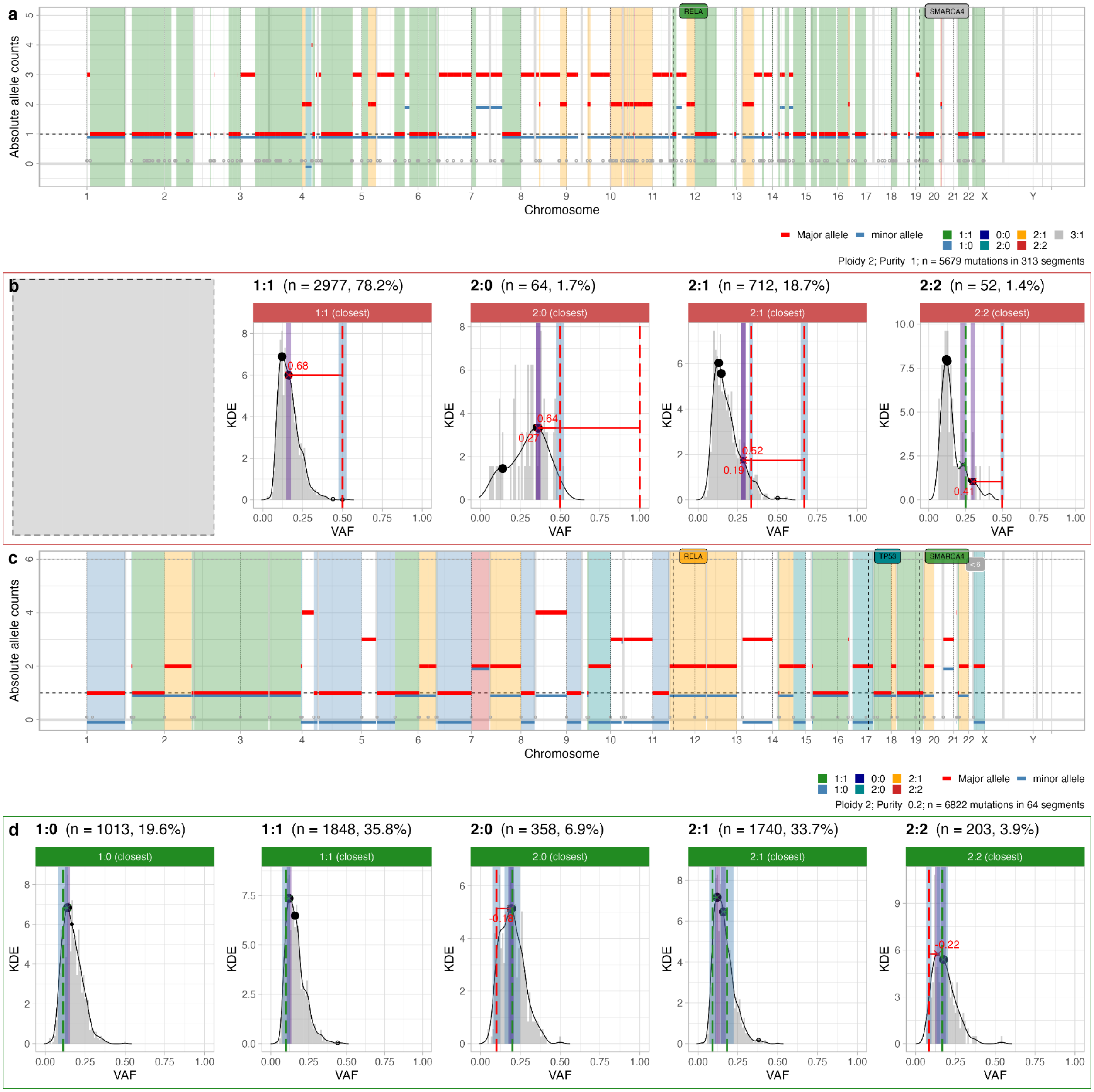


**Supplementary Figure S19.** Example PCAWG sample with 100% purity, against an alternative solution with 20% purity. **a.** The PCAWG copy number profile at purity 100% is diploid on average, with subclonal segments on chromosomes 2 and 17 (not used by CNAqc). **b.** Peak analysis with CNAqc for the PCAWG solution in panel (a). For both diploid and non-diploid regions, all peaks are mismatched (FAIL). Note that this solution has around 80% of its SNVs in diploid tumour regions, where the VAF peaks at ~10%, possibly suggesting a purity well below 100%. **c.** Rerun of this sample with the Sequenza-CNAqc pipeline (Supplementary Figure S17) assigns a diploid profile with purity 20%, obtained from one of the alternative solutions of Sequenza. **d.** VAF peaks are matched for both diploid and non-diploid regions, resulting in PASS status for each karyotype, and for the overall quality control procedure.

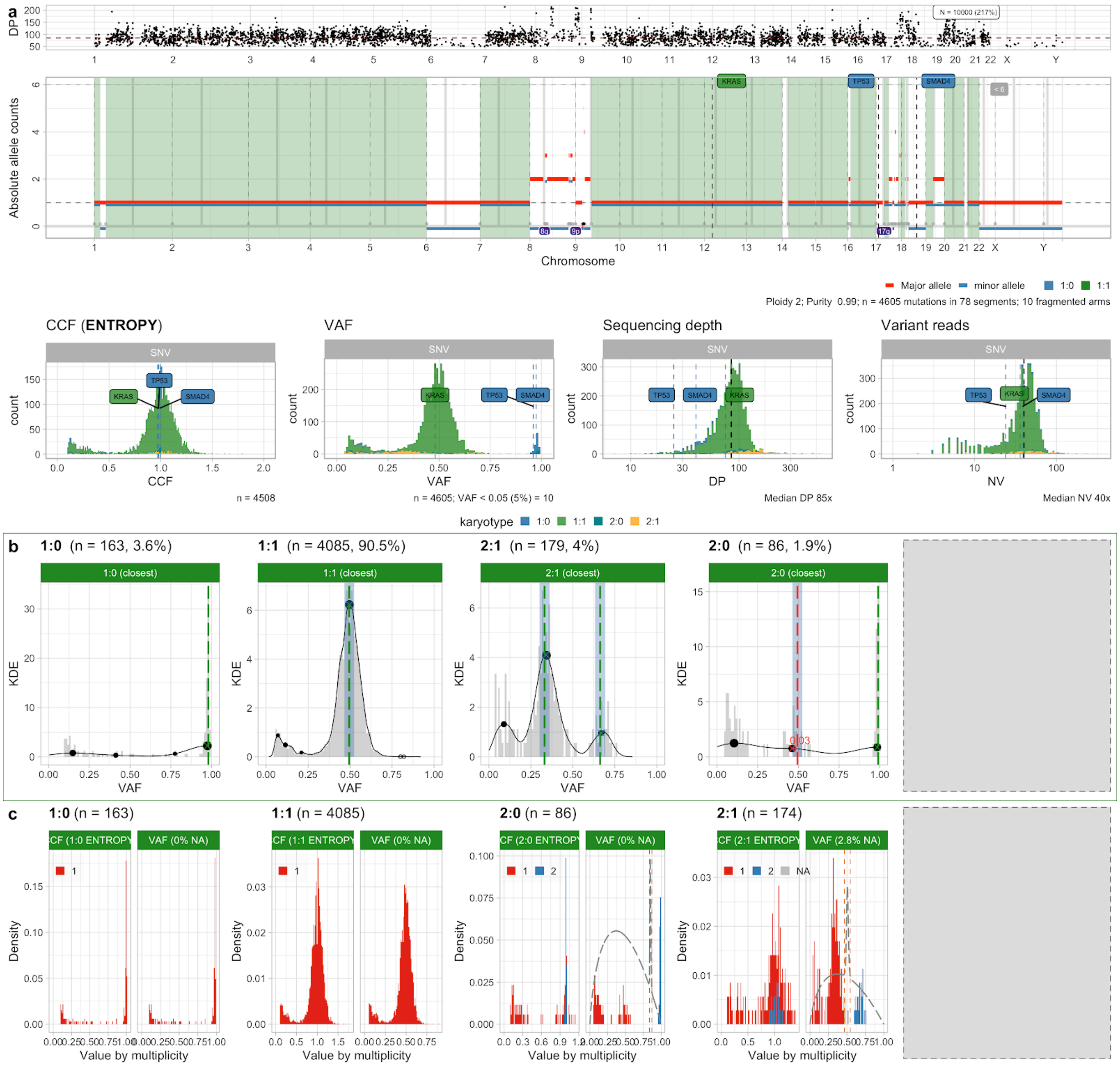


**Supplementary Figure S20.** Example PCAWG pancreatic adenocarcinoma with 99% purity (and 3 possible driver SNVs, 2 of them involving tumour suppressor genes in LOH regions). **a.** Data for the sample (genome-wide CNA segments, CCF and read counts distribution). **b.** This sample has 90% of its SNVs in diploid tumour regions, and the others in a variety of distinct CNA segments. From a peak analysis point of view, all the calls are validated. **c.** CCF values for this sample are also good.

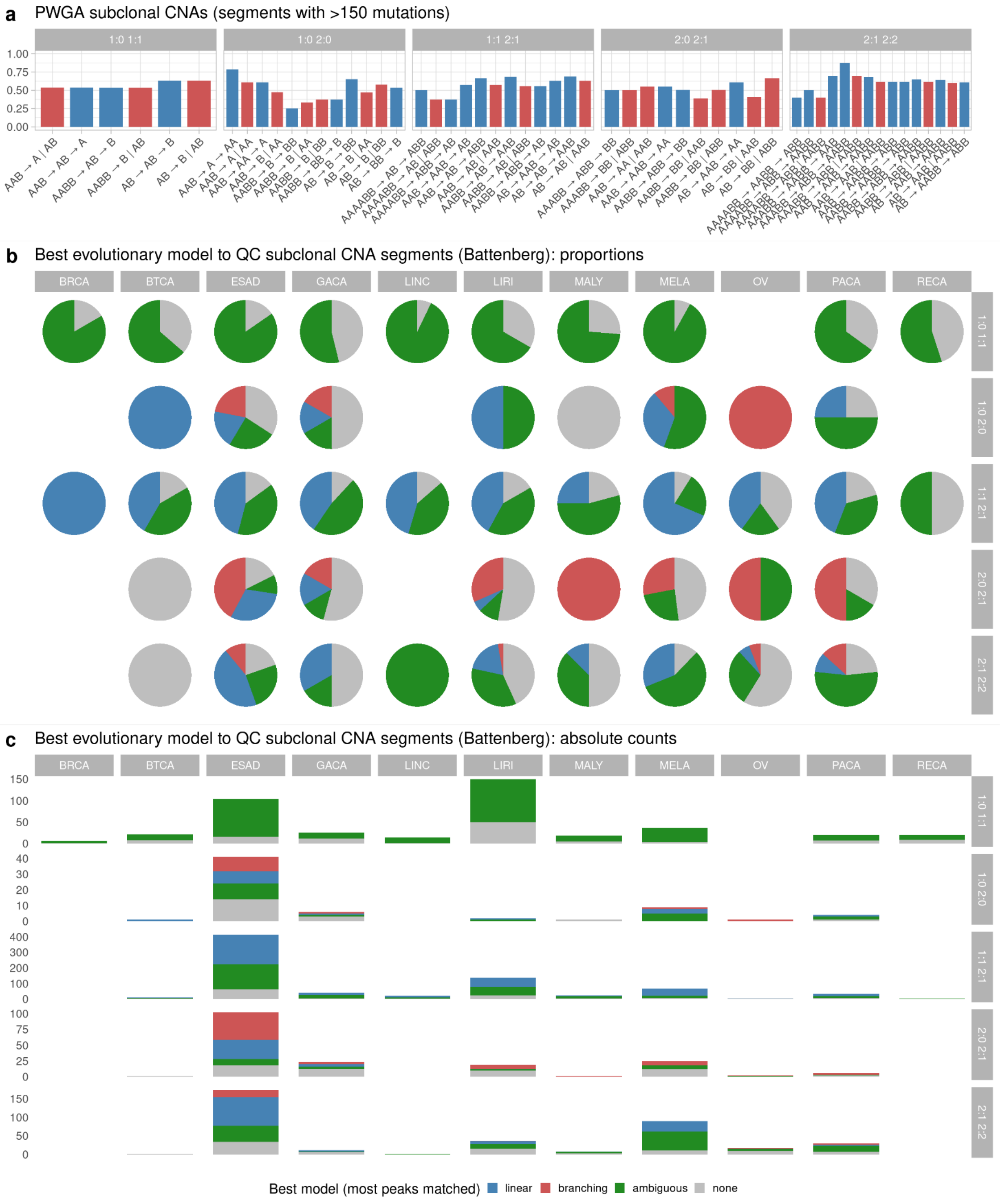


**Supplementary Figure 21. a.** Percentage of matched peaks for each possible model computed for different karyotype combinations. The colour of the bar reflects whether the referred model is branching or linear (red and blue respectively). **b.** Different prevalences of best models across different tumour types and karyotype combinations. Best models are determined according to the percentage of matched peaks. When the number of matched peaks is the same for models belonging to different classes the imputation is uncertain (ambiguous), those cases are coloured in green in the graph. Finally, when no model could be fitted to the data (<50% of the expected peaks matched), the 'none' value is assigned, those cases are coloured in grey. **c.** Absolute counts akin to the pie charts in panel (b).

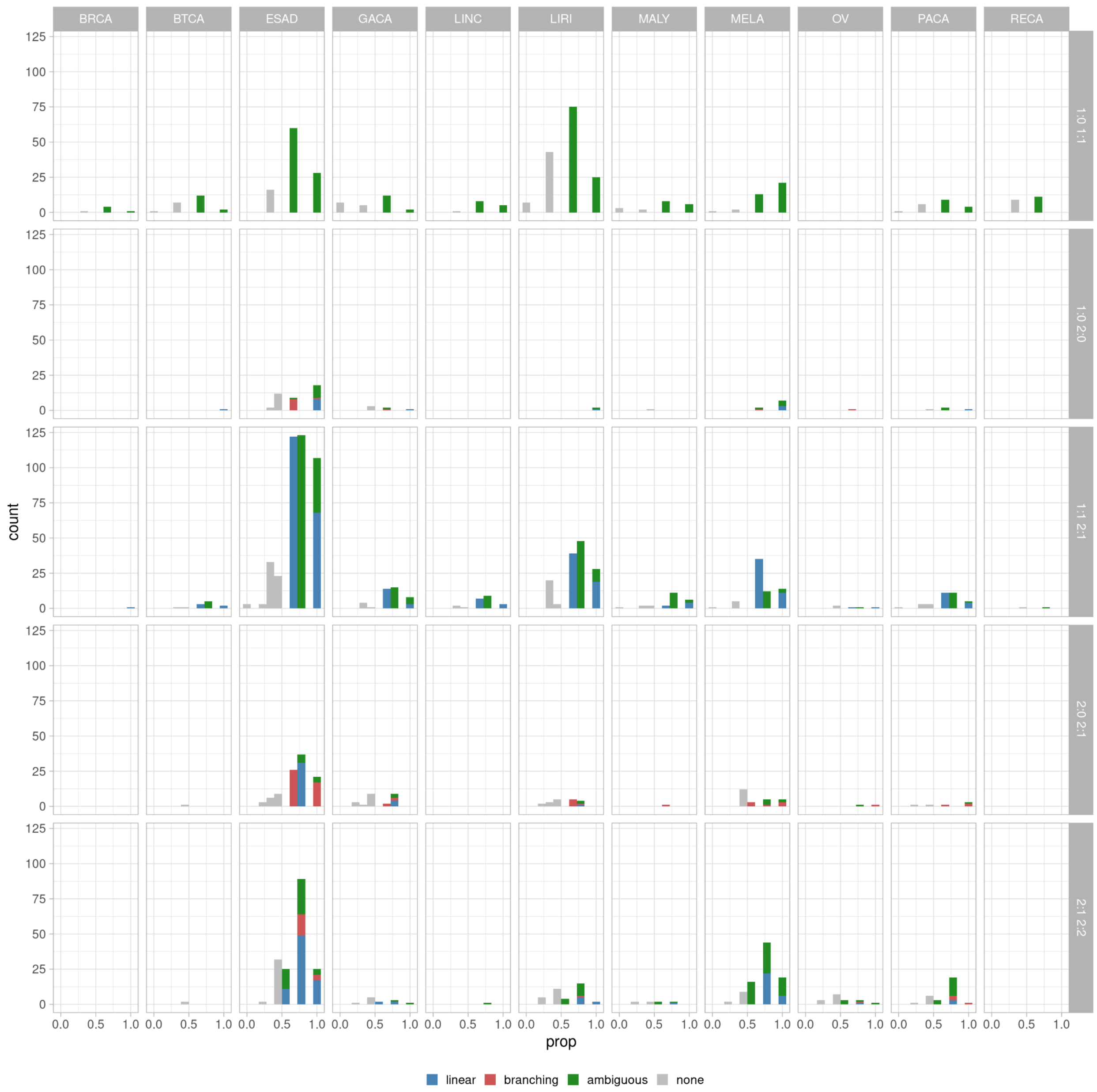


**Supplementary Figure 22.** Number of models assigned to segments and peak matching percentages. Counts are divided according to tumour type and subclonal karyotypes. Linear models explain the data better in most cases with the exception of 2:0-2:1 karyotype where the branching model prevails, and the 1:0-1:1 karyotype where it is impossible to distinguish between models. Interestingly, similar patterns are repeated across tumours, both in terms of relative abundance of the different models and of different karyotypes. This property may suggest that the amplification mechanisms are shared across tumours, although some tumour types display a higher tendency to acquire CNAs.

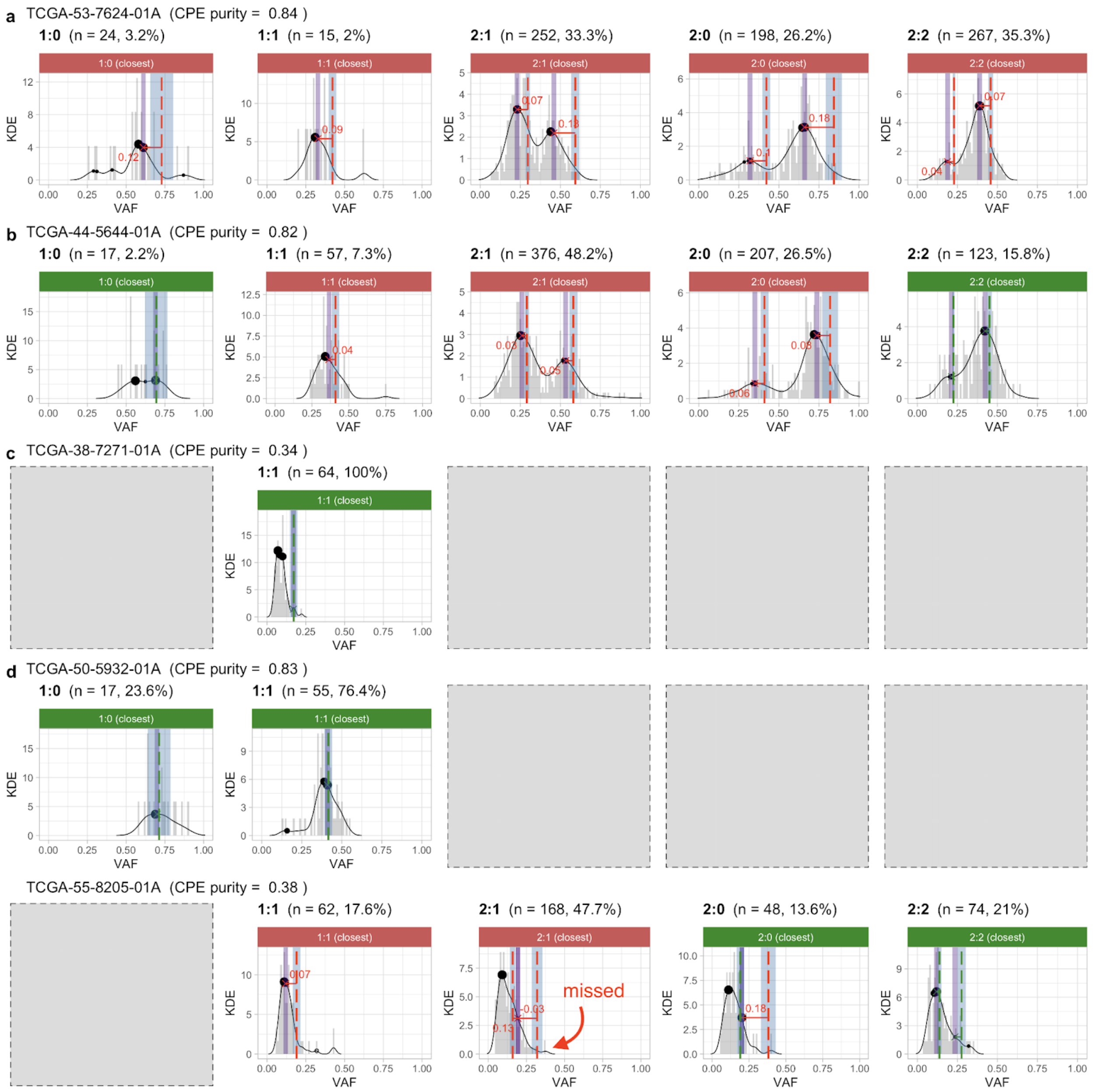


**Supplementary Figure S23. a-d.** CNAqc quality control via peak detection on TCGA whole-exome sequencing data of 5 lung adenocarcinomas (LUAD) with different purity values, selected from a cohort of 48 cases available online.

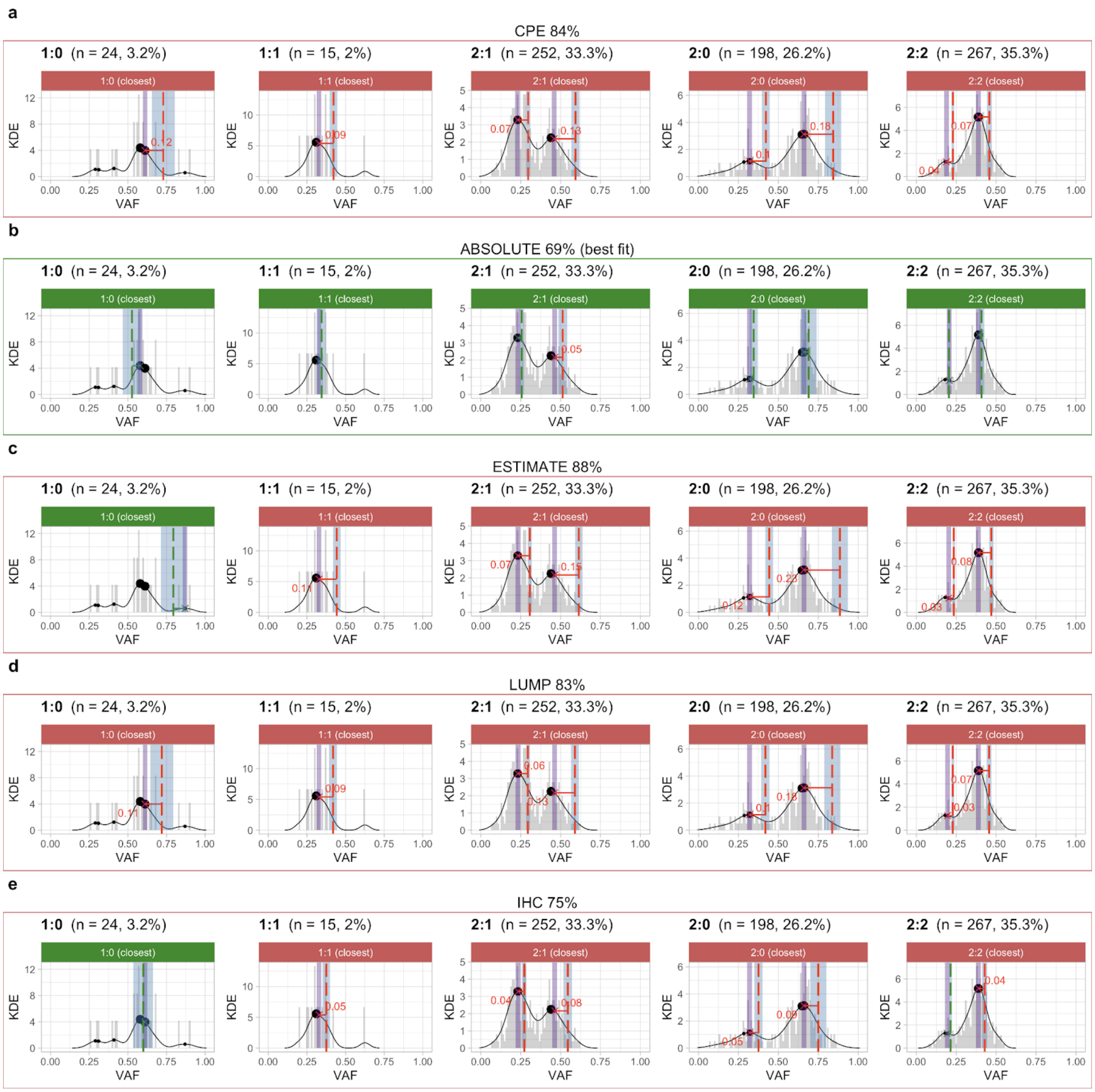


**Supplementary Figure S24. a-e.** CNAqc quality control via peak detection for LUAD sample TCGA-53-7624-01A - panel (a) of Supplementary Figure S25 - using purity estimates from CPE (consensus), ABSOLUTE, ESTIMATE, IHC and LUMP. CNAqc determines that, among all callers, only ABSOLUTE detected the true tumour ploidy (69%).

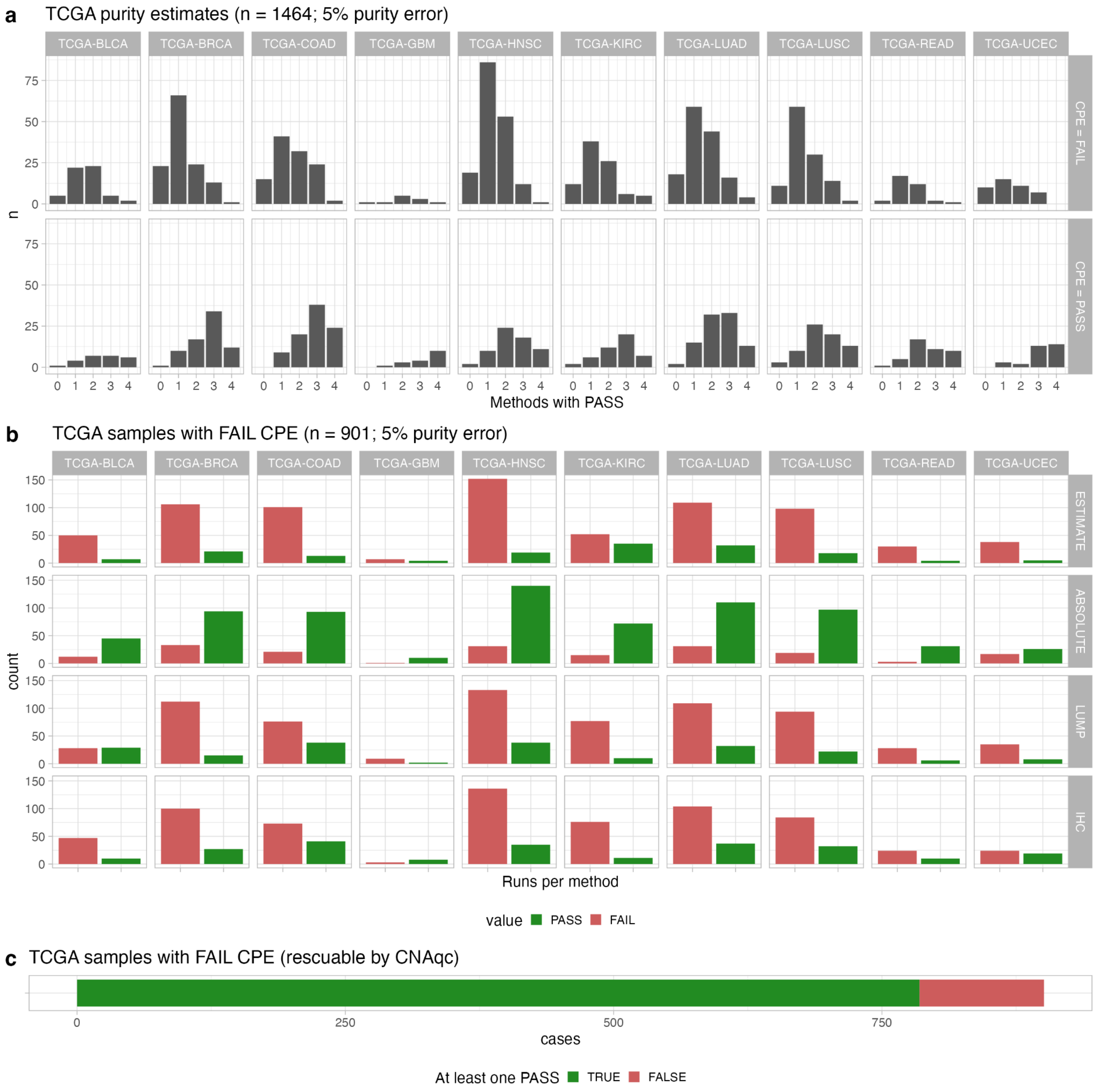


**Supplementary Figure S25. a.** CNAqc quality control via peak detection for 1464 TCGA samples from 10 distinct tumour types with suitable data for our tool. Plots are split by QC status (maximum tolerated purity error 5%) based on TCGA consensus purity (CPE purity score). The bars report the number of methods whose purity is passed by CNAqc. We used purity estimates from ABSOLUTE, ESTIMATE, IHC and LUMP as in Supplementary Figure S26. **b.** For 901 cases where the CPE purity is failed by CNAqc, we report the number of cases where each method is instead passed. Note for instance that ABSOLUTE often provides a purity estimate that would pass the sample according to CNAqc, similarly to the case shown in Supplementary Figure S26. **c.** 785 out of 901 cases (~88%) would be rescued if we selected at least one of the purity estimates that passes CNAqc analysis, avoiding instead of using the CPE consensus purity which contains an error larger than 5%.

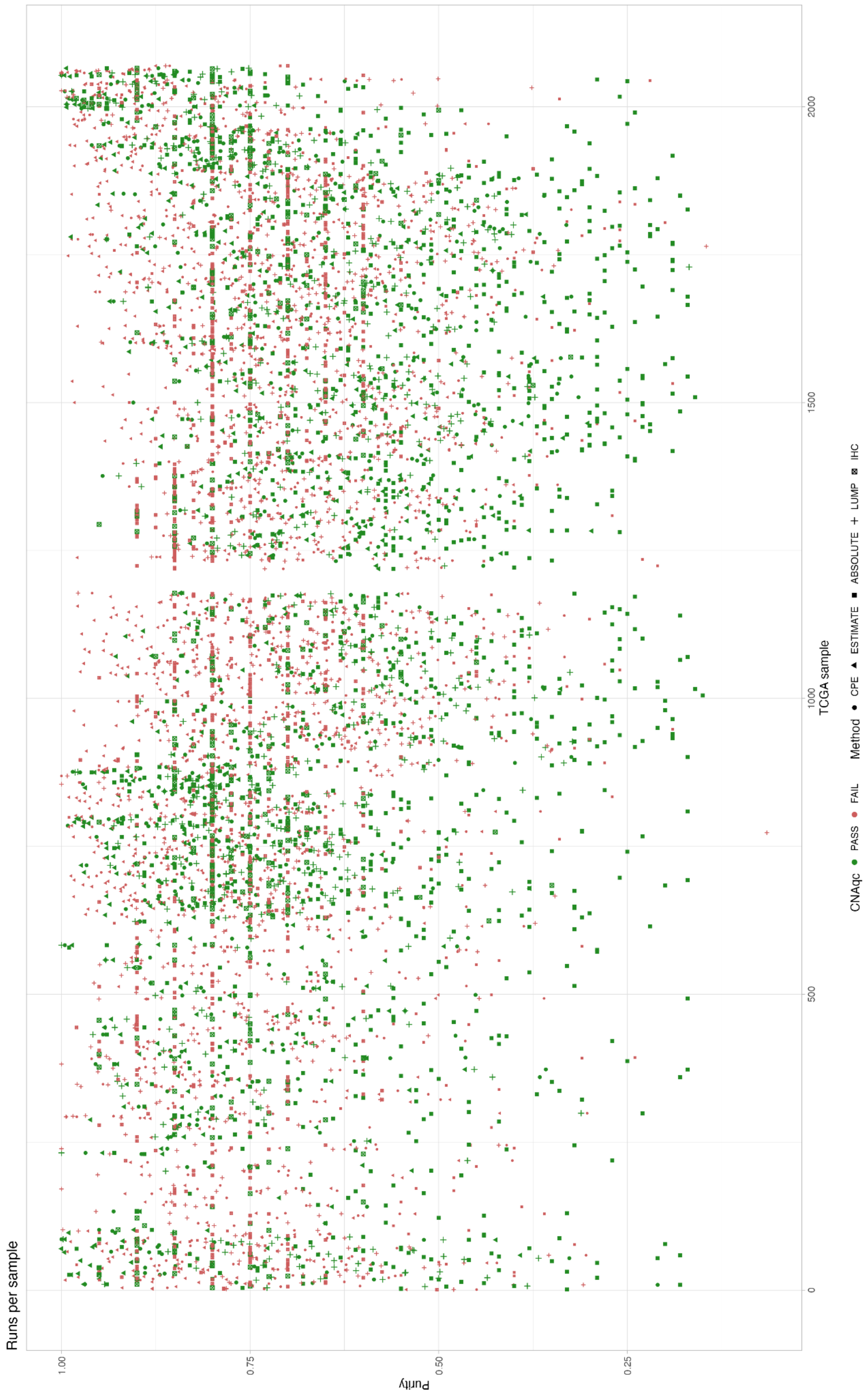


**Supplementary Figure S26.** TCGA runs from Supplementary Figure S27. On the x-axis samples are sorted by sample id; on the y-axis the purity is reported for each method. Each method has a different point shape; the colour reflects CNAqc pass or fail status. Therefore, cases where green (pass) points are at lower purity compared to red ones, represent TCGA samples where some methods over-estimate purity; in the opposite case we observe insead under-estimation.

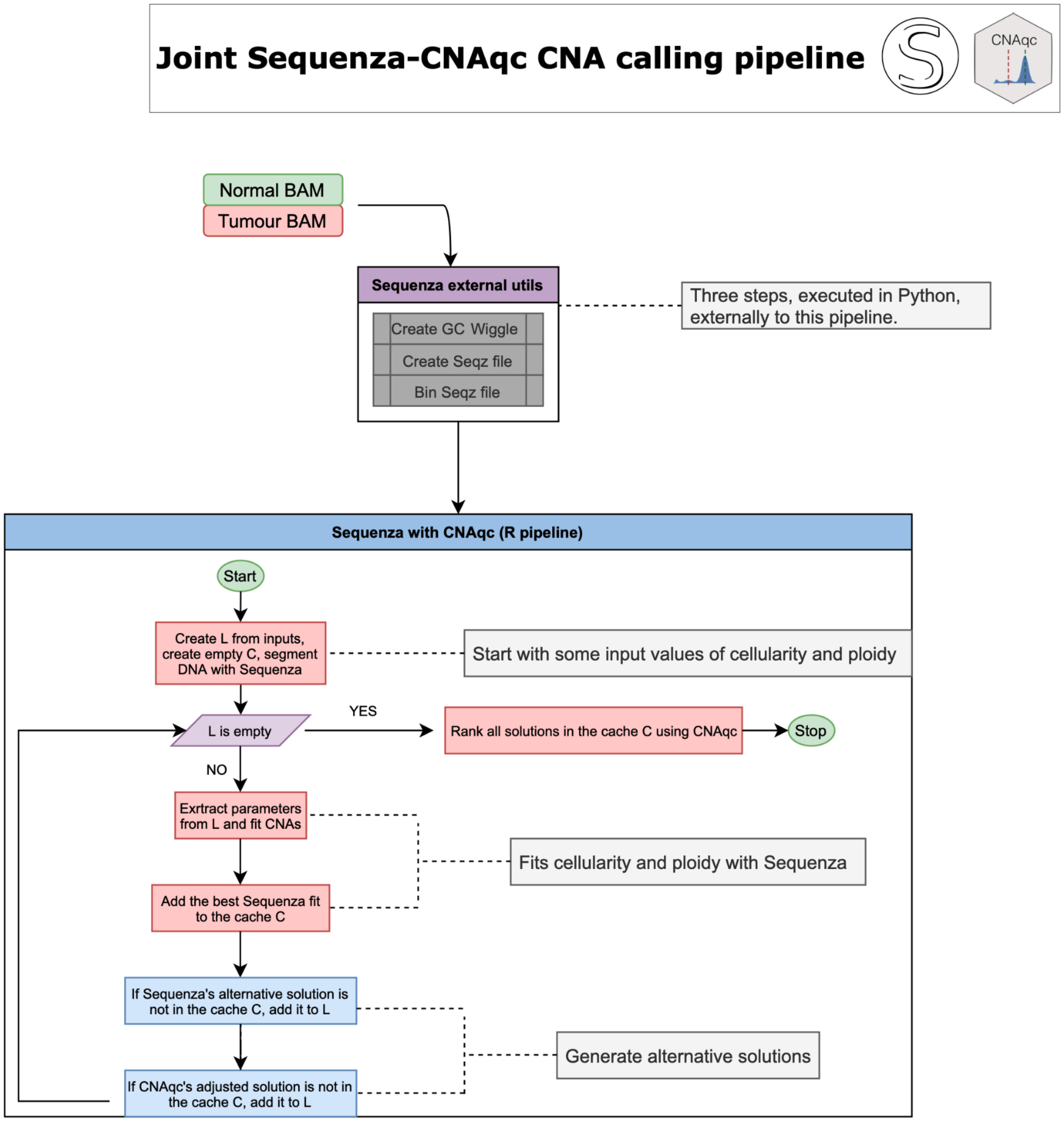


**Supplementary Figure S27.** Flowchart of the joint Sequenza - CNAqc pipeline for CNA calling. Sequenza external utils are first used to generate a binned seqz file. The pipeline then performs segmentation (via Sequenza), and iteratively optimises cellularity (i.e., tumour purity) and ploidy estimation, together with allele-specific CNA segments, using a list of solutions to inspect L, and a cache of already examined solutions C. Starting from the input ranges for cellularity and ploidy, at each step Sequenza is run to fit the data. Alternative solutions are generated via Sequenza and via CNAqc: if the proposed solutions have not been already analysed (i.e., they are not in C), then they are added to the list of solutions to examine (i.e., they are instead in L). At the end, when L is empty, all the cached solutions are examined with CNAqc and ranked to determine the best fit a FAIL/PASS status by peak detection.

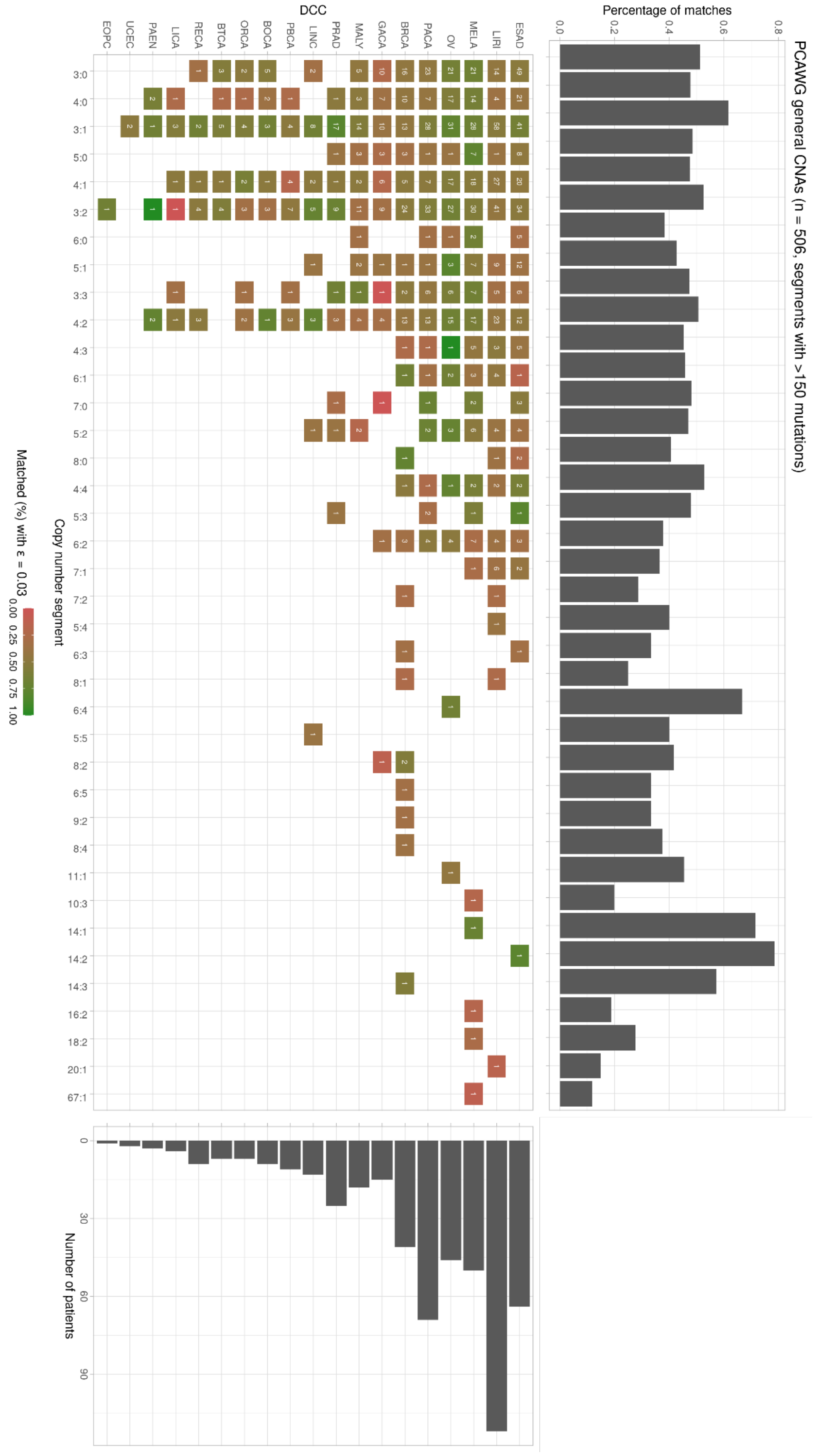


**Supplementary Figure S28.** 570 PCAWG samples with complex CNA segments. For this type of peak analysis, the tool reports if any of a set of expected peaks is detected in the data. Since segments with a low mutational burden carry little peak information and are prone to noise, we show only segments with >150 somatic mutations. The marginal right panel shows the percentage of matched peaks for each rare karyotype found in the PCAWG cohort. In the central panel, segments are divided according to their karyotype and tumour type: each square reports the number of segments and its colour encodes the percentage of matched peaks. The marginal bottom panel shows the number of patients for each tumour type.

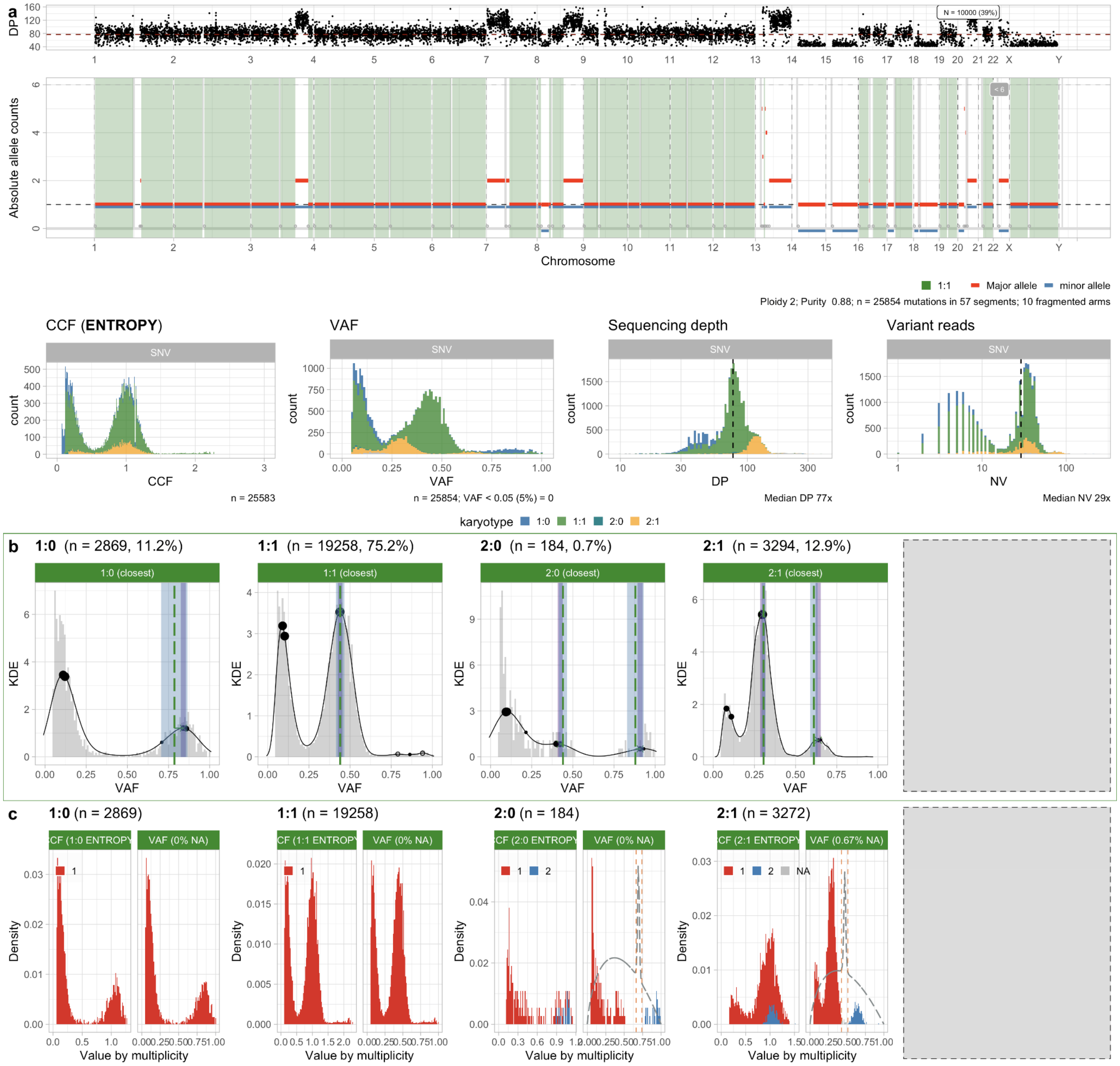

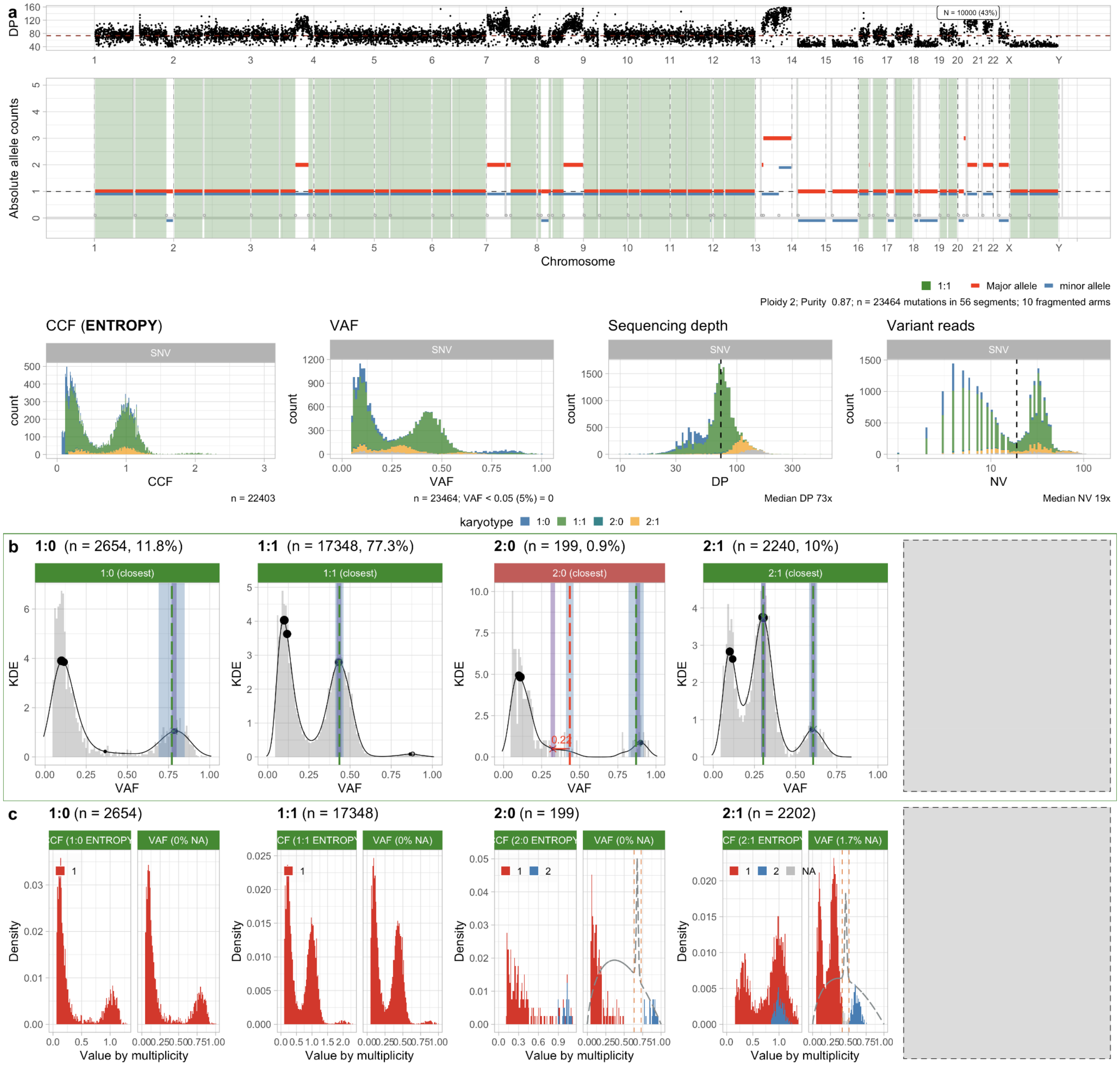
a
DP
N = 10000 (43%)
Absolute allele counts
Chromosome
1:1
Major allele
minor allele
Ploidy 2; Purity 0.87; n = 23464 mutations in 56 segments; 10 fragmented arms
CCF (ENTROPY)
VAF
Sequencing depth
Variant reads
SNV
CCF
n = 22403
n = 23464; VAF < 0.05 (5%) = 0
DP
Median DP 73x
NV
Median NV 19x
karyotype 1:0 1:1 2:0 2:1
b
1:0 (n = 2654, 11.8%)
1:1 (n = 17348, 77.3%)
2:0 (n = 199, 0.9%)
2:1 (n = 2240, 10%)
1:0 (closest)
1:1 (closest)
2:0 (closest)
2:1 (closest)
KDE
0.22
c
1:0 (n = 2654)
1:1 (n = 17348)
2:0 (n = 199)
2:1 (n = 2202)
VAF (0% NA)
VAF (1.7% NA)
Density
Value by multiplicity

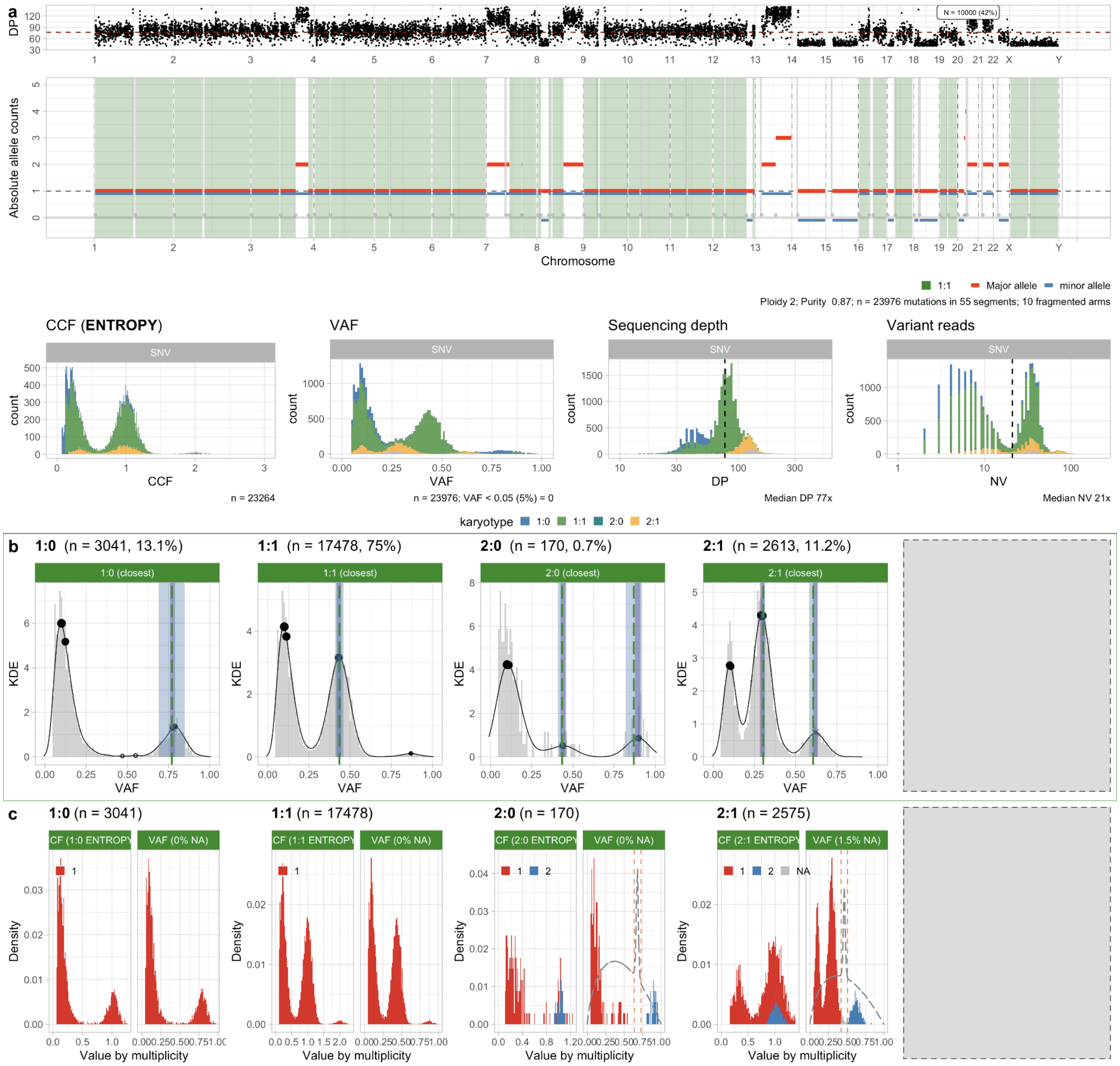
a
DP
Absolute allele counts
Chromosome
1:1 Major allele minor allele
Ploidy 2; Purity 0.87; n = 23976 mutations in 55 segments; 10 fragmented arms
CCF (ENTROPY)
VAF
Sequencing depth
Variant reads
n = 23264
n = 23976; VAF < 0.05 (5%) = 0
Median DP 77x
Median NV 21x
karyotype 1:0 1:1 2:0 2:1
b
1:0 (n = 3041, 13.1%)
1:1 (n = 17478, 75%)
2:0 (n = 170, 0.7%)
2:1 (n = 2613, 11.2%)
c
1:0 (n = 3041)
1:1 (n = 17478)
2:0 (n = 170)
2:1 (n = 2575)
Value by multiplicity

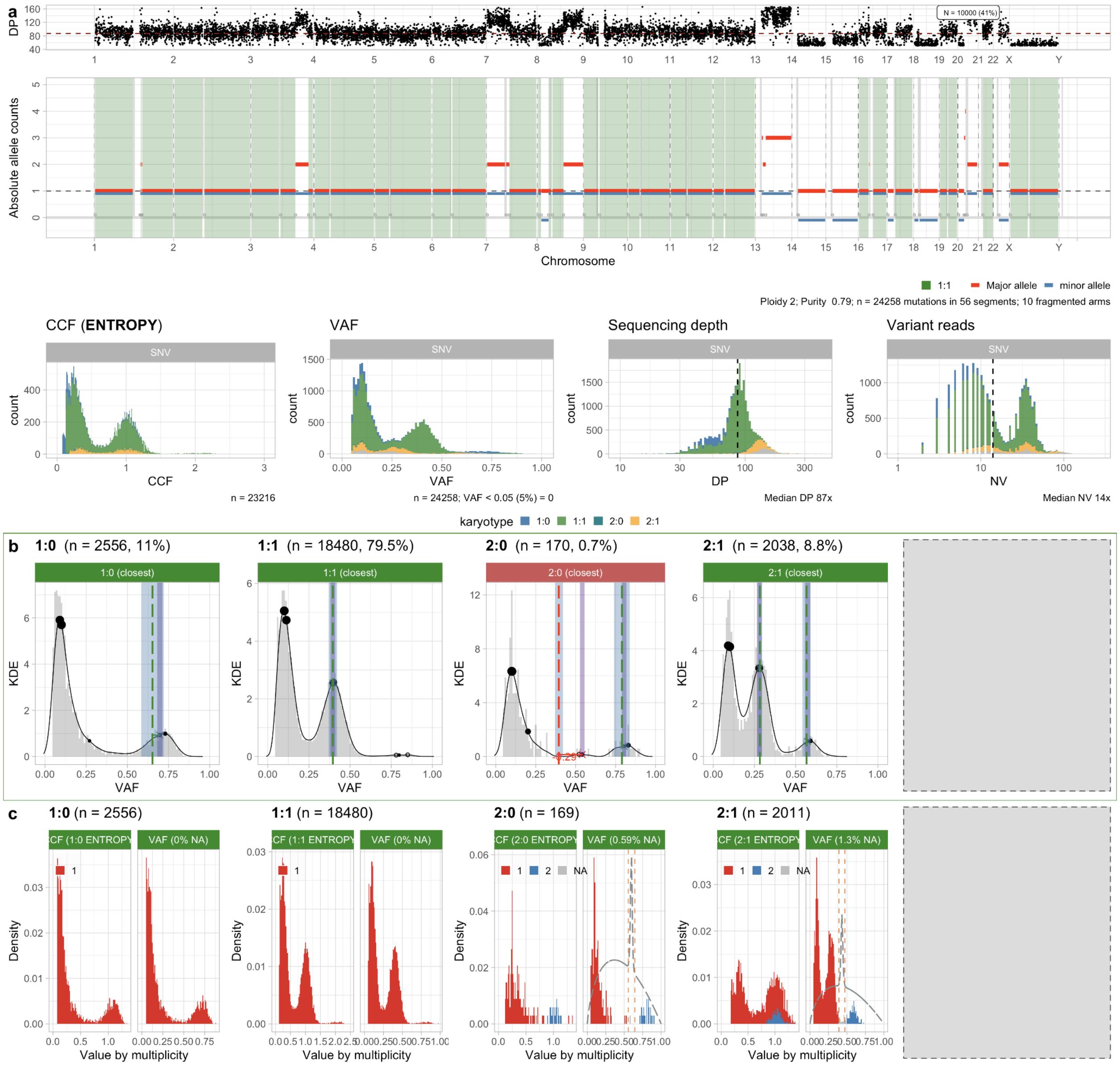

**Supplementary Figure S29 (multiple pages).** Colorectal multi-region samples (one per page): Set7_55, Set7_57, Set7_59 and Set7_62 for patient Set7. **a.** Allele-specific CNAs, and read data distribution (bottom row). **b,c.** Peak analysis and CCF computation for the sample.

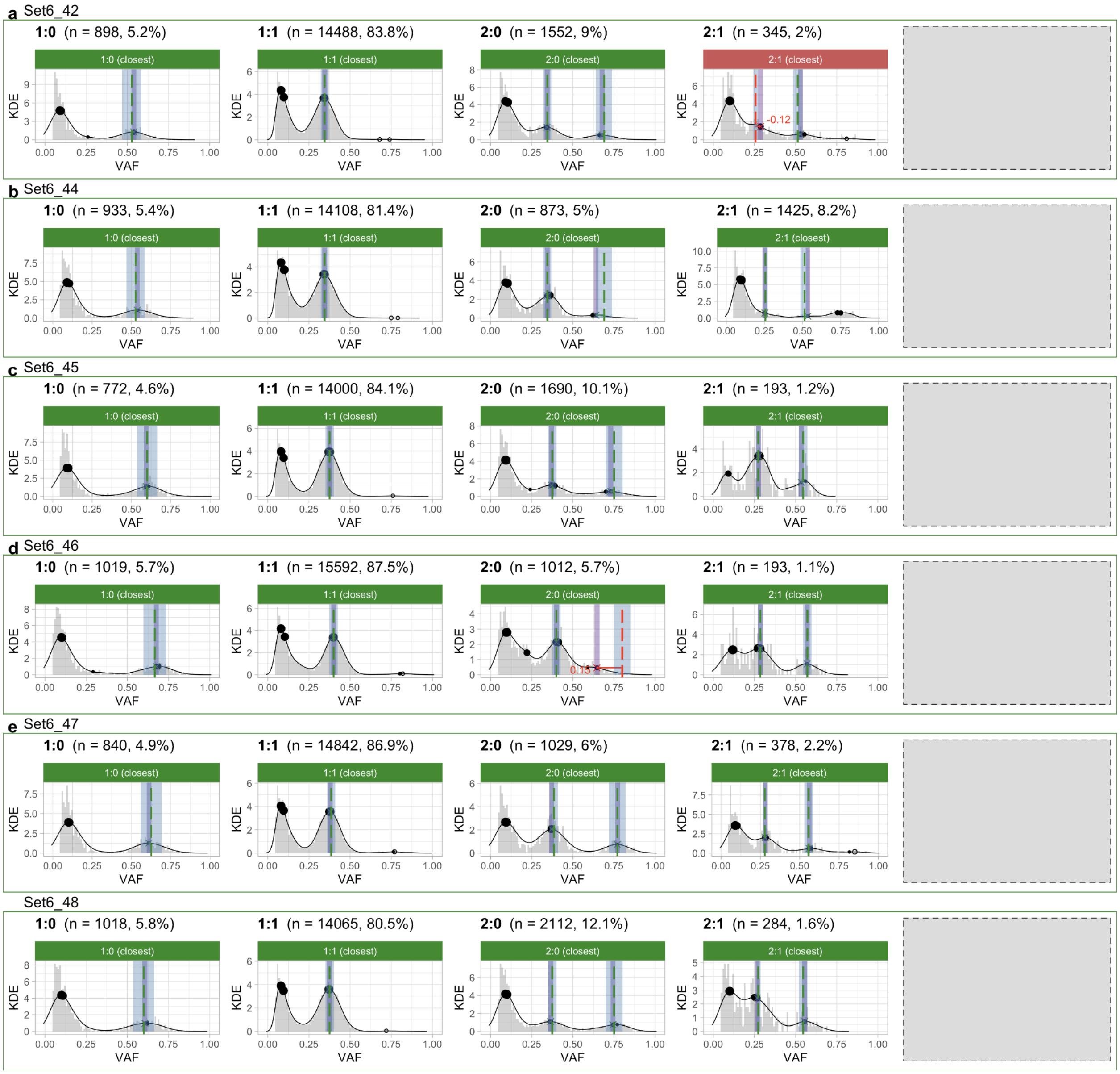


**Supplementary Figure S30. a,b,c,d,e.** Peak detection quality control with CNAqc, run with default parameters on colorectal multi-region samples available for patient Set_6. All calls are passed (surrounding green rectangles).
